# Metabolic Maturation Unveils Left Ventricular Identity in WNT ON/OFF Human Pluripotent Stem Cell-Derived Cardiomyocytes

**DOI:** 10.64898/2025.12.30.696943

**Authors:** Joaquín Smucler, Julia María Halek, Denisse Saulnier, Sheila Lucia Castañeda, Agustina Scarafía, Guadalupe Amín, Alejandra Guberman, Gustavo Sevlever, Santiago Miriuka, Lucía Natalia Moro, Ariel Waisman

## Abstract

Deriving high-purity mature left ventricular (LV) cardiomyocytes (CMs) from human pluripotent stem cells (hPSCs) is a priority for cardiovascular research and for future therapeutic applications. Small-molecule WNT modulation (WNT ON/OFF) is currently the predominant differentiation method; however, a critical discordance exists regarding its cardiac subtype outcome. While lineage tracing suggests a First Heart Field (FHF) bias, phenotypic characterizations report significant heterogeneity regarding definitive ventricular markers, leading to controversy about the cardiac subtypes generated by this method. Here, we demonstrate that this apparent heterogeneity is a result of CM immaturity. Using single-cell protein analysis, we first show that WNT ON/OFF generates an NKX2.5+ progenitor pool that robustly co-expresses HAND1, confirming uniform FHF specification regardless of differentiation efficiency. We then demonstrate that the CM population negative for the ventricular marker MYL2 observed at early differentiation timepoints mostly represents immature LV cardiomyocytes that have not yet acquired their definitive phenotype. By implementing a targeted metabolic maturation regime, we unlocked this identity, achieving 95% MYL2+/HAND1+/TBX5+ LV CMs by day 38, substantially earlier and with higher chamber-specific purity than previously reported. This phenotypic resolution was accompanied by advanced structural maturation, including sarcomeric protein isoform switching, multinucleation, and notably, the assembly of polarized XIRP2+ intercalated discs, a hallmark of postnatal CM maturation not previously described in 2D differentiations. Validated across three independent hPSC lines, these findings provide the field with a rapid, high-fidelity platform for generating pure mature LV cardiomyocytes for disease modeling and therapeutic research.

## INTRODUCTION

Cardiac differentiation of human pluripotent stem cells (hPSCs) provides an invaluable platform for modeling human heart development and offers an essentially unlimited source of cardiomyocytes (CMs) for disease modeling, drug screening, and potential cell-based therapies (Bois et al., 2025). The adult mammalian heart comprises multiple specialized CM subtypes, including left and right ventricular and atrial CMs, nodal cells, and derivatives of outflow tract (OFT) cells, each arising during embryonic development from two spatially and temporally distinct progenitor pools: the first heart field (FHF) and the second heart field (SHF) (Yang et al., 2022). The FHF, characterized by expression of NKX2.5, TBX5, and HAND1, predominantly generates the left ventricle, while the SHF, marked by NKX2.5 and ISL1 expression, contributes to the right ventricle, OFT, and atrial regions (Brade et al., 2013; Ivanovitch et al., 2021; Yang et al., 2022). Precisely directing hPSC differentiation toward these specific cardiac lineages *in vitro* is essential for creating physiologically relevant cardiac models. Of particular importance are left ventricular cardiomyocytes (LV-CMs), which represent the primary cell type affected in ischemic heart disease and constitute the principal target for regenerative therapies (Jiang et al., 2012)

Several methods exist to induce hPSC-derived CMs (hPSC-CMs) *in vitro*, all recapitulating developmental milestones as cells transition through primitive streak–like mesendoderm, cardiac mesoderm, and cardiac progenitor states before differentiating into beating CMs (Brade et al., 2013). Traditional embryoid body (EB) approaches using cytokines such as BMP4, Activin A, and bFGF (Burridge et al., 2011; Yang et al., 2008) may better recapitulate certain aspects of embryonic development but are costly and exhibit variable reproducibility. In contrast, monolayer-based protocols utilizing small-molecule WNT pathway modulators, particularly CHIR99021, offer significant advantages in cost-effectiveness, scalability, and reproducibility and have become the predominant approach for cardiac differentiation (Gonzalez et al., 2011; Lian et al., 2012; Lyra-Leite et al., 2022).

While EB methods have been recently adapted to generate specific cardiac subtypes—including ventricular and atrial CMs (Lee et al., 2017; Schmidt et al., 2023; Yang et al., 2022)—determining subtype identity in hPSC-CMs remains challenging. Ventricular, atrial, and sinoatrial node CMs possess distinct molecular and physiological features (Lyra-Leite et al., 2022), but the immature, embryonic-like state of hPSC-derived CMs makes established adult markers and electrophysiological profiles poorly applicable for clear subtype distinction (Mummery et al., 2012). This challenge is particularly evident in the controversy that persists regarding the chamber-specific identity of hPSC-CMs produced by the standard WNT ON/OFF protocol. The ventricular protein MYL2 (MLC2v) has been the most extensively used marker for subtype identity characterization, yet results across multiple laboratories show substantial variability in its expression, ranging from 5% to ∼90% depending on the day of evaluation and protocol variations (Table 1). Recently, Dark et al observed only ∼35% HAND1+/MYL2+ double-positive cells at day 20 using the standard WNT ON/OFF method, interpreting this low co-expression as evidence for mixed cardiac subtypes and concluding that the protocol generates heterogeneous populations requiring media optimization for LV-CM enrichment (Dark et al., 2023). However, MYL2 expression exhibits maturation-dependent upregulation (Burridge et al., 2014), raising the question of whether early low expression reflects incomplete maturation rather than lineage heterogeneity. Consistent with this interpretation, prior studies using extended culture periods demonstrated progressive MYL2 enrichment over time: Cyganek et al and Biermann et al reported ∼90% MYL2+ cells by days 80-120 (Biermann et al., 2019; Cyganek et al., 2018), while Luo et al documented 87% by day 60 (Luo et al., 2021). These latter temporal studies collectively suggest that WNT ON/OFF produces predominantly ventricular CMs when allowed sufficient maturation time. However, none assessed whether this ventricular population represented left ventricular, right ventricular, or a mixture of both chamber identities.

**Table 1.** LV Characterization of WNT ON/OFF Protocols. : Summary of selected studies reporting hiPSCs-CMs MYL2 levels, obtained using standard WNT ON/OFF differentiation protocols

| Study | Year | Day of assessment | Enrichment | Maturation | Characterization read out |
| --- | --- | --- | --- | --- | --- |
| (Guo et al., 2024) | 2024 | 30 | None | None | **80% MYL2+ CMs** |
| (Galdos et al., 2023) | 2023 | 30 | 2-day glucose deprivation | None | **30% and 69% TBX5+/MYL2+ CMs** (depending on cell line) |
| (Dark et al., 2023) | 2023 | 20 | 2-day lactate enrichment | None | **35% HAND1+/MYL2+ CMs** |
| (Hsueh et al., 2023) | 2023 | 14 | None | None | **<10% MYL2+ CMs** |
| (Luo et al., 2021) | 2021 | 60 | None | Temporal | **87% MYL2+ CMs** |
| (Chirikian et al., 2021) | 2021 | 30 | None | None | **55% MYL2+ CMs** |
| (Biermann et al., 2019) | 2019 | 30 and 80 | None | Temporal | **19% (D30) and 94% (D80) MYL2+ CMs** |
| (Cyganek et al., 2018) | 2018 | 80-120 | 5-day lactate enrichment | Temporal | **89% MYL2+ CMs** |
| (Burridge et al., 2014) | 2014 | 60 | 10-day lactate enrichment | Temporal | **60% MYL2+ CMs** |
| (Lian et al., 2012) | 2012 | 20, 40, 60 | None | Temporal | **5% (D20), 20% (D40)**<br>**and 60% (D60)**<br>**MYL2+ CMs** |

Galdos et al made important advances by employing TBX5-Cre lineage tracing on a MYL2 reporter line to specifically address FHF and LV identity (Galdos et al., 2023). They demonstrated that more than 90% of CMs were TBX5-lineage positive, and their transcriptional profiling revealed ventricular gene expression signatures consistent with FHF origin and left ventricular fate. However, the progenitor characterization relied on bulk RT-qPCR temporal dynamics rather than direct population-level protein profiling, precluding definitive exclusion of SHF progenitor contributions. Most critically, substantial heterogeneity persisted: only 30% and 69% of CMs (depending on the cell line) expressed MYL2 protein at day 30, leaving the complementary population of TBX5+/MYL2- CMs without comprehensive single-cell protein characterization. Collectively, these studies suggest that WNT ON/OFF generates predominantly ventricular CMs, as evidenced by lineage tracing, transcriptional signatures, and time-dependent MYL2 enrichment. However, a critical gap remains: no study has definitively demonstrated high-purity LV-CM populations through comprehensive temporal protein expression analysis combining ventricular markers (MYL2) with other chamber-specific markers (HAND1 for FHF/LV/OFT, TBX5 for FHF/LV/ATRIAL). Without such integrated analysis demonstrating simultaneous expression of multiple chamber-defining markers in mature cells, fundamental questions remain unresolved regarding whether the MYL2 heterogeneity observed at early-to-intermediate timepoints reflects genuine lineage diversity or incomplete maturation of specified LV-CMs.

Here, we systematically address this gap by combining single-cell protein analysis of cardiac progenitors and CMs with targeted metabolic maturation. We demonstrate that the WNT ON/OFF method consistently generates HAND1+/NKX2.5+ FHF progenitors at the single-cell protein level, a critical validation not achieved in prior studies. We show that the MYL2-negative population observed in standard cultures represents immature LV-CMs that have not yet acquired their definitive protein markers, rather than alternative cardiac subtypes. Through temporal analysis and metabolic maturation, we achieve 95% MYL2+/HAND1+/TBX5+ triple-positive LV-CMs by day 38, a substantially higher purity and earlier timepoint than previously reported. This phenotypic resolution is accompanied by advanced structural maturation features, notably including the assembly of polarized XIRP2+ intercalated discs, a hallmark of postnatal cardiac maturation. Validated across three independent hPSC lines, our findings resolve the field gap by demonstrating that apparent subtype heterogeneity largely reflects maturation state rather than true lineage diversity, and establish a rapid, high-fidelity platform for generating pure mature LV-CMs for cardiovascular research.

## RESULTS

### The WNT ON/OFF protocol consistently generates cardiac cells through an intermediate state corresponding to the first heart field

Since its first description in 2011 by Gonzalez et al., the WNT ON/OFF small molecule cardiac differentiation protocol has rapidly become a key method for generating hPSCs-CMs. Quantitative bibliographic analysis allowed us to see that among approximately 1800 freely available papers in PubMed Central describing cardiac differentiation of hPSCs, more than 60% of publications between 2012 and 2025 employ a variation of this 2D protocol (Figure S1A). Embryoid body protocols represented approximately 55% in 2011, with this number subsequently decreasing to the roughly 10% observed up until mid 2025 (Figure S1B). In contrast, the frequency of small molecule WNT ON/OFF protocols has rapidly increased, representing roughly 75% of papers published in 2025. Given its widespread adoption, characterizing the cardiac subtypes generated by this protocol is of critical importance to the field.

To assess which CM subtypes are generated using the WNT ON/OFF small molecule method, we first evaluated the protocol efficiency and reproducibility under different conditions using the human induced pluripotent stem cell (hiPSCs) FN line (Figure 1A). Fine-tuning of GSK-3β inhibitor CHIR99021 (CHIR) during initial differentiation is fundamental for achieving high cardiac induction efficiency from hPSCs (Burridge et al., 2015; Lewis-Israeli et al., 2021; Lian et al., 2012). This concentration-dependent effect aligns with the role of morphogens like BMP4 and Activin A during primitive-streak and mesoderm induction, where precise signaling modulation is critical for cardiac subtype specification (Lee et al., 2017). This led us to examine whether CHIR could also affect cardiac subtype identity.

**Figure 1.**
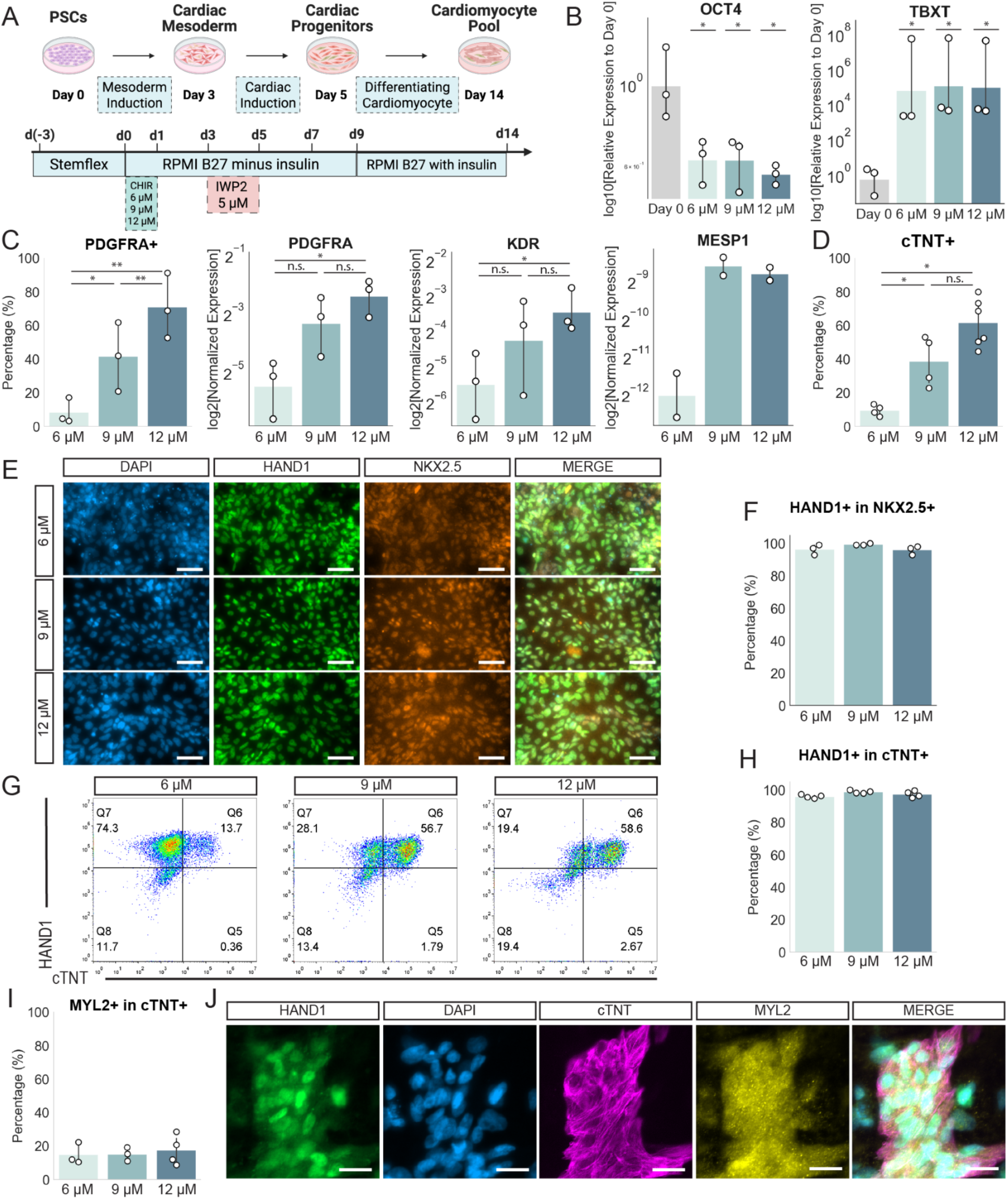
WNT ON/OFF differentiation generates cardiomyocytes via a first heart field–like progenitor stage. A) Schematic of the cardiac differentiation protocol used. CHIR was evaluated at 6 µM, 9 µM, and 12 µM for the first 24 h. B) RT-qPCR of pluripotency marker OCT4 and mesodermal marker TBXT at day 1. Statistic comparison was performed between D0 and 6 μM, 9 μM or 12 μM independently. n=3. C) Left: flow cytometry of PDGFRA+ cells at day 4. Middle and right: RT-qPCR of PDGFRA, KDR, n=3, and MESP1, n=2. D) Flow cytometry of cTNT+ cells efficiency at day 14, n=3. E) Representative immunofluorescence stainings of NKX2.5, HAND1, and DAPI in CMs at day 7. Scale bar = 50 µm. F) Image cytometry quantification of HAND1+/NKX2.5+ expression at day 7, n=3. G) Representative flow cytometry of HAND1+/cTNT+ CMs at day 14. H) Flow cytometry quantification of HAND1+ within cTNT+ CMs at day 14. n=3. I) Flow cytometry quantification of MYL2+ cells within cTNT+ population at day 14, n=3 J) Immunofluorescence staining of HAND1, MYL2, cTNT, and DAPI in CMs at day 14, n=3. Scale bar = 20 µm. Data are represented as mean and individual values. *p < 0.05, **p < 0.01; n.s., not significant.

We evaluated a broad range of CHIR concentrations (6, 9, and 12 μM) to determine their effects on cardiac differentiation. In all conditions, CHIR effectively downregulated the pluripotency marker *OCT4* by day 1 while inducing primitive streak and mesendodermal markers *TBXT*, *LEF1*, and *EOMES* (Figure 1B, Figure S2A). By day 4, we observed concentration-dependent increases in the broad mesoderm marker PDGFRα at mRNA and protein levels, with PDGFRα-positive cells increasing from 10% at 6 μM CHIR to ∼70% at 12 μM CHIR, suggesting more efficient mesoderm specification at higher concentrations (Figure 1C). A similar trend occurred for the cardiac mesoderm marker *KDR*. The transcription factor *MESP1*, the earliest transcriptional regulator committing hPSCs to cardiovascular lineage, also showed increased expression with higher CHIR concentrations. At day 14, while 6 μM CHIR yielded ∼10% cardiac troponin T-positive (cTNT+) cells, higher concentrations induced higher cTNT+ percentages, with 12 μM reaching a mean of 60% cTNT+ CMs (Figure 1D). Moreover, we observed no beating structures at 6 μM CHIR, while 9 and 12 μM treatments generated large beating monolayer clusters (Videos S1-S3). Notably, published protocols report variable optimal CHIR concentrations, with some groups achieving high efficiency at 6 μM while others systematically employ higher concentrations (Dark et al., 2023; Lian et al., 2012). Given the pronounced concentration-dependent effects we observed, these inter-laboratory differences likely reflect batch-to-batch and supplier variability in CHIR potency. Our findings highlight the importance of appropriate WNT signaling strength for efficient cardiac mesoderm induction and CM specification.

We next investigated whether CHIR concentration could influence cardiac progenitor identity beyond overall differentiation efficiency. To assess whether heart field derivatives are modulated by CHIR concentration, we evaluated the FHF marker HAND1 co-expression within NKX2.5-positive cardiac progenitors at day 7. Cardiac progenitors of the FHF express HAND1 at low levels, compared to high expression in extraembryonic mesoderm and epicardial progenitors (Lynch et al., 2025). Notably, across all conditions, almost all NKX2.5-positive cells expressed HAND1 at low levels, strongly indicating FHF identity irrespective of CHIR concentration (Figure 1E-F).

We subsequently evaluated cardiac subtype markers within cTNT-positive CMs at day 14. In agreement with our previous observations, more than 95% of cTNT-positive CMs expressed HAND1 across all CHIR concentrations, a marker positive in both LV and OFT CMs (Figure 1G,H). However, only ∼15% of cTNT+ CMs expressed ventricular marker MYL2 at day 14 (Figure 1I). Consistent with these results, immunofluorescence analysis revealed MYL2+ CMs exhibited diffuse cytoplasmic localization rather than the characteristic sarcomeric patterns of mature CMs (Figure 1J). Similar results were obtained in two independent PSC lines, H1 human embryonic stem cells and the DESS hiPSC line recently derived in our lab (Castañeda et al., 2023). Both lines showed all cTNT-positive CMs at day 14 expressed nuclear HAND1 while showing diffuse and low expression of MYL2 (Figure S2B).

Our results indicate that at this early differentiation timepoint the majority of CMs are HAND1+ and MYL2-, which could be initially compatible with an OFT subtype. However, given that MYL2 exhibits maturation-dependent upregulation during cardiac differentiation, we hypothesized that the apparent cardiac subtype heterogeneity marked by MYL2-negative CMs could reflect varying maturation states rather than true lineage diversity.

### MYL2-negative CMs co-express HAND1 and TBX5, indicating left ventricular identity in an immature state

To further investigate CM subtype identity, we evaluated cardiac markers at later timepoints in lactate-purified populations (Figure 2A). This metabolic enrichment exploits CMs’ unique ability to utilize lactate while partially inducing cardiac maturation (Tohyama et al., 2013; Wickramasinghe et al., 2022). Following purification and culture to day 26, we reassessed CM subtype markers across CHIR conditions (9 and 12 μM; 6 μM was excluded due to low cardiac differentiation efficiency and poor lactate selection survival). Since MYL2 showed diffuse cytoplasmic localization at day 14, we evaluated both expression levels and localization using quantitative image cytometry. On day 26, this ventricular marker displayed robust fibrillar expression in 75-80% of cTNT-positive cells (Figure 2B,C, Figure S3A). The remaining 20% still exhibited sarcomeric MYL2 expression, albeit at lower levels (Figure 2B, arrows). The substantial increase in MYL2+ cells from day 14 (∼15%) to day 26 (∼75%), plus its sarcomeric assembly, is consistent with the well-documented maturation-dependent upregulation of MYL2 rather than with a variation in the cardiac subtype distribution. Critically, all CMs from both CHIR concentrations exhibited co-expression of FHF-derived and LV markers HAND1 and TBX5, indicating that the MYL2-low population represents immature LV CMs rather than alternative cardiac lineages (Figure 2D,E). Similar results were obtained in the H1 and DESS PSC lines (Figure S3B). Thus, our multi-marker analysis suggests that MYL2 heterogeneity at intermediate timepoints reflects maturation state rather than lineage diversity, underscoring the necessity of comprehensive marker co-expression analysis for definitive subtype identification.

**Figure 2.**
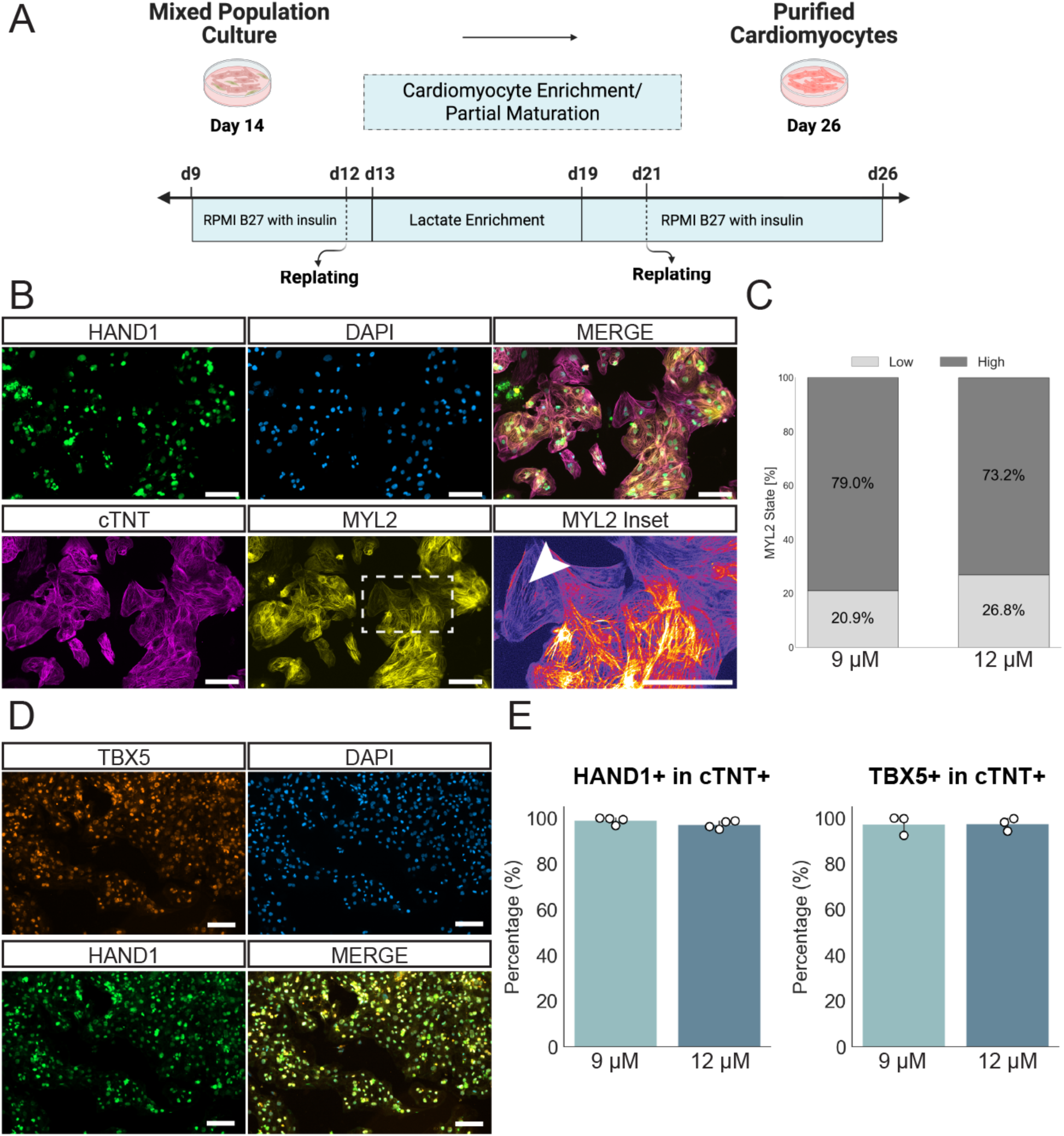
Extended culture and lactate selection enhance MYL2 expression in cardiomyocytes. A) Schematic of the lactate enrichment protocol used to purify CMs. B) Representative immunofluorescence staining of HAND1, MYL2, cTNT, and DAPI of CMs at day 26 for the 12 µM CHIR condition. The bottom right panel shows a zoom-in of the sarcomeric display of MYL2. Scale bar = 100 µm. C) Image cytometry quantification of MYL2 in cTNT+ expression states for 9 µM and 12 µM CHIR conditions, n=4. D) Immunofluorescence staining of HAND1, TBX5, and DAPI of CMs at day 26. Scale bar = 100 µm. E) Image cytometry quantification of HAND1 (n=4) and TBX5 (n=3) expression for 9 µM and 12 µM CHIR conditions. Data are represented as mean and individual values.

### Metabolic maturation enhances cardiomyocyte structural and functional properties

To eliminate the confounder of an immature phenotype on MYL2 expression, we adopted a metabolic maturation protocol designed to promote CM structural and functional maturation (Funakoshi et al., 2021). This protocol was originally optimized for CMs differentiated via embryoid bodies using BMP4, Activin A, and FGF2. Thus, we first characterized this protocol in detail for the 2D WNT ON/OFF method after lactate purification (Figure 3A). The maturation medium (MM) is based on previous reports demonstrating the maturation effects of culturing CMs in low glucose with fatty acids (palmitic acid), hormones (T3, dexamethasone), and a PPAR-α agonist (Correia et al., 2017; Feyen et al., 2020; Gentillon et al., 2019; Parikh et al., 2017; Yang et al., 2014, 2019).

**Figure 3.**
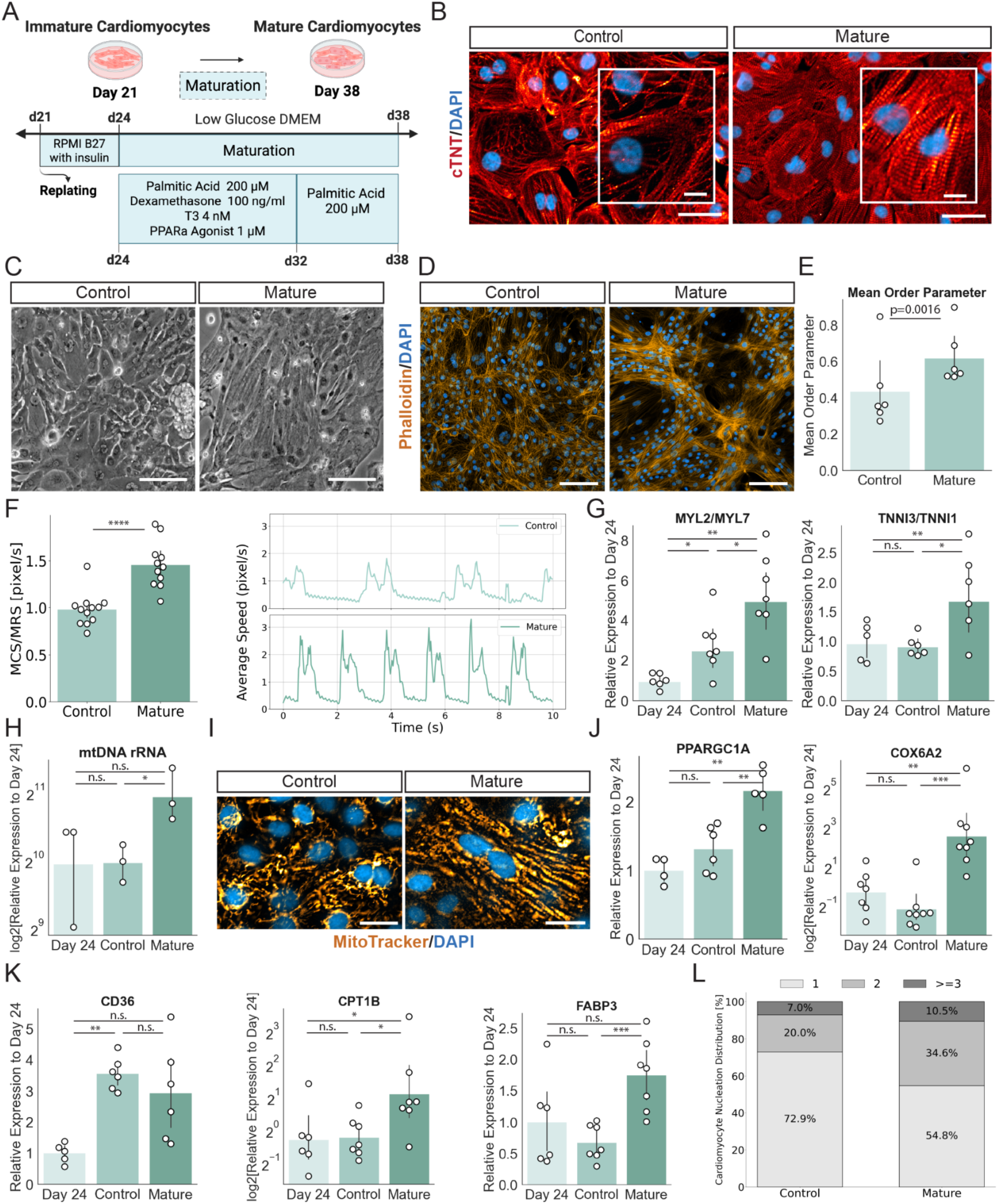
Metabolic maturation enhances structural alignment, contractile performance, and mitochondrial function in cardiomyocytes. A) Schematic of the maturation protocol applied after day 23. B) Representative immunofluorescence staining of cTNT. Scale bar = 100 µm; inset = 20 µm. C) Brightfield images of control (day 38 RPMI with B27) and mature CMs (day 38 maturation media). Scale bar = 100 µm. D) Immunofluorescence staining of phalloidin (F-actin) in control and mature CMs. Scale bar = 100 µm. E) Quantification of cellular alignment using the mean order parameter derived from a 2D Fourier transform algorithm, n=6. F) Contractility analysis showing MCS/MRS ratio quantification and an example of contraction curves, n=5 G) RT-qPCR of sarcomeric isoform switches: MYL2/MYL7 and TNNI3/TNNI1 in day 24, control, and mature cells, n=6. H) RT-qPCR quantification of mtDNA rRNA abundance at day 24, control and mature CMs, n=3. I) Immunostaining with MitoTracker Green FM and DAPI to visualize mitochondrial distribution. Scale bar = 20 µm. J) RT-qPCR of mitochondrial gene expression in control and mature CMs, n=6. K) RT-qPCR of fatty acid uptake genes in control and mature CMs, n=6. L) Quantification of nucleation levels in control vs. mature CMs by DAPI and membrane staining, n=3. Data are represented as mean and individual values. *p < 0.05, **p < 0.01, ***p < 0.001; n.s., not significant.

For these experiments, we selected CMs derived using 12 μM CHIR, as this concentration provided the most consistent differentiation efficiency.

To assess the effect of the maturation media we cultured purified day 24 CMs in either MM or control RPMI medium until day 38, thus analyzing three experimental conditions: Day 24 (D24), Day 38 in MM (Mature), and Day 38 in RPMI (Control). At day 38 we observed major morphological differences between Mature and Control conditions. cTNT immunostaining and widefield microscopy showed that matured cells exhibited a marked increase in sarcomeric density and alignment, as well as a compact phenotype with indivisible cell boundaries, hallmarks of advanced structural maturation (Figure 3B,C, Figure S4A, Video S4-S5). Interestingly, in wells subjected to MM, CMs aligned into multicellular fascicles and interconnected networks, with individual cells forming extensive bundles of actin filaments (Figure 3D). By contrast, cells maintained in the control medium persisted as a flat, uniform monolayer and did not assemble into higher-order structures. This behavior, reminiscent of the architecture of native myocardium, could be efficiently quantified by analyzing the degree of alignment using 2D Fourier transform-based order parameter calculations (Figure 3E).

To determine whether the observed morphological changes were accompanied by enhanced functional properties, we analyzed CM contractility using time-lapse imaging. We quantified the Mean Contraction Speed (MCS) and Mean Relaxation Speed (MRS) in both conditions and found that the MCS was significantly increased in the mature group, resulting in a ∼50% higher MCS/MRS ratio (Figure 3F). This indicates that the maturation protocol effectively enhanced the upstroke speed of contraction, providing functional evidence consistent with improved contractile kinetics and overall maturation of WNT ON/OFF-derived CMs.

We next investigated whether structural maturation was accompanied by isoform-level remodeling of sarcomeric proteins, an established hallmark of CM developmental progression (Guo and Pu, 2020). Specifically, we focused on shifts in the expression ratios of key sarcomeric isoforms that distinguish fetal from mature ventricular CMs. The MM induced an increase in the MYL2/MYL7 ratio compared to both day 24 (pre-treatment) and day 38 control conditions measured by RT-qPCR (Figure 3G). This observation reflects the developmental switch from atrial/embryonic myosin light chain (MYL7) to ventricular/adult myosin light chain (MYL2), and is in line with the view that evaluation of CM subtype exclusively by MYL2 expression on immature cells could underestimate the percentage of ventricular cells. Similarly, we observed strong upregulation of the adult cardiac troponin isoform TNNI3 relative to the fetal isoform TNNI1 in mature conditions, resulting in an increased TNNI3/TNNI1 ratio, a hallmark of CM maturation (Figure 3G, Figure S4B). Collectively, these results demonstrate that the metabolic maturation protocol successfully recapitulates key aspects of *in vivo* cardiac development, driving both structural reorganization and molecular remodeling toward a more adult-like phenotype.

### Cardiomyocyte maturation involves coordinated metabolic, mitochondrial, and nuclear remodeling

CM maturation involves a coordinated transition in metabolism. Adult CM rely primarily on fatty acid oxidation (FAO) for energy production, in contrast to the glycolytic metabolism of embryonic or fetal cells (Karbassi et al., 2020). To assess whether our metabolic conditioning protocol promotes this transition, we next evaluated mitochondrial content. This analysis showed comparable mitochondrial load between day 24 and day 38 control conditions, but a significant increase in mature CMs (Figure 3H). Visualization of mitochondria in CMs cultured in MM showed both increased mitochondrial content and a more organized, longitudinal arrangement of these organelles, resembling the inter-myofibrillar alignment found in native myocardium (Figure 3I). These results suggest that metabolic conditioning induces mitochondrial biogenesis, a conclusion supported by the upregulation of PPARGC1A and COX6A2 (Figure 3J), a master regulator of mitochondrial biogenesis and a subunit of the cytochrome c oxidase, respectively.

We next explored changes in metabolic substrate usage by analyzing the expression of key FAO-related genes. Although expression of CD36, a key fatty acid transporter, was unchanged between matured and control CMs, it was elevated in both conditions relative to day 24 (Figure 3K), suggesting that fatty acid uptake increases as an effect of the time of culture. In contrast, downstream FAO genes CPT1B and FABP3 were significantly upregulated in matured CMs, suggesting that the maturation protocol specifically enhances lipid metabolic processing.

Beyond metabolic remodeling, maturation was also evident at the subcellular level through nuclear changes that parallel *in vivo* development. During the course of our analyses, we noted that many CM contained more than one nuclei, which led us to investigate this in further detail. Mammalian fetal CMs are mainly diploid mononucleated while the majority of postnatal human CMs are polyploid mononucleated, with binucleated cells representing more than 25% of this population (Derks and Bergmann, 2020; Leone and Engel, 2019). Nuclear content analysis revealed a substantial increase in multinucleation under maturation conditions, with nearly 44% of cells exhibiting two or more nuclei compared to fewer than 27% in controls (Figure 3L, Figure S4C). Interestingly, we also noted a slight increase in the level of poliplody in mature versus control CMs that changed from 15% in 20%, respectively (Figure S4D). These changes in the DNA nuclear content reflect hypertrophic remodeling via karyokinesis without cytokinesis, a hallmark of postnatal CM development that complements the metabolic and mitochondrial changes observed in our maturation system.

Collectively, these findings demonstrate that the metabolic maturation protocol orchestrates key aspects of CM development, including mitochondrial biogenesis, metabolic remodeling, fatty acid utilization, and changes in nuclear content.

### Metabolic maturation drives transcriptional and post-transcriptional remodeling

To characterize the molecular programs underlying structural and functional maturation, we performed bulk RNA-sequencing on CMs at the three previously mentioned stages: purified day 24 CMs (D24), day 38 in control medium CMs (Control), and day 38 following metabolic conditioning CMs (Mature) (Figure 4A). This experimental design allowed us to distinguish temporal maturation effects from those specifically induced by our metabolic conditioning protocol.

**Figure 4.**
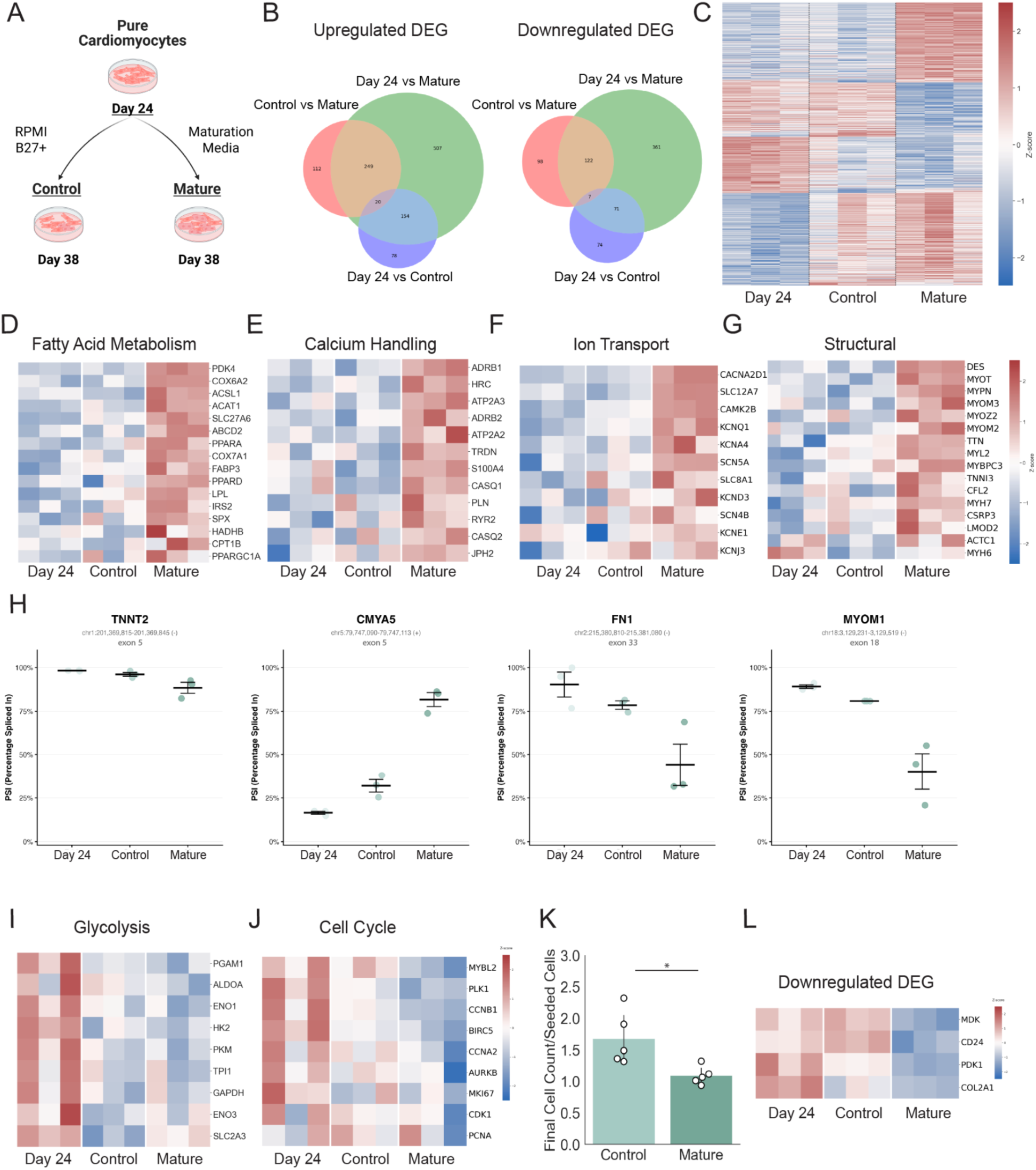
Transcriptomic and phenotypic profiling reveal coordinated metabolic and structural remodelling as well as reduced proliferation during cardiomyocyte maturation. (A) Schematic representation of the sample collection workflow for bulk RNA sequencing. B) Venn diagram showing overlap of upregulated and downregulated DEG across conditions. C) Fuzzy C-Means clustering of the 1,800 DEG across all three conditions. D) Heatmap of genes related to fatty acid metabolism. E) Heatmap of genes related to calcium handling. F) Heatmap of genes involved in ion transport. G) Heatmap of structural genes. H) Alternative splicing PSI plots for TNNT2 (exon 5), CMYA5 (exon 5), FN1 (exon 33), and MYOM1 (exon 18). I) Heatmap of glycolysis-related genes. J) Heatmap of cell cycle-related genes. K) Quantification of total cell number using Neubauer chamber cell counting, n=5 L) Heatmap of downregulated genes. RNA-seq experiment was performed on n = 3 hiPSC-CM batches. Heatmaps display Z-score normalized values. Data are represented as mean and individual values. *p < 0.05, **p < 0.01, ***p < 0.001; n.s., not significant.

Differential gene expression analysis revealed progressive increases in transcriptional changes across conditions, with 404, 608, and 1,491 differentially expressed genes (DEGs) between D24 vs Control, Control vs Mature, and D24 vs Mature, respectively (Figure S5A). Principal component analysis (PCA) confirmed that the three conditions clustered distinctly across all biological replicates (Figure S5B), with the most pronounced transcriptional shift occurring between D24 and Mature conditions. Notably, the majority of DEGs identified in both the D24 vs Control and Control vs Mature comparisons were encompassed within the DEGs of D24 vs Mature (Figure 4B), suggesting that metabolic conditioning substantially amplifies the temporal maturation changes observed with standard RPMI medium. Fuzzy C-Means clustering algorithm of the 1,800 DEGs across all conditions confirmed that Mature CMs formed a distinct transcriptional cluster, clearly separated from both D24 and Control conditions (Figure 4C). Control CMs exhibited an intermediate expression profile, indicating that while extended culture time induces some degree of maturation, the metabolic maturation protocol drives a more comprehensive transcriptional transformation.

Gene Ontology (GO) analysis of genes upregulated in mature CMs revealed significant enrichment of biological processes characteristic of mature CMs, including hypertrophic cardiac growth, sarcomere organization, gap junction assembly, T-tubule organization, regulation of fatty acid oxidation, and ventricular cardiac muscle development (Table S1). Consistent with the expected metabolic shift toward oxidative phosphorylation, we identified robust upregulation of fatty acid metabolism genes, including COX6A2, COX7A1, LPL, ACAT1, PDK4, PPARD, and CPT1B (Figure 4D). Gene Set Enrichment Analysis (GSEA) corroborated this finding, demonstrating significant enrichment of FAO pathways in matured cells (Figure S5C), evidencing the transition from glycolytic to oxidative metabolism that characterizes the functional cardiac maturation.

The maturation program also encompassed critical calcium handling and ion transport machinery. Calcium cycling genes (ADRB1/2, CASQ1/2, ATP2A2/3) and ion channel genes (KCNJ2, SCN5A, ADRB2, KCNQ1, CACNA2D1) were significantly upregulated, supporting the enhanced electrophysiological properties observed in mature CMs (Figures 4E, F). Notably, we observed upregulation of the T-tubule organization genes ANK2, CAV3, CSRP3 (Parikh et al., 2017), which are essential for establishing the mature excitation-contraction coupling architecture that enables efficient calcium release and uptake between the sarcoplasmic reticulum and contractile apparatus (Figure S5D). Additionally, we observed an upregulation of the thyroid hormone receptor THRA and glucocorticoid receptor NR3C1 (Figure S5E), consistent with activation of these signaling pathways by the maturation medium.

Our transcriptional analysis revealed a coordinated upregulation of genes encoding key cardiac structural components and sarcomeric proteins. Among these are the intermediate filament protein desmin (DES), and sarcomeric proteins MYOT, TTN, MYOM3, and MYPN (Figure 4G). The upregulation of these structural genes supports the observation of significantly denser sarcomeric structures observed in the mature conditions (see Figure 3B).

Cardiac maturation involves not only transcriptional changes but also extensive alternative splicing remodeling that fine-tunes protein isoform composition at the post-transcriptional level (Mazin et al., 2021). To assess whether our maturation protocol recapitulates these developmentally regulated splicing patterns, we analyzed percent spliced-in (PSI) values for cardiac genes with well-characterized maturation-dependent exon usage. Consistent with the established postnatal exclusion of TNNT2 exon 5 (Jin and Lin, 1989; Wei and Jin, 2016), mature CMs showed significantly reduced exon 5 inclusion compared to day 23 and day 38 control conditions (Figure 4H). Similarly, we observed maturation-associated exon exclusion in MYOM1 (exon 18) and FN1 (exon 33), and increased exon inclusion in CMYA5 (exon 5), all matching the splicing transitions recently characterized during human cardiac development (Gomes-Silva et al., 2025). These coordinated splicing changes across multiple cardiac structural genes provide independent molecular evidence that metabolic conditioning drives comprehensive maturation at both transcriptional and post-transcriptional levels.

Complementing these multi-layered regulatory changes, GO analysis of downregulated genes in mature CMs revealed significant association with glycolytic processes, an observation supported by gene set enrichment analysis (Figure 4I, Figure S5F, Table S2). This downregulation, coupled with the upregulation of oxidative metabolism genes, confirms the expected metabolic reprogramming from glycolysis to oxidative phosphorylation. Importantly, mature CMs exhibited reduced expression of cell cycle genes associated with mitotic progression, including MYBL2, BIRC5, PLK1, KI67, and CCNB1 (Figure 4J). Evaluation of CM proliferation by standard methods is difficult due to the confounding effects of endoreplication, polynucleation and hypertrophy (Bois et al., 2025). To validate the transcriptional evidence of proliferative arrest, we quantified cell numbers on day 38 after plating equal numbers of CMs on day 21 in each condition. While control CMs maintained in RPMI exhibited approximately 50% increase in cell number, metabolically matured CMs showed no significant change with respect to the number of cells plated (Figure 4K), confirming cell cycle exit characteristic of postnatal CM development.

Finally, the RNA-seq analysis also allowed us to identify additional genes that are specifically downregulated in mature CMs compared to control conditions (Figure 4L). Among these, CD24 (heat stable antigen, HSA) has been recently shown to mark a population of immature CMs in adult mice, that retain the capacity to proliferate within the first postnatal week and that later in development remain as mononucleated cells (Valente et al., 2019).

As a whole, our findings demonstrate that the metabolic maturation protocol orchestrates key aspects of CM development, including mitochondrial biogenesis, structural and metabolic remodeling, fatty acid utilization, ion transport reorganization, and proliferative arrest. Together, these features define a transition toward a more mature and physiologically relevant cardiac phenotype.

### Metabolic maturation initiates polarized intercalated disc assembly

Among the most striking transcriptional changes in mature CMs was the robust upregulation of genes encoding intercalated disc (ICD) components (Estigoy et al., 2009). ICDs are highly specialized structures at CM termini that coordinate mechanical coupling and electrical synchronization in the myocardium (Nielsen et al., 2023). In mammals, ICDs assemble primarily during postnatal development, representing a critical hallmark of cardiac maturation not previously described in the literature for hPSC-CMs in standard 2D conditions (Karbassi et al., 2020; Vreeker et al., 2014). Comprehensive analysis of ICD-associated genes revealed widespread upregulation of this protein network in mature CMs (Figure 5A). These included essential components of mechanical junctions (desmosomal proteins such as DSC2), electrical coupling (KCNJ2, SCN5A, GJA1), and anchoring complexes (XIRP2, NRAP, ANK, MYZAP). This coordinated transcriptional program suggested that metabolic conditioning initiates the assembly of intercalated disc architecture.

**Figure 5.**
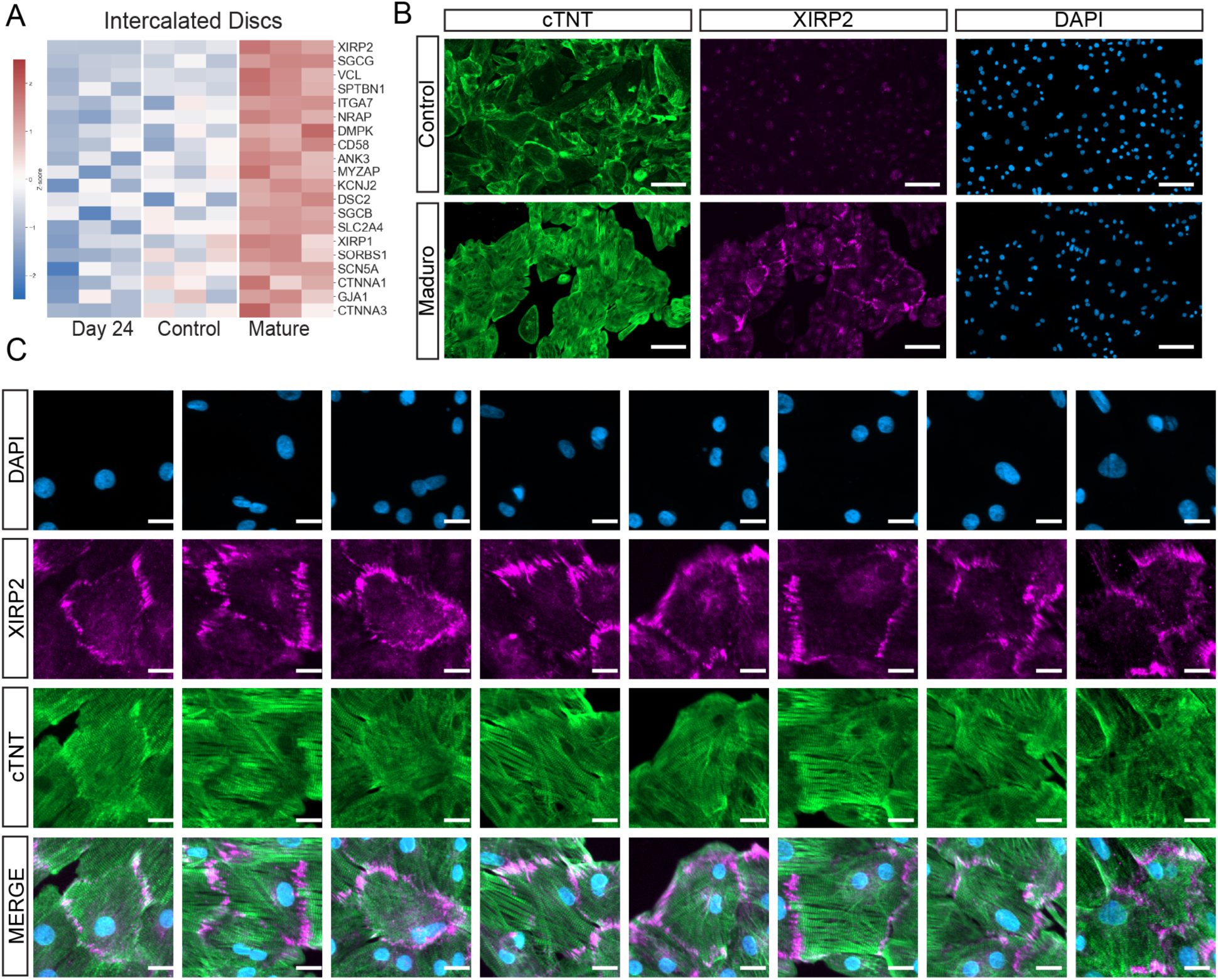
Metabolic maturation induces upregulation of intercalated disc components and polarized XIRP2 assembly. A) Heatmap showing differential expression of intercalated disc-associated genes across day 24, day 38 control, and day 38 mature conditions (Z-score normalized). B) Representative low-magnification immunofluorescence images of day 38 control and mature CMs stained cTNT, XIRP2, and nuclei (DAPI). Control CMs show no detectable XIRP2 expression, while mature CMs display robust XIRP2 protein accumulation. Scale bars= 100 µm. C) High-magnification images showing heterogeneous XIRP2 localization patterns in mature CMs. XIRP2 distribution ranges from circumferential membrane localization to polarized concentration at cellular termini, indicating progressive stages of intercalated disc assembly. Scale bars= 20 µm.

To validate ICD assembly at the protein level, we focused on XIRP2, the most highly upregulated ICD gene in our transcriptome (fold change of ∼40 between Mature and Control conditions) and a protein shown to be among the earliest to polarize during postnatal development. XIRP2 is essential for redistribution of intercellular junction components from lateral membrane locations to cardiomyocyte termini, establishing the spatial organization characteristic of mature ICDs (Wang et al., 2013). Immunofluorescence analysis revealed substantially different expression patterns between conditions: while day 38 control CMs showed no detectable XIRP2 expression in cell membranes across the entire population, mature CMs displayed robust XIRP2 protein accumulation (Figure 5B). At the single cell level, XIRP2 localization in mature CMs exhibited heterogeneity, with some cells showing circumferential membrane distribution and others displaying clear polarized concentration at cellular termini (Figure 5C). This polarized distribution at cell ends, while not yet uniform across all mature CMs, represents direct evidence of ICD structural assembly occurring at the protein level. The presence of polarized XIRP2 in a substantial fraction of cells suggests that metabolic conditioning drives CMs toward postnatal-stage maturation, initiating the spatial reorganization of ICD components that defines mature myocardial architecture.

### The WNT ON/OFF protocol generates a homogeneous population of left ventricular cardiomyocytes when assessed after metabolic maturation

Having established that the maturation protocol promotes a phenotypic transition with hallmarks of postnatal CM development, we next sought to conclusively determine cardiac subtype identity through comprehensive multi-marker analysis. As expected, RT-qPCR analysis revealed progressive MYL2 upregulation across our experimental timeline, with mature CMs (day 38) exhibiting significantly higher expression than both day 24 and day 38 control conditions (Figure S6A). This temporal pattern was then validated at the protein level. Flow cytometry and quantitative image cytometry revealed that more than 95% of mature CMs expressed MYL2 at day 38 (Figure S6B,C), in agreement with previous temporal studies demonstrating progressive ventricular enrichment in WNT ON/OFF-derived CMs (Biermann et al., 2019; Cyganek et al., 2018; Luo et al., 2021). However, our metabolic maturation approach achieves this degree of ventricular marker expression substantially earlier, by day 38 compared to the 60-120 days required in prior reports, providing a more efficient route to mature ventricular populations.

In agreement with our previous results, multi-marker single-cell analysis revealed that more than 95% of mature CMs co-expressed the chamber-defining combination of MYL2, HAND1, and TBX5 (Figure 6A, Figure S6D), a triple-positive signature specific for left ventricular identity. This near-homogeneous co-expression pattern demonstrates that metabolic maturation is essential for revealing the complete left ventricular phenotype: while HAND1 and TBX5 were already expressed at earlier timepoints, the maturation-dependent expression of MYL2 in the majority of the cells resolves the apparent heterogeneity observed with standard culture conditions.

**Figure 6.**
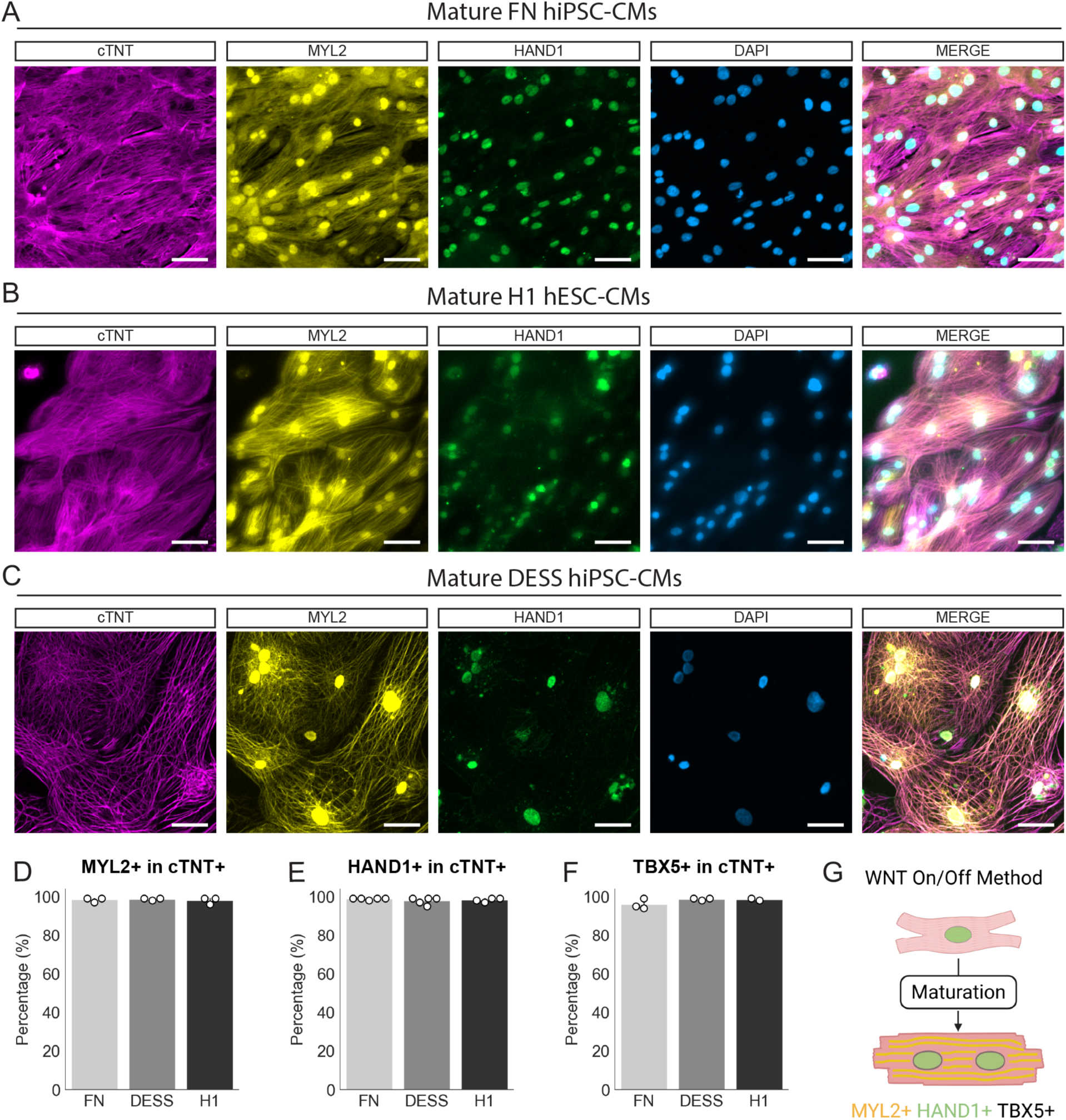
Mature cardiomyocytes derived via WNT ON/OFF differentiation exhibit strong left ventricular identity. A,B,C) Representative immunofluorescence stainings of HAND1, MYL2, cTNT, and DAPI in mature CMs for the FN hiPSC, H1 hESC and DESS hiPSC cell lines. Scale bar = 100 µm. D) Image cytometry quantification of MYL2 within the cTNT+ population for the three cell lines, n=3. E) Image cytometry quantification of HAND1 within the cTNT+ population for the three cell lines, n=4. F) Image cytometry quantification of TBX5 within the cTNT+ population for the three cell lines, n=2. Data are represented as mean and individual values. G) Proposed model for LV identity of mature WNT ON/OFF CMs.

Mature CMs were negative for the left atrial-specific marker PITX2 and the cardiac conduction system marker TBX3 (Figure S6E), confirming the absence of these lineages. Consistent with this ventricular identity, only approximately 5% of cells expressed the atrial marker NR2F2, with these NR2F2-positive cells largely corresponding to the small MYL2-negative fraction (Figure S6F), reinforcing minimal contamination with other cardiac subtypes.

To generalize our findings across different hPSC lines, we generated day 38 mature CMs from the H1 hESC line and the DESS hiPSCs. In agreement with our previous results, more than 95% of CMs from these additional cell lines were positive for MYL2, TBX5, and HAND1 (Figure 6B-F). Interestingly, in all three cell lines tested, we observed MYL2 localization both in sarcomeres and in the nucleus. This nuclear localization pattern is consistent with results previously observed in rat myocardium (Zhang et al., 2015) and represents an intriguing finding that warrants future investigation.

Overall, our results establish that the standard WNT ON/OFF protocol generates a near-homogeneous population of left ventricular cardiomyocytes. The multi-marker analysis at the single-cell level, combined with metabolic maturation, resolves the longstanding uncertainty about cardiac subtype identity in WNT ON/OFF-derived CMs. By demonstrating that more than 95% of mature CMs co-express MYL2, HAND1, and TBX5 across three independent hPSC lines, we provide evidence for left ventricular specification and establish a validated platform for generating high-purity mature LV-CMs for cardiovascular research (Figure 6G).

## DISCUSSION

This study resolves a longstanding question in cardiac differentiation: we show that the standard WNT ON/OFF protocol generates cardiomyocytes with consistent left ventricular identity when assessed in appropriately matured cells. Our systematic analysis demonstrates that the apparent subtype variability reported across the literature largely reflects differences in maturation state rather than true lineage heterogeneity. By combining single-cell protein analysis of CMs with targeted metabolic maturation, we provide a rapid, high-fidelity platform for generating mature LV-CMs with postnatal features more suitable for modeling cardiac development and disease.

MYL2 has traditionally been amongst the most used protein markers to determine CM subtype identity (Burridge et al., 2015; Chirikian et al., 2021; Cyganek et al., 2018; Dark et al., 2023; Galdos et al., 2023; Lian et al., 2012). However, the reported variability in its expression has fueled uncertainty about WNT ON/OFF CM chamber subtype, with ventricular yields ranging from 5% to 90% depending on timing and methodology (see Table 1). While it has been widely reported that MYL2 expression is maturation dependent (Burridge et al., 2014), and that WNT ON/OFF generates approximately 90% MYL2+ ventricular CMs with extended culture (60-120 days) (Biermann et al., 2019; Cyganek et al., 2018; Luo et al., 2021), these latter studies did not address right versus left chamber identity in these conditions. At the same time, recent publications raise concern about the subtype identity of the standard WNT ON/OFF protocol (Dark et al., 2023). Thus, there is a need in the field to provide faithful and reproducible protocols that generate homogeneous populations of LV-CMs.

Our temporal analysis is in agreement with MYL2 dependency on maturation status: the progression from 15% MYL2+ cells at day 14 to 80% at day 26, and ultimately 95% in metabolically matured cells at day 38, demonstrates that low MYL2 expression in early-stage cells cannot be interpreted as evidence for non-ventricular identity. Importantly, our metabolic conditioning approach achieves comparable or higher MYL2+ CMs substantially earlier than previously reported, by day 38 versus 60-120 days. This time reduction offers significant advantages in cost, efficiency, and experimental throughput, particularly relevant given that WNT ON/OFF protocols now represent more than 75% of cardiac differentiation methods in current use.

Our findings both complement and substantially advance recent studies addressing WNT ON/OFF chamber identity. Galdos et al provided elegant lineage-tracing evidence that more than 90% of CMs derive from TBX5-positive progenitors with transcriptional signatures consistent with FHF and left ventricular fate (Galdos et al., 2023). However, their observation that only 30-69% of cells co-expressed TBX5 and MYL2 protein at day 30 left unresolved whether this represented mixed lineages or incomplete maturation. Our work directly addresses this gap. By combining single-cell protein quantification of FHF identity at the cardiac progenitor stage with multi-marker analysis of mature CMs (MYL2, HAND1, TBX5), we demonstrate that the MYL2-negative fraction within the TBX5-lineage population represents immature but specified LV-CMs. Through metabolic maturation, we achieve 95% MYL2+/HAND1+/TBX5+ triple-positive cells, with only a small fraction of NR2F2+ atrial cells. This demonstrates that metabolic maturation unlocks the left ventricular phenotype at the protein level, resolving the heterogeneity observed by Galdos et al and establishing a platform suitable for modeling cardiac disease and development. The validation across three independent hPSC lines establishes the generalizability and robustness of our approach.

Beyond resolving the chamber identity question, we show that the metabolic maturation protocol developed by Funakoshi et al specifically for CMs derived from EB protocols works effectively in the 2D WNT ON/OFF monolayer protocol. While this cross-compatibility might be expected given the shared principles of cardiac differentiation, our work provides important validation that this maturation protocol, which combines key treatments including fatty acids, low glucose, dexamethasone, PPAR agonists, and T3, functions well in the differentiation method most widely used in the field. There is currently a wide variety of maturation protocols that incorporate different stimuli such as mechanical stretch, electrical stimulation, three-dimensional culture, and various metabolic or hormonal treatments (Maroli and Braun, 2021). While many of these protocols faithfully recapitulate features of the mature state, the lack of standardization and reproducibility between maturation protocols presents a challenge to the field. Our systematic characterization of a well-defined metabolic maturation protocol in the context of 2D WNT ON/OFF differentiation provides a reproducible foundation that can be adopted broadly.

Our transcriptional profiling revealed robust enrichment of processes characteristic of mature CMs, including sarcomere organization, fatty acid oxidation, and calcium handling. At the functional level, mature CMs exhibited enhanced contractile kinetics, mitochondrial remodelling, polynucleation and proliferative arrest characteristic of postnatal development (Derks and Bergmann, 2020; Leone and Engel, 2019). Importantly, our analysis of alternative splicing patterns across cardiac structural genes provides independent molecular evidence that metabolic conditioning drives cells toward a postnatal splicing program. The coordinated exclusion of TNNT2, MYOM1, and FN1 exons, along with CMYA5 exon inclusion, matches splicing transitions recently characterized during human cardiac development (Gomes-Silva et al., 2025), demonstrating that maturation encompasses post-transcriptional regulatory layers beyond gene expression changes.

While it is accepted that the transition to postnatal life provides the most important shift in CM identity, function and cardiac regeneration capacity, there are many changes that occur both during embryonic development and after birth (Maroli and Braun, 2021). Thus, accurately framing the maturation phenotype to a specific *in vivo* developmental window is of paramount importance to correlate the *in vitro* generated mature CMs with actual *in vivo* CMs. In our work, the most striking evidence of postnatal-stage maturation is the initiation of polarized intercalated discs assembly. While previous studies have reported upregulation of individual ICD components, evidence for polarized ICD assembly had only been obtained in complex engineered heart tissues and not in standard 2D CM differentiations (Karbassi et al., 2020). Our observation that a network of ICD genes are upregulated after metabolic maturation, coupled with protein-level validation of polarized XIRP2 distribution, demonstrates that metabolic conditioning initiates the molecular program for ICD formation in this experimental setup. This is particularly significant because XIRP2 is among the earliest ICD proteins to polarize and is essential for redistribution of intercellular junction components from lateral membrane locations to cellular termini during postnatal development (Wang et al., 2013). To our knowledge this is the first report of polarized ICD assembly in standard 2D hPSC-CMs, an achievement that has important implications for disease modeling, as many cardiomyopathies, including arrhythmogenic cardiomyopathy and certain forms of dilated cardiomyopathy, involve ICD dysfunction (Calore et al., 2015; Zhao et al., 2019). This protocol therefore provides a framework to study ICD-related pathologies that was not previously possible with standard hPSC-CM culture conditions. Furthermore, while the heterogeneity in polarization we observed provides a metric for further protocol optimization, the achievement of polarized ICD assembly in a substantial fraction of cells represents a critical benchmark anchoring our protocol to postnatal-stage maturation.

In conclusion, in this work we resolve a fundamental question in cardiac differentiation by demonstrating that the standard WNT ON/OFF protocol generates a near-homogeneous population of left ventricular cardiomyocytes when assessed with appropriate maturation and multi-marker analysis. The apparent subtype variability that has confounded the field reflects maturation state rather than lineage diversity. Beyond clarifying this fundamental question, we provide the field with a rapid, high-fidelity platform for generating mature LV-CMs with postnatal features in a more reasonable time frame that reduces overall protocol cost and effort. As the cardiovascular research community increasingly relies on hPSC-derived cardiomyocytes for disease modeling, drug discovery, and regenerative medicine (Bois et al., 2025), the establishment of validated, chamber-specific, mature CM populations represents a critical foundation for translating these cellular models into clinically relevant insights and therapies.

## EXPERIMENTAL PROCEDURES

### hiPSC culture

The FN2.1 (Questa et al., 2016) and DESS (Castañeda et al., 2023) hiPSC lines and H1 hESCs were maintained on Geltrex™-coated dishes in StemFlex medium under standard conditions (37 °C, 5% CO₂). Cells were replated every 3–4 days using TrypLE, with 10 µM Y-27632 added for the first 24 h after plating. Cultures were routinely tested for mycoplasma contamination.

### Cardiac differentiation

CM were generated from hPSCs using the small-molecule WNT modulation protocol (Lian et al., 2012). After seeding, monolayers of 90 - 100% confluency (day 0) were treated with CHIR99021 for 24 h at indicated concentrations, followed by 5 μM IWP2 from day 3 to 5. Beating clusters appeared by day 8, after which the medium was switched to RPMI/B27 with insulin. On day 12, cells were replated and enriched by lactate selection for 6 days before downstream maturation.

### Cardiac maturation

From day 24, replated CMs were cultured in either control medium (RPMI/B27 with insulin) or maturation medium containing low-glucose DMEM supplemented with fatty acids, ascorbic acid, holo-transferrin, and GlutaMAX, with or without additional dexamethasone, T3, and a PPAR-α agonist. Mature condition cells received supplemented medium for 9 days followed by basal medium for 5 days, with media refreshed every 2 days. A detailed explanation of the maturation conditions can be found in the supplemental experimental procedures.

### Statistics

All experimental data are presented as mean and individual values with 95% confidence intervals, unless otherwise specified. Statistical analyses were performed using Python (v3.10) and SciPy library. Each experiment was independently repeated at least three times unless otherwise stated in the figure legend. When the magnitude of fold changes hindered clear visualization of differences between groups, a log2 transformation was applied to the axis scale to improve data interpretation while preserving relative relationships.

For comparisons between D23, Control, and Mature groups, a difference-in-differences test was performed using D23 as baseline to analyze the effects of time versus time plus treatment. For comparisons between CHIR concentrations in progenitor analyses, one-way ANOVA was performed followed by Tukey’s HSD post-hoc test for multiple comparisons. For comparisons between 9 μM and 12 μM CHIR after lactate enrichment protocols, unpaired two-tailed Student’s t-test was used. For comparisons between Control and Mature groups where D23 samples were not available, unpaired two-tailed Student’s t-test was performed. Statistical significance was defined as p < 0.05.

## RESOURCE AVAILABILITY

### Material, data and code availability

Further information and requests for resources and reagents should be directed to the lead contact, A.W.

### ACCESSION NUMBERS

Raw RNA-sequencing data have been deposited in the Gene Expression Omnibus (accession number GEO: GSE305566).

## ACKNOWLEDGMENTS

We thank the members of the LIAN laboratory at FLENI Neurological Hospital for their valuable contributions. We are grateful to Fundación FLENI and Fundación Perez Companc for their continuous support. This work was also supported by the following grants: PICT-2018-02672, PICT-2018-00836, PICT-2018-01722, PICT-2020-03553, and PICT-2020-03523.

## AUTHOR CONTRIBUTIONS

J.S. and A.W. conceived the experiments. J.S., J.H., D.S., S.C., A.S., and G.A. performed the experiments and analyzed data. G.S., L.M., S.M., and A.W. obtained research funding. J.S. and A.W. wrote the manuscript. A.G., S.M., and L.M. edited the manuscript.

## DECLARATION OF INTERESTS

The authors declare no conflict of interests

## Declaration of generative AI and AI-assisted technologies in the manuscript preparation process

During the preparation of this work the author(s) used Claude.AI in order to improve readability and language since they are not native english speakers. After using this tool, the authors reviewed and edited the content as needed and take full responsibility for the content of the publication.

## Notes

### Competing Interest Statement

The authors have declared no competing interest.

