## Supplementary material for "Metabolic Maturation Unveils Left Ventricular Identity in WNT ON/OFF Human Pluripotent Stem Cell-Derived Cardiomyocytes": Table S1

**Table S1. Gene Ontology (GO) analysis of upregulated genes in mature cardiomyocytes**: Enriched biological processes identified by GO analysis of genes significantly upregulated in mature cardiomyocytes. GO terms are listed with corresponding fold enrichment, raw p-value, and false discovery rate (FDR).

| **GO biological process complete** | **Fold Enrichment** | **Raw P-value** | **FDR** |
| --- | --- | --- | --- |
| cardiac muscle thin filament assembly (GO:0071691) | 21.26 | 1.04E-04 | 6.17E-03 |
| negative regulation of lipoprotein metabolic process (GO:0050748) | 17.01 | 2.34E-05 | 1.70E-03 |
| response to hydrogen sulfide (GO:1904880) | 15.95 | 4.00E-04 | 1.79E-02 |
| low-density lipoprotein particle mediated signaling (GO:0055096) | 15.95 | 4.00E-04 | 1.78E-02 |
| T-tubule organization (GO:0033292) | 14.17 | 6.76E-05 | 4.30E-03 |
| regulation of membrane depolarization during cardiac muscle cell action potential (GO:1900825) | 12.76 | 9.66E-04 | 3.42E-02 |
| lipoprotein particle mediated signaling (GO:0055095) | 12.76 | 9.66E-04 | 3.41E-02 |
| myosin filament organization (GO:0031033) | 12.76 | 9.66E-04 | 3.40E-02 |
| skeletal muscle atrophy (GO:0014732) | 12.76 | 9.66E-04 | 3.39E-02 |
| regulation of membrane depolarization during action potential (GO:0098902) | 12.15 | 1.52E-04 | 8.52E-03 |
| detection of muscle stretch (GO:0035995) | 12.15 | 1.52E-04 | 8.49E-03 |
| regulation of cardiac muscle cell membrane potential (GO:0086036) | 12.15 | 1.52E-04 | 8.46E-03 |
| striated muscle atrophy (GO:0014891) | 12.15 | 1.52E-04 | 8.43E-03 |
| positive regulation of fatty acid beta-oxidation (GO:0032000) | 10.63 | 4.71E-05 | 3.16E-03 |
| gap junction assembly (GO:0016264) | 10.63 | 2.92E-04 | 1.40E-02 |
| relaxation of cardiac muscle (GO:0055119) | 9.81 | 1.38E-05 | 1.06E-03 |
| muscle atrophy (GO:0014889) | 9.45 | 5.07E-04 | 2.09E-02 |
| skeletal myofibril assembly (GO:0014866) | 9.45 | 5.07E-04 | 2.08E-02 |
| regulation of skeletal muscle contraction (GO:0014819) | 8.95 | 1.11E-06 | 1.13E-04 |
| skeletal muscle adaptation (GO:0043501) | 8.50 | 8.13E-04 | 3.01E-02 |
| regulation of the force of heart contraction (GO:0002026) | 8.18 | 1.35E-07 | 1.75E-05 |
| relaxation of muscle (GO:0090075) | 8.10 | 2.74E-06 | 2.47E-04 |
| striated muscle adaptation (GO:0014888) | 7.87 | 2.05E-07 | 2.54E-05 |
| cardiac myofibril assembly (GO:0055003) | 7.83 | 1.53E-05 | 1.17E-03 |
| antral ovarian follicle growth (GO:0001547) | 7.73 | 1.23E-03 | 4.12E-02 |
| long-chain fatty acid import into cell (GO:0044539) | 7.73 | 1.23E-03 | 4.12E-02 |
| ovulation (GO:0030728) | 7.59 | 3.20E-04 | 1.49E-02 |
| bundle of His cell to Purkinje myocyte communication (GO:0086069) | 7.59 | 3.20E-04 | 1.48E-02 |
| atrial cardiac muscle cell to AV node cell signaling (GO:0086026) | 7.59 | 3.20E-04 | 1.48E-02 |
| atrial cardiac muscle cell action potential (GO:0086014) | 7.59 | 3.20E-04 | 1.47E-02 |
| membrane depolarization during cardiac muscle cell action potential (GO:0086012) | 7.50 | 8.44E-05 | 5.21E-03 |
| adenylate cyclase-activating adrenergic receptor signaling pathway (GO:0071880) | 7.09 | 4.61E-04 | 1.98E-02 |
| atrial cardiac muscle cell to AV node cell communication (GO:0086066) | 7.09 | 4.61E-04 | 1.98E-02 |
| cell communication by electrical coupling involved in cardiac conduction (GO:0086064) | 7.09 | 4.61E-04 | 1.97E-02 |
| cardiac muscle hypertrophy in response to stress (GO:0014898) | 7.09 | 4.61E-04 | 1.97E-02 |
| cardiac muscle adaptation (GO:0014887) | 7.09 | 4.61E-04 | 1.96E-02 |
| muscle filament sliding (GO:0030049) | 7.09 | 4.61E-04 | 1.96E-02 |
| muscle hypertrophy in response to stress (GO:0003299) | 7.09 | 4.61E-04 | 1.95E-02 |
| positive regulation of fatty acid oxidation (GO:0046321) | 7.09 | 4.61E-04 | 1.95E-02 |
| cardiac muscle contraction (GO:0060048) | 6.67 | 1.13E-15 | 1.05E-12 |
| antigen processing and presentation of peptide antigen via MHC class Ib (GO:0002428) | 6.64 | 6.44E-04 | 2.51E-02 |
| regulation of cell communication by electrical coupling (GO:0010649) | 6.64 | 6.44E-04 | 2.50E-02 |
| response to muscle stretch (GO:0035994) | 6.54 | 1.70E-05 | 1.29E-03 |
| adrenergic receptor signaling pathway (GO:0071875) | 6.47 | 6.29E-05 | 4.11E-03 |
| ventricular cardiac muscle cell action potential (GO:0086005) | 6.47 | 6.29E-05 | 4.09E-03 |
| sarcomere organization (GO:0045214) | 6.43 | 5.00E-08 | 6.99E-06 |
| cardiac muscle hypertrophy (GO:0003300) | 6.38 | 6.35E-06 | 5.20E-04 |
| positive regulation of glycogen biosynthetic process (GO:0045725) | 6.25 | 8.78E-04 | 3.20E-02 |
| heart contraction (GO:0060047) | 6.25 | 2.21E-16 | 2.98E-13 |
| actin-myosin filament sliding (GO:0033275) | 6.25 | 8.78E-04 | 3.19E-02 |
| collateral sprouting (GO:0048668) | 6.25 | 8.78E-04 | 3.18E-02 |
| myofibril assembly (GO:0030239) | 6.25 | 2.22E-11 | 5.88E-09 |
| negative regulation of reactive oxygen species biosynthetic process (GO:1903427) | 6.25 | 8.78E-04 | 3.17E-02 |
| cardiac muscle cell contraction (GO:0086003) | 6.18 | 2.68E-09 | 4.84E-07 |
| striated muscle hypertrophy (GO:0014897) | 6.17 | 8.57E-06 | 6.90E-04 |
| cell communication involved in cardiac conduction (GO:0086065) | 6.07 | 3.27E-07 | 3.85E-05 |
| antigen processing and presentation of endogenous peptide antigen via MHC class I (GO:0019885) | 6.07 | 3.15E-04 | 1.48E-02 |
| striated muscle contraction (GO:0006941) | 6.03 | 8.45E-19 | 4.17E-15 |
| regulation of cardiac conduction (GO:1903779) | 5.95 | 1.14E-04 | 6.60E-03 |
| heart process (GO:0003015) | 5.92 | 1.33E-16 | 2.19E-13 |
| regulation of fatty acid beta-oxidation (GO:0031998) | 5.91 | 1.17E-03 | 3.94E-02 |
| muscle adaptation (GO:0043500) | 5.91 | 4.21E-06 | 3.56E-04 |
| keratan sulfate proteoglycan metabolic process (GO:0042339) | 5.91 | 1.17E-03 | 3.93E-02 |
| cardiac muscle cell development (GO:0055013) | 5.89 | 6.66E-10 | 1.30E-07 |
| cardiac muscle cell action potential involved in contraction (GO:0086002) | 5.88 | 1.61E-07 | 2.04E-05 |
| cardiac muscle cell action potential (GO:0086001) | 5.86 | 6.30E-09 | 1.06E-06 |
| muscle hypertrophy (GO:0014896) | 5.80 | 1.51E-05 | 1.16E-03 |
| regulation of cardiac muscle cell contraction (GO:0086004) | 5.75 | 5.52E-06 | 4.57E-04 |
| positive regulation of D-glucose import (GO:0046326) | 5.75 | 5.52E-06 | 4.54E-04 |
| positive regulation of D-glucose transmembrane transport (GO:0010828) | 5.67 | 7.47E-07 | 8.08E-05 |
| regulation of release of sequestered calcium ion into cytosol by sarcoplasmic reticulum (GO:0010880) | 5.63 | 1.96E-05 | 1.46E-03 |
| positive regulation of reactive oxygen species biosynthetic process (GO:1903428) | 5.59 | 1.53E-03 | 4.87E-02 |
| regulation of actin filament-based movement (GO:1903115) | 5.57 | 2.63E-06 | 2.41E-04 |
| actin-mediated cell contraction (GO:0070252) | 5.57 | 2.85E-11 | 7.17E-09 |
| antigen processing and presentation of endogenous peptide antigen (GO:0002483) | 5.55 | 5.40E-04 | 2.17E-02 |
| membrane depolarization during action potential (GO:0086010) | 5.55 | 5.40E-04 | 2.17E-02 |
| regulation of cardiac muscle contraction by regulation of the release of sequestered calcium ion (GO:0010881) | 5.55 | 5.40E-04 | 2.16E-02 |
| regulation of striated muscle contraction (GO:0006942) | 5.53 | 5.47E-13 | 2.13E-10 |
| positive regulation of lipid catabolic process (GO:0050996) | 5.51 | 1.93E-04 | 1.00E-02 |
| cardiac cell development (GO:0055006) | 5.39 | 3.18E-09 | 5.68E-07 |
| regulation of cardiac muscle contraction by calcium ion signaling (GO:0010882) | 5.32 | 8.92E-05 | 5.42E-03 |
| cardiac conduction (GO:0061337) | 5.24 | 1.39E-08 | 2.10E-06 |
| cell-cell signaling involved in cardiac conduction (GO:0086019) | 5.10 | 8.74E-04 | 3.20E-02 |
| negative regulation of cardiac muscle cell apoptotic process (GO:0010667) | 5.10 | 8.74E-04 | 3.19E-02 |
| actin filament-based movement (GO:0030048) | 4.97 | 1.98E-11 | 5.44E-09 |
| regulation of reactive oxygen species biosynthetic process (GO:1903426) | 4.94 | 2.35E-05 | 1.70E-03 |
| muscle cell cellular homeostasis (GO:0046716) | 4.91 | 1.09E-03 | 3.76E-02 |
| positive regulation of ATP metabolic process (GO:1903580) | 4.91 | 6.42E-05 | 4.16E-03 |
| cardiac muscle cell differentiation (GO:0055007) | 4.87 | 4.74E-10 | 9.63E-08 |
| skeletal muscle contraction (GO:0003009) | 4.86 | 1.76E-04 | 9.42E-03 |
| platelet-derived growth factor receptor signaling pathway (GO:0048008) | 4.86 | 1.76E-04 | 9.39E-03 |
| regulation of response to oxidative stress (GO:1902882) | 4.86 | 1.76E-04 | 9.35E-03 |
| striated muscle cell development (GO:0055002) | 4.76 | 1.11E-13 | 5.67E-11 |
| regulation of membrane repolarization (GO:0060306) | 4.72 | 2.17E-04 | 1.11E-02 |
| regulation of cardiac muscle contraction (GO:0055117) | 4.72 | 3.00E-08 | 4.31E-06 |
| regulation of heart rate (GO:0002027) | 4.72 | 1.53E-10 | 3.29E-08 |
| regulation of membrane depolarization (GO:0003254) | 4.68 | 1.63E-05 | 1.24E-03 |
| positive regulation of purine nucleotide metabolic process (GO:1900544) | 4.67 | 9.76E-05 | 5.88E-03 |
| positive regulation of nucleotide metabolic process (GO:0045981) | 4.67 | 9.76E-05 | 5.85E-03 |
| regulation of heart contraction (GO:0008016) | 4.63 | 1.35E-18 | 4.98E-15 |
| regulation of cardiac muscle cell apoptotic process (GO:0010665) | 4.60 | 2.66E-04 | 1.29E-02 |
| actomyosin structure organization (GO:0031032) | 4.57 | 5.91E-11 | 1.41E-08 |
| cell-substrate junction organization (GO:0150115) | 4.56 | 1.19E-04 | 6.86E-03 |
| regulation of macrophage derived foam cell differentiation (GO:0010743) | 4.51 | 7.27E-04 | 2.76E-02 |
| heterotypic cell-cell adhesion (GO:0034113) | 4.48 | 3.23E-04 | 1.48E-02 |
| muscle contraction (GO:0006936) | 4.46 | 1.03E-19 | 7.63E-16 |
| muscle cell development (GO:0055001) | 4.45 | 2.90E-14 | 1.65E-11 |
| regulation of fatty acid oxidation (GO:0046320) | 4.38 | 8.79E-04 | 3.17E-02 |
| regulation of calcineurin-NFAT signaling cascade (GO:0070884) | 4.36 | 3.90E-04 | 1.74E-02 |
| insulin receptor signaling pathway (GO:0008286) | 4.31 | 3.25E-06 | 2.87E-04 |
| regulation of sodium ion transport (GO:0002028) | 4.31 | 1.48E-06 | 1.44E-04 |
| regulation of muscle contraction (GO:0006937) | 4.25 | 6.01E-13 | 2.28E-10 |
| antigen processing and presentation of peptide antigen via MHC class I (GO:0002474) | 4.25 | 1.05E-03 | 3.65E-02 |
| cell-substrate junction assembly (GO:0007044) | 4.25 | 4.67E-04 | 1.96E-02 |
| regulation of calcineurin-mediated signaling (GO:0106056) | 4.25 | 4.67E-04 | 1.95E-02 |
| regulation of striated muscle cell apoptotic process (GO:0010662) | 4.25 | 4.67E-04 | 1.94E-02 |
| muscle system process (GO:0003012) | 4.24 | 1.96E-21 | 2.91E-17 |
| cardiac muscle tissue development (GO:0048738) | 4.23 | 1.42E-14 | 8.76E-12 |
| regulation of muscle adaptation (GO:0043502) | 4.20 | 9.58E-07 | 1.01E-04 |
| response to ischemia (GO:0002931) | 4.11 | 2.69E-05 | 1.90E-03 |
| musculoskeletal movement (GO:0050881) | 4.07 | 2.95E-04 | 1.41E-02 |
| regulation of lipid storage (GO:0010883) | 4.07 | 2.95E-04 | 1.40E-02 |
| negative regulation of JNK cascade (GO:0046329) | 4.02 | 1.49E-03 | 4.79E-02 |
| cellular response to amino acid starvation (GO:0034198) | 4.01 | 1.56E-04 | 8.50E-03 |
| regulation of muscle system process (GO:0090257) | 4.00 | 1.15E-15 | 1.00E-12 |
| cellular anatomical entity morphogenesis (GO:0032989) | 3.96 | 6.88E-09 | 1.14E-06 |
| cellular component assembly involved in morphogenesis (GO:0010927) | 3.96 | 6.88E-09 | 1.13E-06 |
| regulation of heart rate by cardiac conduction (GO:0086091) | 3.96 | 7.78E-04 | 2.90E-02 |
| cardiocyte differentiation (GO:0035051) | 3.91 | 1.76E-08 | 2.63E-06 |
| multicellular organismal movement (GO:0050879) | 3.90 | 4.08E-04 | 1.81E-02 |
| regulation of blood circulation (GO:1903522) | 3.87 | 5.35E-16 | 5.66E-13 |
| regulation of action potential (GO:0098900) | 3.83 | 1.14E-04 | 6.67E-03 |
| response to fatty acid (GO:0070542) | 3.83 | 1.14E-04 | 6.65E-03 |
| cardiac muscle tissue morphogenesis (GO:0055008) | 3.83 | 1.14E-04 | 6.62E-03 |
| response to amino acid starvation (GO:1990928) | 3.80 | 2.51E-04 | 1.26E-02 |
| regulation of purine nucleotide catabolic process (GO:0033121) | 3.80 | 2.51E-04 | 1.26E-02 |
| regulation of glycolytic process (GO:0006110) | 3.80 | 2.51E-04 | 1.25E-02 |
| mitophagy (GO:0000423) | 3.80 | 2.51E-04 | 1.25E-02 |
| regulation of nucleotide catabolic process (GO:0030811) | 3.80 | 2.51E-04 | 1.24E-02 |
| regulation of D-glucose import (GO:0046324) | 3.80 | 2.51E-04 | 1.24E-02 |
| striated muscle tissue development (GO:0014706) | 3.72 | 1.47E-18 | 4.35E-15 |
| vascular transport (GO:0010232) | 3.67 | 1.20E-05 | 9.44E-04 |
| transport across blood-brain barrier (GO:0150104) | 3.67 | 1.20E-05 | 9.39E-04 |
| negative regulation of actin filament polymerization (GO:0030837) | 3.61 | 7.45E-04 | 2.81E-02 |
| muscle tissue development (GO:0060537) | 3.58 | 5.00E-18 | 1.06E-14 |
| action potential (GO:0001508) | 3.57 | 1.94E-07 | 2.41E-05 |
| striated muscle cell differentiation (GO:0051146) | 3.56 | 8.41E-12 | 2.54E-09 |
| regulation of carbohydrate catabolic process (GO:0043470) | 3.49 | 2.70E-04 | 1.31E-02 |
| regulation of vascular associated smooth muscle cell proliferation (GO:1904705) | 3.49 | 5.14E-04 | 2.09E-02 |
| autophagy of mitochondrion (GO:0000422) | 3.45 | 1.62E-04 | 8.79E-03 |
| ventricular cardiac muscle tissue development (GO:0003229) | 3.43 | 5.87E-04 | 2.31E-02 |
| regulation of actin filament depolymerization (GO:0030834) | 3.42 | 1.12E-03 | 3.82E-02 |
| skeletal muscle tissue development (GO:0007519) | 3.38 | 1.60E-07 | 2.04E-05 |
| regulation of D-glucose transmembrane transport (GO:0010827) | 3.36 | 2.11E-04 | 1.08E-02 |
| negative regulation of supramolecular fiber organization (GO:1902904) | 3.32 | 1.25E-07 | 1.65E-05 |
| regulation of cardiac muscle hypertrophy (GO:0010611) | 3.30 | 1.45E-03 | 4.68E-02 |
| regulation of muscle cell apoptotic process (GO:0010660) | 3.29 | 4.53E-04 | 1.96E-02 |
| negative regulation of cytoskeleton organization (GO:0051494) | 3.27 | 2.98E-07 | 3.56E-05 |
| regulation of ATP metabolic process (GO:1903578) | 3.27 | 2.71E-04 | 1.31E-02 |
| negative regulation of protein depolymerization (GO:1901880) | 3.25 | 5.13E-04 | 2.09E-02 |
| steroid hormone receptor signaling pathway (GO:0043401) | 3.23 | 3.06E-04 | 1.45E-02 |
| nuclear receptor-mediated steroid hormone signaling pathway (GO:0030518) | 3.23 | 3.06E-04 | 1.44E-02 |
| skeletal muscle organ development (GO:0060538) | 3.22 | 2.29E-07 | 2.78E-05 |
| nuclear receptor-mediated signaling pathway (GO:0141193) | 3.21 | 3.50E-05 | 2.43E-03 |
| negative regulation of protein polymerization (GO:0032272) | 3.17 | 1.10E-03 | 3.76E-02 |
| muscle tissue morphogenesis (GO:0060415) | 3.16 | 6.51E-04 | 2.51E-02 |
| muscle cell differentiation (GO:0042692) | 3.15 | 2.29E-11 | 5.96E-09 |
| muscle organ morphogenesis (GO:0048644) | 3.11 | 4.35E-04 | 1.89E-02 |
| regulation of protein depolymerization (GO:1901879) | 3.10 | 1.55E-04 | 8.53E-03 |
| muscle organ development (GO:0007517) | 3.08 | 5.92E-12 | 1.83E-09 |
| cellular response to insulin stimulus (GO:0032869) | 3.08 | 2.62E-06 | 2.41E-04 |
| regulation of potassium ion transmembrane transport (GO:1901379) | 3.04 | 1.54E-03 | 4.90E-02 |
| cellular response to hexose stimulus (GO:0071331) | 2.96 | 1.14E-03 | 3.85E-02 |
| regulation of system process (GO:0044057) | 2.95 | 7.33E-18 | 1.36E-14 |
| regulation of purine nucleotide metabolic process (GO:1900542) | 2.93 | 7.51E-04 | 2.82E-02 |
| energy homeostasis (GO:0097009) | 2.93 | 7.51E-04 | 2.82E-02 |
| regulation of release of sequestered calcium ion into cytosol (GO:0051279) | 2.92 | 1.26E-03 | 4.22E-02 |
| positive regulation of carbohydrate metabolic process (GO:0045913) | 2.92 | 1.26E-03 | 4.21E-02 |
| regulation of nucleotide metabolic process (GO:0006140) | 2.90 | 8.33E-04 | 3.07E-02 |
| circulatory system process (GO:0003013) | 2.89 | 6.31E-16 | 6.23E-13 |
| muscle structure development (GO:0061061) | 2.89 | 4.16E-16 | 4.74E-13 |
| cellular response to monosaccharide stimulus (GO:0071326) | 2.89 | 1.40E-03 | 4.57E-02 |
| positive regulation of macroautophagy (GO:0016239) | 2.88 | 5.50E-04 | 2.19E-02 |
| regulation of protein-containing complex disassembly (GO:0043244) | 2.86 | 5.75E-05 | 3.79E-03 |
| positive regulation of smooth muscle cell proliferation (GO:0048661) | 2.85 | 1.55E-03 | 4.91E-02 |
| negative regulation of protein-containing complex disassembly (GO:0043242) | 2.85 | 1.55E-03 | 4.90E-02 |
| cellular response to carbohydrate stimulus (GO:0071322) | 2.83 | 1.02E-03 | 3.55E-02 |
| cellular response to starvation (GO:0009267) | 2.83 | 2.73E-06 | 2.48E-04 |
| regulation of generation of precursor metabolites and energy (GO:0043467) | 2.79 | 7.71E-05 | 4.86E-03 |
| cellular response to peptide hormone stimulus (GO:0071375) | 2.72 | 1.84E-07 | 2.31E-05 |
| positive regulation of transmembrane transport (GO:0034764) | 2.71 | 8.73E-06 | 6.99E-04 |
| blood circulation (GO:0008015) | 2.71 | 1.69E-11 | 4.74E-09 |
| regulation of reactive oxygen species metabolic process (GO:2000377) | 2.69 | 1.23E-04 | 7.07E-03 |
| cellular response to nutrient levels (GO:0031669) | 2.68 | 5.98E-07 | 6.67E-05 |
| response to starvation (GO:0042594) | 2.67 | 2.23E-06 | 2.08E-04 |
| positive regulation of cell-substrate adhesion (GO:0010811) | 2.66 | 3.38E-04 | 1.54E-02 |
| positive regulation of small molecule metabolic process (GO:0062013) | 2.64 | 1.07E-04 | 6.29E-03 |
| response to insulin (GO:0032868) | 2.63 | 4.37E-06 | 3.68E-04 |
| actin filament-based process (GO:0030029) | 2.62 | 1.33E-14 | 8.54E-12 |
| regulation of carbohydrate metabolic process (GO:0006109) | 2.60 | 4.02E-05 | 2.72E-03 |
| regulation of sequestering of calcium ion (GO:0051282) | 2.60 | 4.39E-04 | 1.91E-02 |
| negative regulation of protein-containing complex assembly (GO:0031333) | 2.58 | 3.16E-04 | 1.48E-02 |
| regulation of actin filament-based process (GO:0032970) | 2.58 | 1.90E-09 | 3.48E-07 |
| fat cell differentiation (GO:0045444) | 2.56 | 5.20E-04 | 2.11E-02 |
| hormone-mediated signaling pathway (GO:0009755) | 2.55 | 2.47E-04 | 1.25E-02 |
| regulation of smooth muscle cell proliferation (GO:0048660) | 2.50 | 6.67E-04 | 2.56E-02 |
| cellular response to steroid hormone stimulus (GO:0071383) | 2.48 | 3.43E-04 | 1.55E-02 |
| lysosomal transport (GO:0007041) | 2.47 | 1.09E-03 | 3.76E-02 |
| calcium-mediated signaling (GO:0019722) | 2.47 | 1.77E-04 | 9.35E-03 |
| small GTPase-mediated signal transduction (GO:0007264) | 2.47 | 3.47E-06 | 3.00E-04 |
| response to monosaccharide (GO:0034284) | 2.46 | 5.60E-04 | 2.23E-02 |
| actin filament organization (GO:0007015) | 2.46 | 4.16E-06 | 3.54E-04 |
| positive regulation of protein kinase activity (GO:0045860) | 2.45 | 8.46E-04 | 3.10E-02 |
| signal transduction in response to DNA damage (GO:0042770) | 2.44 | 6.06E-04 | 2.37E-02 |
| response to hexose (GO:0009746) | 2.43 | 9.15E-04 | 3.28E-02 |
| regulation of cell-substrate adhesion (GO:0010810) | 2.43 | 6.43E-05 | 4.14E-03 |
| regulation of small GTPase mediated signal transduction (GO:0051056) | 2.40 | 1.88E-06 | 1.78E-04 |
| regulation of muscle cell differentiation (GO:0051147) | 2.38 | 1.15E-03 | 3.88E-02 |
| regulation of membrane potential (GO:0042391) | 2.38 | 1.03E-08 | 1.62E-06 |
| cellular response to hormone stimulus (GO:0032870) | 2.38 | 5.86E-10 | 1.16E-07 |
| heart development (GO:0007507) | 2.37 | 1.44E-10 | 3.15E-08 |
| supramolecular fiber organization (GO:0097435) | 2.37 | 3.60E-11 | 8.90E-09 |
| regulation of metal ion transport (GO:0010959) | 2.37 | 3.13E-07 | 3.71E-05 |
| positive regulation of autophagy (GO:0010508) | 2.36 | 1.03E-03 | 3.57E-02 |
| positive regulation of phosphorus metabolic process (GO:0010562) | 2.36 | 1.37E-06 | 1.37E-04 |
| positive regulation of phosphate metabolic process (GO:0045937) | 2.36 | 1.37E-06 | 1.36E-04 |
| glucose homeostasis (GO:0042593) | 2.35 | 1.71E-04 | 9.22E-03 |
| monoatomic anion transport (GO:0006820) | 2.35 | 1.06E-03 | 3.65E-02 |
| regulation of TOR signaling (GO:0032006) | 2.35 | 1.53E-03 | 4.89E-02 |
| regulation of transmembrane transport (GO:0034762) | 2.35 | 1.09E-07 | 1.49E-05 |
| response to peptide hormone (GO:0043434) | 2.34 | 4.64E-07 | 5.33E-05 |
| carbohydrate homeostasis (GO:0033500) | 2.34 | 1.79E-04 | 9.43E-03 |
| negative regulation of organelle organization (GO:0010639) | 2.33 | 1.66E-06 | 1.61E-04 |
| regulation of monoatomic cation transmembrane transport (GO:1904062) | 2.33 | 1.75E-05 | 1.32E-03 |
| vascular process in circulatory system (GO:0003018) | 2.32 | 1.86E-05 | 1.39E-03 |
| cellular response to metal ion (GO:0071248) | 2.31 | 5.69E-04 | 2.25E-02 |
| intracellular receptor signaling pathway (GO:0030522) | 2.29 | 3.08E-04 | 1.45E-02 |
| regulation of lipid biosynthetic process (GO:0046890) | 2.29 | 6.21E-04 | 2.43E-02 |
| actin cytoskeleton organization (GO:0030036) | 2.27 | 6.21E-09 | 1.06E-06 |
| regulation of monoatomic ion transport (GO:0043269) | 2.27 | 2.18E-07 | 2.67E-05 |
| neuromuscular process (GO:0050905) | 2.26 | 1.32E-03 | 4.38E-02 |
| import across plasma membrane (GO:0098739) | 2.26 | 1.32E-03 | 4.37E-02 |
| cell surface receptor protein tyrosine kinase signaling pathway (GO:0007169) | 2.26 | 2.31E-07 | 2.78E-05 |
| regulation of monoatomic ion transmembrane transport (GO:0034765) | 2.25 | 2.33E-05 | 1.70E-03 |
| regulation of blood pressure (GO:0008217) | 2.24 | 5.35E-04 | 2.16E-02 |
| regulation of actin cytoskeleton organization (GO:0032956) | 2.23 | 5.11E-06 | 4.25E-04 |
| glycerolipid biosynthetic process (GO:0045017) | 2.23 | 4.23E-04 | 1.85E-02 |
| response to carbohydrate (GO:0009743) | 2.22 | 1.52E-03 | 4.86E-02 |
| positive regulation of monoatomic ion transport (GO:0043270) | 2.21 | 8.39E-04 | 3.08E-02 |
| intracellular signaling cassette (GO:0141124) | 2.21 | 3.15E-12 | 1.06E-09 |
| regulation of supramolecular fiber organization (GO:1902903) | 2.20 | 3.03E-06 | 2.69E-04 |
| circulatory system development (GO:0072359) | 2.19 | 2.50E-13 | 1.16E-10 |
| connective tissue development (GO:0061448) | 2.18 | 3.22E-04 | 1.48E-02 |
| anatomical structure formation involved in morphogenesis (GO:0048646) | 2.17 | 1.51E-13 | 7.44E-11 |
| glycerophospholipid biosynthetic process (GO:0046474) | 2.15 | 1.54E-03 | 4.91E-02 |
| response to steroid hormone (GO:0048545) | 2.13 | 1.59E-04 | 8.61E-03 |
| cellular response to nitrogen compound (GO:1901699) | 2.12 | 5.82E-08 | 7.98E-06 |
| response to hormone (GO:0009725) | 2.11 | 3.54E-10 | 7.28E-08 |
| inorganic ion homeostasis (GO:0098771) | 2.11 | 3.60E-06 | 3.10E-04 |
| phosphorylation (GO:0016310) | 2.10 | 1.13E-05 | 8.94E-04 |
| heart morphogenesis (GO:0003007) | 2.10 | 3.41E-04 | 1.55E-02 |
| response to nutrient levels (GO:0031667) | 2.09 | 1.47E-06 | 1.44E-04 |
| regulation of cytoskeleton organization (GO:0051493) | 2.08 | 5.11E-07 | 5.77E-05 |
| positive regulation of catabolic process (GO:0009896) | 2.07 | 6.84E-07 | 7.56E-05 |
| regulation of neuron apoptotic process (GO:0043523) | 2.07 | 5.48E-04 | 2.19E-02 |
| regulation of actin filament organization (GO:0110053) | 2.05 | 6.51E-04 | 2.52E-02 |
| positive regulation of protein phosphorylation (GO:0001934) | 2.05 | 8.00E-04 | 2.97E-02 |
| cellular homeostasis (GO:0019725) | 2.03 | 4.13E-08 | 5.89E-06 |
| regulation of autophagy (GO:0010506) | 2.03 | 8.36E-05 | 5.18E-03 |
| positive regulation of cell migration (GO:0030335) | 2.00 | 2.00E-06 | 1.88E-04 |
| extracellular matrix organization (GO:0030198) | 1.99 | 6.63E-04 | 2.55E-02 |
| intracellular signal transduction (GO:0035556) | 1.98 | 3.29E-18 | 8.12E-15 |
| positive regulation of cell motility (GO:2000147) | 1.98 | 1.92E-06 | 1.81E-04 |
| extracellular structure organization (GO:0043062) | 1.98 | 6.90E-04 | 2.64E-02 |
| regulation of phosphatidylinositol 3-kinase/protein kinase B signal transduction (GO:0051896) | 1.97 | 9.19E-04 | 3.29E-02 |
| external encapsulating structure organization (GO:0045229) | 1.97 | 7.19E-04 | 2.74E-02 |
| enzyme-linked receptor protein signaling pathway (GO:0007167) | 1.97 | 8.00E-07 | 8.58E-05 |
| intracellular chemical homeostasis (GO:0055082) | 1.97 | 2.89E-06 | 2.60E-04 |
| positive regulation of locomotion (GO:0040017) | 1.97 | 1.73E-06 | 1.67E-04 |
| positive regulation of phosphorylation (GO:0042327) | 1.95 | 1.32E-03 | 4.36E-02 |
| monoatomic cation homeostasis (GO:0055080) | 1.95 | 1.16E-05 | 9.18E-04 |
| monoatomic ion homeostasis (GO:0050801) | 1.95 | 1.29E-05 | 1.00E-03 |
| protein phosphorylation (GO:0006468) | 1.95 | 3.03E-04 | 1.44E-02 |
| response to oxidative stress (GO:0006979) | 1.94 | 2.65E-04 | 1.29E-02 |
| angiogenesis (GO:0001525) | 1.94 | 4.80E-04 | 1.99E-02 |
| positive regulation of MAPK cascade (GO:0043410) | 1.91 | 7.29E-05 | 4.61E-03 |
| chemical homeostasis (GO:0048878) | 1.90 | 2.31E-08 | 3.40E-06 |
| cellular response to oxygen-containing compound (GO:1901701) | 1.90 | 1.56E-09 | 2.89E-07 |
| regulation of cellular response to growth factor stimulus (GO:0090287) | 1.90 | 1.28E-03 | 4.25E-02 |
| response to decreased oxygen levels (GO:0036293) | 1.89 | 1.45E-03 | 4.69E-02 |
| response to metal ion (GO:0010038) | 1.89 | 7.41E-04 | 2.80E-02 |
| regulation of small molecule metabolic process (GO:0062012) | 1.89 | 9.77E-04 | 3.41E-02 |
| regulation of cell migration (GO:0030334) | 1.89 | 1.82E-08 | 2.70E-06 |
| regulation of MAPK cascade (GO:0043408) | 1.89 | 3.34E-06 | 2.93E-04 |
| import into cell (GO:0098657) | 1.88 | 1.76E-06 | 1.68E-04 |
| blood vessel morphogenesis (GO:0048514) | 1.88 | 1.55E-04 | 8.51E-03 |
| positive regulation of intracellular signal transduction (GO:1902533) | 1.87 | 1.28E-09 | 2.40E-07 |
| regulation of lipid metabolic process (GO:0019216) | 1.86 | 1.43E-03 | 4.63E-02 |
| regulation of cell motility (GO:2000145) | 1.86 | 1.30E-08 | 2.00E-06 |
| positive regulation of transport (GO:0051050) | 1.84 | 4.49E-07 | 5.20E-05 |
| membraneless organelle assembly (GO:0140694) | 1.84 | 1.40E-03 | 4.57E-02 |
| intracellular monoatomic ion homeostasis (GO:0006873) | 1.84 | 3.48E-04 | 1.57E-02 |
| cytoskeleton organization (GO:0007010) | 1.83 | 5.19E-10 | 1.04E-07 |
| regulation of locomotion (GO:0040012) | 1.83 | 2.38E-08 | 3.45E-06 |
| negative regulation of cellular component organization (GO:0051129) | 1.82 | 6.75E-06 | 5.50E-04 |
| regulation of apoptotic signaling pathway (GO:2001233) | 1.82 | 9.04E-04 | 3.25E-02 |
| vasculature development (GO:0001944) | 1.82 | 6.64E-05 | 4.26E-03 |
| tissue development (GO:0009888) | 1.82 | 1.61E-13 | 7.67E-11 |
| regulation of epithelial cell proliferation (GO:0050678) | 1.82 | 1.52E-03 | 4.87E-02 |
| response to nitrogen compound (GO:1901698) | 1.81 | 1.39E-07 | 1.79E-05 |
| positive regulation of programmed cell death (GO:0043068) | 1.80 | 1.19E-04 | 6.86E-03 |
| response to endogenous stimulus (GO:0009719) | 1.80 | 6.19E-09 | 1.07E-06 |
| anatomical structure morphogenesis (GO:0009653) | 1.80 | 2.40E-16 | 2.96E-13 |
| regulation of phosphate metabolic process (GO:0019220) | 1.80 | 5.35E-05 | 3.57E-03 |
| regulation of phosphorus metabolic process (GO:0051174) | 1.79 | 5.49E-05 | 3.65E-03 |
| response to wounding (GO:0009611) | 1.79 | 6.33E-04 | 2.47E-02 |
| positive regulation of apoptotic process (GO:0043065) | 1.78 | 2.20E-04 | 1.12E-02 |
| blood vessel development (GO:0001568) | 1.78 | 1.75E-04 | 9.41E-03 |
| regulation of intracellular signal transduction (GO:1902531) | 1.78 | 5.41E-14 | 2.86E-11 |
| intracellular monoatomic cation homeostasis (GO:0030003) | 1.78 | 7.95E-04 | 2.96E-02 |
| cellular response to endogenous stimulus (GO:0071495) | 1.77 | 1.10E-06 | 1.13E-04 |
| regulation of protein-containing complex assembly (GO:0043254) | 1.77 | 1.13E-03 | 3.85E-02 |
| positive regulation of signal transduction (GO:0009967) | 1.76 | 1.07E-10 | 2.40E-08 |
| regulation of anatomical structure size (GO:0090066) | 1.75 | 4.14E-04 | 1.82E-02 |
| cell junction organization (GO:0034330) | 1.75 | 1.87E-04 | 9.77E-03 |
| tube morphogenesis (GO:0035239) | 1.73 | 5.54E-05 | 3.66E-03 |
| protein ubiquitination (GO:0016567) | 1.72 | 2.44E-04 | 1.24E-02 |
| positive regulation of cell communication (GO:0010647) | 1.72 | 6.42E-11 | 1.51E-08 |
| regulation of localization (GO:0032879) | 1.71 | 5.51E-12 | 1.74E-09 |
| homeostatic process (GO:0042592) | 1.71 | 5.59E-09 | 9.75E-07 |
| positive regulation of signaling (GO:0023056) | 1.70 | 1.42E-10 | 3.13E-08 |
| response to oxygen-containing compound (GO:1901700) | 1.70 | 4.70E-09 | 8.30E-07 |
| lipid biosynthetic process (GO:0008610) | 1.70 | 4.82E-04 | 1.99E-02 |
| regulation of transport (GO:0051049) | 1.68 | 7.91E-09 | 1.29E-06 |
| regulation of anatomical structure morphogenesis (GO:0022603) | 1.67 | 3.94E-05 | 2.71E-03 |
| endocytosis (GO:0006897) | 1.67 | 1.37E-03 | 4.51E-02 |
| negative regulation of intracellular signal transduction (GO:1902532) | 1.67 | 1.55E-04 | 8.49E-03 |
| response to abiotic stimulus (GO:0009628) | 1.67 | 2.26E-06 | 2.10E-04 |
| behavior (GO:0007610) | 1.66 | 4.65E-04 | 1.95E-02 |
| cellular response to lipid (GO:0071396) | 1.65 | 1.34E-03 | 4.40E-02 |
| tissue morphogenesis (GO:0048729) | 1.65 | 9.64E-04 | 3.42E-02 |
| regulation of organelle organization (GO:0033043) | 1.64 | 3.45E-06 | 3.01E-04 |
| positive regulation of protein metabolic process (GO:0051247) | 1.64 | 1.30E-04 | 7.38E-03 |
| animal organ morphogenesis (GO:0009887) | 1.64 | 2.02E-05 | 1.49E-03 |
| negative regulation of transcription by RNA polymerase II (GO:0000122) | 1.63 | 3.17E-05 | 2.22E-03 |
| cell population proliferation (GO:0008283) | 1.62 | 3.86E-04 | 1.73E-02 |
| phosphorus metabolic process (GO:0006793) | 1.62 | 5.11E-07 | 5.73E-05 |
| regulation of cell adhesion (GO:0030155) | 1.62 | 2.26E-04 | 1.15E-02 |
| protein modification by small protein conjugation (GO:0032446) | 1.62 | 7.77E-04 | 2.91E-02 |
| negative regulation of signal transduction (GO:0009968) | 1.61 | 9.83E-07 | 1.02E-04 |
| negative regulation of programmed cell death (GO:0043069) | 1.61 | 8.62E-05 | 5.28E-03 |
| metal ion transport (GO:0030001) | 1.61 | 9.83E-04 | 3.43E-02 |
| phosphate-containing compound metabolic process (GO:0006796) | 1.61 | 7.29E-07 | 8.00E-05 |
| regulation of signal transduction (GO:0009966) | 1.60 | 2.73E-14 | 1.62E-11 |
| cell migration (GO:0016477) | 1.60 | 1.00E-04 | 5.98E-03 |
| negative regulation of cell population proliferation (GO:0008285) | 1.60 | 7.35E-04 | 2.79E-02 |
| tube development (GO:0035295) | 1.60 | 9.36E-05 | 5.66E-03 |
| negative regulation of cell differentiation (GO:0045596) | 1.60 | 1.13E-03 | 3.84E-02 |
| cellular response to chemical stimulus (GO:0070887) | 1.60 | 8.46E-09 | 1.36E-06 |
| regulation of multicellular organismal process (GO:0051239) | 1.60 | 2.54E-13 | 1.14E-10 |
| negative regulation of cell communication (GO:0010648) | 1.59 | 8.00E-07 | 8.53E-05 |
| negative regulation of signaling (GO:0023057) | 1.59 | 8.13E-07 | 8.61E-05 |
| response to lipid (GO:0033993) | 1.59 | 2.57E-04 | 1.26E-02 |
| regulation of catabolic process (GO:0009894) | 1.58 | 5.98E-05 | 3.92E-03 |
| regulation of cellular component biogenesis (GO:0044087) | 1.58 | 7.71E-05 | 4.84E-03 |
| regulation of cell communication (GO:0010646) | 1.57 | 5.17E-15 | 3.65E-12 |
| negative regulation of developmental process (GO:0051093) | 1.57 | 1.89E-04 | 9.82E-03 |
| monoatomic ion transport (GO:0006811) | 1.57 | 1.06E-04 | 6.29E-03 |
| regulation of cell differentiation (GO:0045595) | 1.56 | 1.47E-06 | 1.45E-04 |
| system process (GO:0003008) | 1.56 | 9.73E-09 | 1.55E-06 |
| negative regulation of apoptotic process (GO:0043066) | 1.56 | 4.54E-04 | 1.96E-02 |
| regulation of cell population proliferation (GO:0042127) | 1.55 | 1.13E-06 | 1.14E-04 |
| regulation of signaling (GO:0023051) | 1.55 | 4.26E-14 | 2.34E-11 |
| negative regulation of response to stimulus (GO:0048585) | 1.55 | 7.38E-07 | 8.04E-05 |
| regulation of programmed cell death (GO:0043067) | 1.55 | 4.89E-06 | 4.09E-04 |
| animal organ development (GO:0048513) | 1.54 | 3.63E-11 | 8.83E-09 |
| cell development (GO:0048468) | 1.54 | 1.04E-08 | 1.62E-06 |
| positive regulation of response to stimulus (GO:0048584) | 1.54 | 1.33E-08 | 2.02E-06 |
| monoatomic ion transmembrane transport (GO:0034220) | 1.53 | 9.23E-04 | 3.29E-02 |
| negative regulation of RNA biosynthetic process (GO:1902679) | 1.53 | 3.51E-05 | 2.43E-03 |
| monoatomic cation transport (GO:0006812) | 1.52 | 1.11E-03 | 3.78E-02 |
| negative regulation of DNA-templated transcription (GO:0045892) | 1.52 | 4.47E-05 | 3.01E-03 |
| organophosphate metabolic process (GO:0019637) | 1.52 | 4.61E-04 | 1.94E-02 |
| system development (GO:0048731) | 1.52 | 5.21E-13 | 2.09E-10 |
| cell death (GO:0008219) | 1.51 | 2.53E-04 | 1.24E-02 |
| organelle assembly (GO:0070925) | 1.51 | 9.40E-04 | 3.35E-02 |
| regulation of protein metabolic process (GO:0051246) | 1.50 | 2.14E-05 | 1.57E-03 |
| regulation of cellular localization (GO:0060341) | 1.50 | 5.97E-04 | 2.35E-02 |
| regulation of apoptotic process (GO:0042981) | 1.49 | 3.84E-05 | 2.64E-03 |
| multicellular organism development (GO:0007275) | 1.49 | 2.74E-13 | 1.19E-10 |
| regulation of cellular component organization (GO:0051128) | 1.49 | 5.51E-08 | 7.62E-06 |
| positive regulation of cell population proliferation (GO:0008284) | 1.49 | 1.45E-03 | 4.68E-02 |
| regulation of multicellular organismal development (GO:2000026) | 1.48 | 8.34E-05 | 5.19E-03 |
| regulation of response to stimulus (GO:0048583) | 1.48 | 3.02E-13 | 1.28E-10 |
| regulation of developmental process (GO:0050793) | 1.48 | 1.11E-07 | 1.50E-05 |
| negative regulation of multicellular organismal process (GO:0051241) | 1.48 | 5.07E-04 | 2.08E-02 |
| plasma membrane bounded cell projection organization (GO:0120036) | 1.48 | 4.03E-04 | 1.79E-02 |
| cell motility (GO:0048870) | 1.47 | 4.77E-04 | 1.98E-02 |
| cell projection organization (GO:0030030) | 1.47 | 4.09E-04 | 1.80E-02 |
| programmed cell death (GO:0012501) | 1.46 | 9.71E-04 | 3.40E-02 |
| apoptotic process (GO:0006915) | 1.45 | 1.35E-03 | 4.44E-02 |
| cellular developmental process (GO:0048869) | 1.44 | 1.74E-10 | 3.69E-08 |
| negative regulation of RNA metabolic process (GO:0051253) | 1.44 | 2.58E-04 | 1.26E-02 |
| lipid metabolic process (GO:0006629) | 1.44 | 9.59E-04 | 3.41E-02 |
| cell differentiation (GO:0030154) | 1.44 | 2.99E-10 | 6.23E-08 |
| anatomical structure development (GO:0048856) | 1.43 | 6.22E-15 | 4.19E-12 |
| neurogenesis (GO:0022008) | 1.43 | 4.17E-04 | 1.83E-02 |
| developmental process (GO:0032502) | 1.41 | 2.76E-15 | 2.16E-12 |
| protein modification process (GO:0036211) | 1.41 | 1.83E-04 | 9.59E-03 |
| negative regulation of biological process (GO:0048519) | 1.39 | 1.29E-12 | 4.65E-10 |
| nervous system development (GO:0007399) | 1.39 | 2.58E-05 | 1.86E-03 |
| cell communication (GO:0007154) | 1.39 | 1.63E-12 | 5.75E-10 |
| regulation of biological quality (GO:0065008) | 1.39 | 1.13E-06 | 1.15E-04 |
| signaling (GO:0023052) | 1.38 | 3.25E-12 | 1.07E-09 |
| negative regulation of cellular process (GO:0048523) | 1.38 | 2.13E-11 | 5.73E-09 |
| negative regulation of nucleobase-containing compound metabolic process (GO:0045934) | 1.38 | 1.10E-03 | 3.78E-02 |
| signal transduction (GO:0007165) | 1.38 | 7.87E-11 | 1.79E-08 |
| positive regulation of cellular process (GO:0048522) | 1.36 | 1.74E-12 | 5.99E-10 |
| positive regulation of biological process (GO:0048518) | 1.35 | 1.03E-12 | 3.83E-10 |
| cell surface receptor signaling pathway (GO:0007166) | 1.35 | 2.77E-04 | 1.33E-02 |
| multicellular organismal process (GO:0032501) | 1.33 | 3.34E-12 | 1.07E-09 |
| cellular response to stimulus (GO:0051716) | 1.32 | 2.84E-11 | 7.25E-09 |
| response to chemical (GO:0042221) | 1.31 | 9.92E-06 | 7.90E-04 |
| positive regulation of metabolic process (GO:0009893) | 1.30 | 2.75E-05 | 1.94E-03 |
| negative regulation of metabolic process (GO:0009892) | 1.30 | 3.77E-04 | 1.70E-02 |
| response to stress (GO:0006950) | 1.29 | 3.98E-05 | 2.71E-03 |
| transport (GO:0006810) | 1.29 | 2.65E-05 | 1.89E-03 |
| organelle organization (GO:0006996) | 1.29 | 1.83E-04 | 9.58E-03 |
| localization (GO:0051179) | 1.27 | 7.10E-06 | 5.75E-04 |
| response to stimulus (GO:0050896) | 1.26 | 1.61E-11 | 4.57E-09 |
| positive regulation of macromolecule metabolic process (GO:0010604) | 1.25 | 8.20E-04 | 3.03E-02 |
| cellular component organization (GO:0016043) | 1.25 | 9.68E-07 | 1.01E-04 |
| establishment of localization (GO:0051234) | 1.25 | 1.52E-04 | 8.42E-03 |
| regulation of biological process (GO:0050789) | 1.22 | 1.38E-16 | 2.04E-13 |
| cellular component organization or biogenesis (GO:0071840) | 1.22 | 1.76E-05 | 1.31E-03 |
| regulation of cellular process (GO:0050794) | 1.20 | 3.24E-13 | 1.33E-10 |
| biological regulation (GO:0065007) | 1.20 | 2.44E-15 | 2.01E-12 |
| cellular process (GO:0009987) | 1.13 | 4.96E-15 | 3.67E-12 |
| biological_process (GO:0008150) | 1.08 | 9.94E-12 | 2.95E-09 |
| nucleobase-containing compound metabolic process (GO:0006139) | .75 | 1.39E-03 | 4.54E-02 |
| nucleobase-containing compound biosynthetic process (GO:0034654) | .66 | 6.84E-04 | 2.62E-02 |
| nucleic acid metabolic process (GO:0090304) | .60 | 2.95E-06 | 2.63E-04 |
| RNA metabolic process (GO:0016070) | .59 | 8.15E-05 | 5.10E-03 |
| nucleic acid biosynthetic process (GO:0141187) | .55 | 2.60E-05 | 1.86E-03 |
| RNA biosynthetic process (GO:0032774) | .54 | 2.76E-05 | 1.94E-03 |
| Unclassified (UNCLASSIFIED) | .51 | 9.94E-12 | 2.89E-09 |
| detection of stimulus (GO:0051606) | .43 | 3.21E-04 | 1.48E-02 |
| mRNA metabolic process (GO:0016071) | .38 | 1.47E-04 | 8.29E-03 |
| chromosome organization (GO:0051276) | .35 | 1.49E-03 | 4.79E-02 |
| RNA splicing (GO:0008380) | .28 | 5.31E-04 | 2.15E-02 |
| RNA processing (GO:0006396) | .22 | 8.62E-10 | 1.64E-07 |
| detection of stimulus involved in sensory perception (GO:0050906) | .19 | 4.68E-07 | 5.34E-05 |
| mRNA processing (GO:0006397) | .18 | 3.85E-06 | 3.30E-04 |
| sensory perception of chemical stimulus (GO:0007606) | .15 | 1.11E-07 | 1.49E-05 |
| ribosome biogenesis (GO:0042254) | .14 | 6.75E-05 | 4.31E-03 |
| sensory perception of smell (GO:0007608) | .14 | 4.47E-07 | 5.22E-05 |
| detection of chemical stimulus (GO:0009593) | .12 | 4.97E-08 | 7.01E-06 |
| rRNA processing (GO:0006364) | .10 | 5.63E-04 | 2.23E-02 |
| ribonucleoprotein complex biogenesis (GO:0022613) | .09 | 1.27E-07 | 1.66E-05 |
| mRNA splicing, via spliceosome (GO:0000398) | .09 | 1.94E-04 | 1.00E-02 |
| RNA splicing, via transesterification reactions with bulged adenosine as nucleophile (GO:0000377) | .09 | 1.94E-04 | 9.98E-03 |
| RNA splicing, via transesterification reactions (GO:0000375) | .08 | 1.27E-04 | 7.25E-03 |
| rRNA metabolic process (GO:0016072) | .08 | 1.28E-04 | 7.27E-03 |
| protein-RNA complex organization (GO:0071826) | < 0.01 | 8.69E-05 | 5.30E-03 |
| protein-DNA complex organization (GO:0071824) | < 0.01 | 3.96E-05 | 2.71E-03 |
| protein-DNA complex assembly (GO:0065004) | < 0.01 | 8.56E-05 | 5.26E-03 |
| detection of chemical stimulus involved in sensory perception of smell (GO:0050911) | < 0.01 | 8.32E-10 | 1.60E-07 |
| detection of chemical stimulus involved in sensory perception (GO:0050907) | < 0.01 | 7.48E-11 | 1.73E-08 |
| protein-RNA complex assembly (GO:0022618) | < 0.01 | 1.28E-04 | 7.27E-03 |
