## Supplementary material for "Metabolic Maturation Unveils Left Ventricular Identity in WNT ON/OFF Human Pluripotent Stem Cell-Derived Cardiomyocytes": Table S2

**Table S2. Gene Ontology (GO) analysis of downregulated genes in mature cardiomyocytes:**

GO analysis of genes significantly downregulated in mature cardiomyocytes showed enrichment of glycolytic and related metabolic processes. GO terms are listed with corresponding fold enrichment, raw p-value, and false discovery rate (FDR)

| **GO biological process complete** | **Fold Enrichment** | **Raw P-value** | **FDR** |
| --- | --- | --- | --- |
| renal interstitial fibroblast development (GO:0072141) | 32.56 | 9.42E-04 | 3.91E-02 |
| kidney interstitial fibroblast differentiation (GO:0072071) | 32.56 | 9.42E-04 | 3.90E-02 |
| negative regulation of hair follicle maturation (GO:0048817) | 32.56 | 9.42E-04 | 3.89E-02 |
| positive regulation of hepatic stellate cell migration (GO:0061870) | 32.56 | 9.42E-04 | 3.88E-02 |
| regulation of hepatic stellate cell migration (GO:0061869) | 32.56 | 9.42E-04 | 3.86E-02 |
| response to acrylamide (GO:1903937) | 32.56 | 9.42E-04 | 3.85E-02 |
| negative regulation of cerebellar granule cell precursor proliferation (GO:0021941) | 32.56 | 9.42E-04 | 3.84E-02 |
| lateral sprouting involved in mammary gland duct morphogenesis (GO:0060599) | 32.56 | 9.42E-04 | 3.83E-02 |
| muscle cell fate determination (GO:0007521) | 32.56 | 9.42E-04 | 3.82E-02 |
| positive regulation of cell proliferation in bone marrow (GO:0071864) | 19.54 | 2.75E-04 | 1.53E-02 |
| negative regulation of hair follicle development (GO:0051799) | 19.54 | 2.75E-04 | 1.53E-02 |
| negative regulation of glomerular filtration (GO:0003105) | 19.54 | 2.75E-04 | 1.52E-02 |
| regulation of cell proliferation in bone marrow (GO:0071863) | 16.28 | 5.38E-04 | 2.58E-02 |
| convergent extension involved in axis elongation (GO:0060028) | 16.28 | 5.38E-04 | 2.57E-02 |
| 'de novo' IMP biosynthetic process (GO:0006189) | 16.28 | 5.38E-04 | 2.56E-02 |
| positive regulation of ubiquitin-protein transferase activity (GO:0051443) | 16.28 | 7.09E-08 | 2.02E-05 |
| positive regulation of ubiquitin protein ligase activity (GO:1904668) | 16.28 | 5.59E-05 | 4.06E-03 |
| regulation of hair follicle maturation (GO:0048819) | 14.47 | 9.81E-05 | 6.49E-03 |
| positive regulation of artery morphogenesis (GO:1905653) | 13.96 | 9.20E-04 | 3.87E-02 |
| regulation of artery morphogenesis (GO:1905651) | 13.96 | 9.20E-04 | 3.86E-02 |
| positive regulation of odontogenesis (GO:0042482) | 13.96 | 9.20E-04 | 3.85E-02 |
| olfactory bulb interneuron development (GO:0021891) | 13.96 | 9.20E-04 | 3.84E-02 |
| pulmonary valve morphogenesis (GO:0003184) | 13.03 | 6.88E-08 | 2.00E-05 |
| spindle elongation (GO:0051231) | 12.52 | 2.82E-05 | 2.23E-03 |
| mitotic spindle midzone assembly (GO:0051256) | 11.84 | 2.45E-04 | 1.42E-02 |
| mitotic spindle elongation (GO:0000022) | 11.84 | 2.45E-04 | 1.41E-02 |
| glucose catabolic process to pyruvate (GO:0061718) | 11.40 | 1.36E-06 | 1.94E-04 |
| canonical glycolysis (GO:0061621) | 11.40 | 1.36E-06 | 1.92E-04 |
| pulmonary valve development (GO:0003177) | 10.85 | 3.60E-07 | 6.67E-05 |
| glycolytic process through glucose-6-phosphate (GO:0061620) | 10.36 | 2.84E-06 | 3.42E-04 |
| glycolytic process through fructose-6-phosphate (GO:0061615) | 10.36 | 2.84E-06 | 3.40E-04 |
| regulation of hair follicle development (GO:0051797) | 9.77 | 2.20E-05 | 1.83E-03 |
| negative regulation of synapse organization (GO:1905809) | 9.30 | 6.90E-04 | 3.14E-02 |
| glucose catabolic process (GO:0006007) | 9.30 | 1.37E-06 | 1.91E-04 |
| fructose 6-phosphate metabolic process (GO:0006002) | 9.30 | 6.90E-04 | 3.13E-02 |
| spindle midzone assembly (GO:0051255) | 9.30 | 6.90E-04 | 3.12E-02 |
| glomerulus vasculature development (GO:0072012) | 8.88 | 4.01E-05 | 3.02E-03 |
| negative regulation of vascular permeability (GO:0043116) | 8.57 | 2.19E-04 | 1.27E-02 |
| regulation of ubiquitin-protein transferase activity (GO:0051438) | 8.14 | 1.68E-05 | 1.50E-03 |
| kidney vasculature development (GO:0061440) | 8.14 | 6.87E-05 | 4.80E-03 |
| renal system vasculature development (GO:0061437) | 8.14 | 6.87E-05 | 4.78E-03 |
| mitotic spindle assembly (GO:0090307) | 8.14 | 4.04E-09 | 1.81E-06 |
| RNA capping (GO:0036260) | 8.14 | 1.19E-03 | 4.63E-02 |
| epithelial tube branching involved in lung morphogenesis (GO:0060441) | 7.86 | 2.16E-05 | 1.82E-03 |
| establishment of apical/basal cell polarity (GO:0035089) | 7.40 | 4.59E-04 | 2.29E-02 |
| ventricular septum morphogenesis (GO:0060412) | 7.24 | 8.40E-07 | 1.27E-04 |
| mesenchymal cell proliferation (GO:0010463) | 7.24 | 1.40E-04 | 8.73E-03 |
| embryonic placenta morphogenesis (GO:0060669) | 7.24 | 1.40E-04 | 8.70E-03 |
| one-carbon metabolic process (GO:0006730) | 7.08 | 5.71E-04 | 2.70E-02 |
| labyrinthine layer morphogenesis (GO:0060713) | 7.08 | 5.71E-04 | 2.70E-02 |
| cardiac septum morphogenesis (GO:0060411) | 6.86 | 1.02E-09 | 5.78E-07 |
| regulation of megakaryocyte differentiation (GO:0045652) | 6.78 | 7.03E-04 | 3.15E-02 |
| regulation of insulin-like growth factor receptor signaling pathway (GO:0043567) | 6.78 | 7.03E-04 | 3.14E-02 |
| establishment of monopolar cell polarity (GO:0061162) | 6.78 | 7.03E-04 | 3.13E-02 |
| regulation of hair cycle (GO:0042634) | 6.74 | 2.13E-04 | 1.25E-02 |
| aortic valve morphogenesis (GO:0003180) | 6.68 | 2.01E-05 | 1.73E-03 |
| semi-lunar valve development (GO:1905314) | 6.65 | 1.94E-06 | 2.54E-04 |
| establishment or maintenance of monopolar cell polarity (GO:0061339) | 6.51 | 8.57E-04 | 3.66E-02 |
| monosaccharide catabolic process (GO:0046365) | 6.51 | 7.36E-07 | 1.17E-04 |
| aortic valve development (GO:0003176) | 6.51 | 7.56E-06 | 7.68E-04 |
| mesenchyme morphogenesis (GO:0072132) | 6.40 | 8.90E-07 | 1.29E-04 |
| hexose catabolic process (GO:0019320) | 6.38 | 2.85E-06 | 3.38E-04 |
| regulation of non-canonical Wnt signaling pathway (GO:2000050) | 6.26 | 1.03E-03 | 4.13E-02 |
| outflow tract septum morphogenesis (GO:0003148) | 6.26 | 1.03E-03 | 4.12E-02 |
| aorta development (GO:0035904) | 6.20 | 4.04E-07 | 7.21E-05 |
| glycolytic process (GO:0006096) | 6.11 | 1.32E-05 | 1.22E-03 |
| positive regulation of chromosome separation (GO:1905820) | 6.11 | 3.75E-04 | 1.95E-02 |
| regulation of neutrophil chemotaxis (GO:0090022) | 6.11 | 3.75E-04 | 1.94E-02 |
| presynapse organization (GO:0099172) | 6.11 | 3.75E-04 | 1.94E-02 |
| cardiac septum development (GO:0003279) | 6.05 | 3.07E-11 | 3.80E-08 |
| outflow tract morphogenesis (GO:0003151) | 6.03 | 2.16E-08 | 7.61E-06 |
| negative regulation of neural precursor cell proliferation (GO:2000178) | 6.03 | 1.24E-03 | 4.79E-02 |
| axis elongation (GO:0003401) | 6.03 | 1.24E-03 | 4.78E-02 |
| cell cycle G2/M phase transition (GO:0044839) | 5.92 | 5.84E-06 | 6.32E-04 |
| pyridine-containing compound catabolic process (GO:0072526) | 5.92 | 5.84E-06 | 6.27E-04 |
| pteridine-containing compound metabolic process (GO:0042558) | 5.92 | 4.47E-04 | 2.25E-02 |
| G2/M transition of mitotic cell cycle (GO:0000086) | 5.86 | 1.87E-05 | 1.64E-03 |
| mitotic spindle organization (GO:0007052) | 5.80 | 1.61E-09 | 8.23E-07 |
| ADP catabolic process (GO:0046032) | 5.75 | 2.20E-05 | 1.82E-03 |
| ribonucleoside diphosphate catabolic process (GO:0009191) | 5.71 | 8.16E-06 | 8.17E-04 |
| NADH metabolic process (GO:0006734) | 5.70 | 1.92E-04 | 1.14E-02 |
| endocardial cushion morphogenesis (GO:0003203) | 5.70 | 1.92E-04 | 1.13E-02 |
| heart valve morphogenesis (GO:0003179) | 5.69 | 3.03E-06 | 3.56E-04 |
| pyridine nucleotide catabolic process (GO:0019364) | 5.64 | 2.59E-05 | 2.08E-03 |
| ADP metabolic process (GO:0046031) | 5.61 | 9.59E-06 | 9.29E-04 |
| positive regulation of animal organ morphogenesis (GO:0110110) | 5.58 | 6.22E-04 | 2.88E-02 |
| ventricular septum development (GO:0003281) | 5.57 | 4.94E-07 | 8.31E-05 |
| nucleoside diphosphate catabolic process (GO:0009134) | 5.52 | 1.12E-05 | 1.07E-03 |
| purine ribonucleoside diphosphate catabolic process (GO:0009181) | 5.43 | 3.55E-05 | 2.72E-03 |
| purine nucleoside diphosphate catabolic process (GO:0009137) | 5.43 | 3.55E-05 | 2.71E-03 |
| cell differentiation involved in kidney development (GO:0061005) | 5.43 | 9.65E-05 | 6.44E-03 |
| mitotic sister chromatid segregation (GO:0000070) | 5.24 | 7.67E-11 | 8.11E-08 |
| non-canonical Wnt signaling pathway (GO:0035567) | 5.21 | 1.30E-04 | 8.23E-03 |
| sister chromatid segregation (GO:0000819) | 5.09 | 1.36E-10 | 1.26E-07 |
| mitotic nuclear division (GO:0140014) | 5.01 | 1.26E-11 | 1.86E-08 |
| endocardial cushion development (GO:0003197) | 5.01 | 1.73E-04 | 1.04E-02 |
| artery development (GO:0060840) | 4.92 | 1.47E-07 | 3.30E-05 |
| cardiac chamber morphogenesis (GO:0003206) | 4.90 | 4.43E-09 | 1.87E-06 |
| head morphogenesis (GO:0060323) | 4.88 | 1.29E-03 | 4.88E-02 |
| glomerulus development (GO:0032835) | 4.80 | 9.56E-05 | 6.44E-03 |
| embryonic digit morphogenesis (GO:0042733) | 4.80 | 9.56E-05 | 6.41E-03 |
| purine ribonucleotide catabolic process (GO:0009154) | 4.77 | 7.37E-06 | 7.53E-04 |
| spindle assembly (GO:0051225) | 4.65 | 7.46E-07 | 1.16E-04 |
| purine ribonucleoside diphosphate metabolic process (GO:0009179) | 4.65 | 5.24E-05 | 3.90E-03 |
| purine nucleoside diphosphate metabolic process (GO:0009135) | 4.65 | 5.24E-05 | 3.88E-03 |
| heart valve development (GO:0003170) | 4.65 | 2.23E-05 | 1.83E-03 |
| cardiac chamber development (GO:0003205) | 4.65 | 1.61E-10 | 1.40E-07 |
| labyrinthine layer development (GO:0060711) | 4.65 | 6.98E-04 | 3.14E-02 |
| mitotic metaphase chromosome alignment (GO:0007080) | 4.57 | 3.32E-04 | 1.77E-02 |
| cardiac ventricle development (GO:0003231) | 4.44 | 1.22E-07 | 3.02E-05 |
| ribonucleoside diphosphate metabolic process (GO:0009185) | 4.42 | 3.62E-05 | 2.75E-03 |
| ribonucleotide catabolic process (GO:0009261) | 4.39 | 1.74E-05 | 1.53E-03 |
| regulation of granulocyte chemotaxis (GO:0071622) | 4.38 | 1.00E-03 | 4.03E-02 |
| regulation of heart growth (GO:0060420) | 4.34 | 4.74E-04 | 2.35E-02 |
| microtubule cytoskeleton organization involved in mitosis (GO:1902850) | 4.31 | 1.94E-07 | 4.23E-05 |
| purine nucleotide catabolic process (GO:0006195) | 4.29 | 2.18E-05 | 1.83E-03 |
| regulation of chromosome separation (GO:1905818) | 4.28 | 1.07E-04 | 7.01E-03 |
| regulation of mitotic sister chromatid separation (GO:0010965) | 4.27 | 5.31E-04 | 2.55E-02 |
| cellular response to BMP stimulus (GO:0071773) | 4.25 | 2.44E-05 | 2.00E-03 |
| response to BMP (GO:0071772) | 4.25 | 2.44E-05 | 1.99E-03 |
| artery morphogenesis (GO:0048844) | 4.25 | 2.51E-04 | 1.44E-02 |
| face development (GO:0060324) | 4.22 | 1.26E-03 | 4.85E-02 |
| regulation of cardiac muscle tissue growth (GO:0055021) | 4.22 | 1.26E-03 | 4.84E-02 |
| spindle organization (GO:0007051) | 4.19 | 1.46E-08 | 5.56E-06 |
| nucleoside diphosphate metabolic process (GO:0009132) | 4.17 | 6.36E-05 | 4.47E-03 |
| mesoderm formation (GO:0001707) | 4.13 | 3.13E-04 | 1.71E-02 |
| branching morphogenesis of an epithelial tube (GO:0048754) | 4.13 | 3.74E-07 | 6.77E-05 |
| pyruvate metabolic process (GO:0006090) | 4.07 | 3.48E-04 | 1.84E-02 |
| embryonic limb morphogenesis (GO:0030326) | 4.04 | 2.21E-06 | 2.80E-04 |
| embryonic appendage morphogenesis (GO:0035113) | 4.04 | 2.21E-06 | 2.78E-04 |
| regulation of organ growth (GO:0046620) | 4.02 | 8.74E-05 | 5.94E-03 |
| mesoderm morphogenesis (GO:0048332) | 4.01 | 3.86E-04 | 1.97E-02 |
| ovulation cycle (GO:0042698) | 4.01 | 8.17E-04 | 3.54E-02 |
| collagen fibril organization (GO:0030199) | 4.01 | 8.17E-04 | 3.53E-02 |
| regulation of mitotic metaphase/anaphase transition (GO:0030071) | 3.89 | 1.18E-04 | 7.66E-03 |
| epithelial to mesenchymal transition (GO:0001837) | 3.88 | 2.48E-04 | 1.43E-02 |
| nuclear chromosome segregation (GO:0098813) | 3.85 | 1.15E-09 | 6.31E-07 |
| mesenchyme development (GO:0060485) | 3.81 | 7.40E-10 | 4.57E-07 |
| positive regulation of axonogenesis (GO:0050772) | 3.81 | 5.76E-04 | 2.70E-02 |
| morphogenesis of a branching epithelium (GO:0061138) | 3.79 | 3.54E-07 | 6.64E-05 |
| regulation of metaphase/anaphase transition of cell cycle (GO:1902099) | 3.77 | 1.58E-04 | 9.69E-03 |
| limb morphogenesis (GO:0035108) | 3.76 | 1.52E-06 | 2.10E-04 |
| appendage morphogenesis (GO:0035107) | 3.76 | 1.52E-06 | 2.08E-04 |
| mesenchymal cell differentiation (GO:0048762) | 3.74 | 4.27E-07 | 7.44E-05 |
| canonical Wnt signaling pathway (GO:0060070) | 3.72 | 9.15E-05 | 6.19E-03 |
| cell cycle phase transition (GO:0044770) | 3.71 | 1.83E-06 | 2.44E-04 |
| BMP signaling pathway (GO:0030509) | 3.71 | 6.96E-04 | 3.14E-02 |
| mitotic cytokinesis (GO:0000281) | 3.66 | 3.99E-04 | 2.02E-02 |
| purine-containing compound catabolic process (GO:0072523) | 3.62 | 1.20E-04 | 7.75E-03 |
| mitotic cell cycle phase transition (GO:0044772) | 3.59 | 1.02E-05 | 9.83E-04 |
| embryonic placenta development (GO:0001892) | 3.58 | 4.77E-04 | 2.36E-02 |
| morphogenesis of a branching structure (GO:0001763) | 3.58 | 8.79E-07 | 1.30E-04 |
| roof of mouth development (GO:0060021) | 3.54 | 5.21E-04 | 2.55E-02 |
| regulation of animal organ morphogenesis (GO:2000027) | 3.49 | 1.09E-03 | 4.29E-02 |
| establishment of cell polarity (GO:0030010) | 3.46 | 2.97E-05 | 2.33E-03 |
| metaphase chromosome alignment (GO:0051310) | 3.45 | 1.18E-03 | 4.60E-02 |
| regulation of chromosome segregation (GO:0051983) | 3.43 | 6.10E-05 | 4.35E-03 |
| nucleotide catabolic process (GO:0009166) | 3.43 | 2.02E-04 | 1.19E-02 |
| ureteric bud development (GO:0001657) | 3.41 | 1.29E-03 | 4.88E-02 |
| carbohydrate catabolic process (GO:0016052) | 3.39 | 1.25E-04 | 8.04E-03 |
| cardiocyte differentiation (GO:0035051) | 3.39 | 1.25E-04 | 8.01E-03 |
| negative regulation of Wnt signaling pathway (GO:0030178) | 3.37 | 7.23E-06 | 7.44E-04 |
| regulation of sister chromatid segregation (GO:0033045) | 3.35 | 4.51E-04 | 2.26E-02 |
| cytoskeleton-dependent cytokinesis (GO:0061640) | 3.34 | 2.58E-04 | 1.45E-02 |
| cardiac muscle tissue development (GO:0048738) | 3.32 | 2.81E-06 | 3.41E-04 |
| establishment of chromosome localization (GO:0051303) | 3.32 | 8.59E-04 | 3.66E-02 |
| positive regulation of protein ubiquitination (GO:0031398) | 3.29 | 5.29E-04 | 2.57E-02 |
| positive regulation of endothelial cell migration (GO:0010595) | 3.29 | 5.29E-04 | 2.56E-02 |
| cytokinesis (GO:0000910) | 3.28 | 3.02E-04 | 1.66E-02 |
| establishment or maintenance of cell polarity (GO:0007163) | 3.27 | 2.04E-06 | 2.66E-04 |
| negative regulation of canonical Wnt signaling pathway (GO:0090090) | 3.26 | 1.07E-04 | 7.02E-03 |
| odontogenesis (GO:0042476) | 3.26 | 3.26E-04 | 1.76E-02 |
| nucleosome organization (GO:0034728) | 3.26 | 5.72E-04 | 2.69E-02 |
| heart morphogenesis (GO:0003007) | 3.22 | 1.75E-07 | 3.88E-05 |
| limb development (GO:0060173) | 3.19 | 9.01E-06 | 8.90E-04 |
| appendage development (GO:0048736) | 3.19 | 9.01E-06 | 8.84E-04 |
| postsynapse organization (GO:0099173) | 3.17 | 7.18E-04 | 3.17E-02 |
| cell-substrate adhesion (GO:0031589) | 3.17 | 1.69E-05 | 1.50E-03 |
| developmental growth involved in morphogenesis (GO:0060560) | 3.17 | 1.44E-04 | 8.90E-03 |
| nucleoside phosphate catabolic process (GO:1901292) | 3.16 | 2.52E-04 | 1.44E-02 |
| epithelial tube morphogenesis (GO:0060562) | 3.08 | 5.33E-08 | 1.64E-05 |
| regulation of synapse assembly (GO:0051963) | 3.05 | 7.84E-05 | 5.38E-03 |
| kidney epithelium development (GO:0072073) | 3.05 | 3.61E-04 | 1.89E-02 |
| cell-cell junction assembly (GO:0007043) | 3.01 | 6.76E-04 | 3.09E-02 |
| nuclear division (GO:0000280) | 2.99 | 1.06E-07 | 2.71E-05 |
| regulation of mitotic nuclear division (GO:0007088) | 2.98 | 1.18E-03 | 4.61E-02 |
| actomyosin structure organization (GO:0031032) | 2.96 | 1.27E-03 | 4.85E-02 |
| epithelial tube formation (GO:0072175) | 2.94 | 8.28E-04 | 3.55E-02 |
| mitotic cell cycle process (GO:1903047) | 2.93 | 7.65E-11 | 8.72E-08 |
| ear morphogenesis (GO:0042471) | 2.89 | 9.44E-04 | 3.82E-02 |
| chromosome segregation (GO:0007059) | 2.88 | 2.33E-07 | 4.66E-05 |
| renal system development (GO:0072001) | 2.88 | 6.09E-07 | 9.92E-05 |
| kidney development (GO:0001822) | 2.87 | 9.86E-07 | 1.42E-04 |
| regulation of epithelial cell differentiation (GO:0030856) | 2.87 | 4.04E-04 | 2.04E-02 |
| regulation of nuclear division (GO:0051783) | 2.86 | 6.58E-04 | 3.03E-02 |
| positive regulation of Wnt signaling pathway (GO:0030177) | 2.86 | 6.58E-04 | 3.02E-02 |
| organelle fission (GO:0048285) | 2.84 | 1.97E-07 | 4.23E-05 |
| cell division (GO:0051301) | 2.84 | 2.13E-10 | 1.66E-07 |
| morphogenesis of embryonic epithelium (GO:0016331) | 2.84 | 7.02E-04 | 3.15E-02 |
| positive regulation of transferase activity (GO:0051347) | 2.81 | 3.20E-04 | 1.74E-02 |
| regulation of Wnt signaling pathway (GO:0030111) | 2.79 | 7.45E-07 | 1.17E-04 |
| positive regulation of neuron projection development (GO:0010976) | 2.77 | 8.99E-04 | 3.79E-02 |
| mitotic cell cycle (GO:0000278) | 2.75 | 1.02E-10 | 1.01E-07 |
| tissue morphogenesis (GO:0048729) | 2.75 | 1.67E-10 | 1.37E-07 |
| positive regulation of cell cycle process (GO:0090068) | 2.74 | 2.01E-05 | 1.72E-03 |
| stem cell differentiation (GO:0048863) | 2.69 | 3.38E-04 | 1.79E-02 |
| morphogenesis of an epithelium (GO:0002009) | 2.67 | 4.54E-08 | 1.46E-05 |
| positive regulation of cell cycle (GO:0045787) | 2.66 | 4.15E-06 | 4.66E-04 |
| placenta development (GO:0001890) | 2.66 | 1.28E-03 | 4.88E-02 |
| regulation of endothelial cell migration (GO:0010594) | 2.65 | 8.85E-04 | 3.75E-02 |
| cell-cell junction organization (GO:0045216) | 2.64 | 6.12E-04 | 2.84E-02 |
| chromosome organization (GO:0051276) | 2.63 | 3.24E-07 | 6.16E-05 |
| regulation of synapse organization (GO:0050807) | 2.63 | 1.20E-05 | 1.13E-03 |
| ear development (GO:0043583) | 2.62 | 1.34E-04 | 8.42E-03 |
| Wnt signaling pathway (GO:0016055) | 2.62 | 2.78E-05 | 2.21E-03 |
| positive regulation of catalytic activity (GO:0043085) | 2.59 | 7.20E-06 | 7.46E-04 |
| regulation of synapse structure or activity (GO:0050803) | 2.58 | 1.66E-05 | 1.49E-03 |
| organophosphate catabolic process (GO:0046434) | 2.57 | 8.06E-04 | 3.50E-02 |
| monosaccharide metabolic process (GO:0005996) | 2.56 | 8.51E-04 | 3.64E-02 |
| membraneless organelle assembly (GO:0140694) | 2.55 | 6.52E-06 | 6.86E-04 |
| regulation of neuron projection development (GO:0010975) | 2.52 | 8.49E-07 | 1.27E-04 |
| inner ear development (GO:0048839) | 2.50 | 7.26E-04 | 3.20E-02 |
| heart development (GO:0007507) | 2.50 | 3.96E-08 | 1.30E-05 |
| vasculature development (GO:0001944) | 2.50 | 5.95E-08 | 1.80E-05 |
| regulation of canonical Wnt signaling pathway (GO:0060828) | 2.50 | 1.75E-04 | 1.06E-02 |
| blood vessel development (GO:0001568) | 2.49 | 1.35E-07 | 3.22E-05 |
| positive regulation of neurogenesis (GO:0050769) | 2.43 | 7.14E-04 | 3.17E-02 |
| gliogenesis (GO:0042063) | 2.42 | 1.81E-04 | 1.08E-02 |
| tube morphogenesis (GO:0035239) | 2.42 | 4.41E-09 | 1.92E-06 |
| regulation of cellular response to growth factor stimulus (GO:0090287) | 2.40 | 1.10E-04 | 7.16E-03 |
| nucleoside triphosphate metabolic process (GO:0009141) | 2.40 | 1.14E-03 | 4.46E-02 |
| actin cytoskeleton organization (GO:0030036) | 2.40 | 3.74E-07 | 6.85E-05 |
| cell projection morphogenesis (GO:0048858) | 2.40 | 7.57E-07 | 1.17E-04 |
| in utero embryonic development (GO:0001701) | 2.40 | 8.95E-06 | 8.90E-04 |
| embryonic organ development (GO:0048568) | 2.37 | 3.90E-06 | 4.45E-04 |
| cell junction assembly (GO:0034329) | 2.36 | 2.57E-04 | 1.45E-02 |
| purine ribonucleotide metabolic process (GO:0009150) | 2.35 | 3.85E-04 | 1.97E-02 |
| cell morphogenesis (GO:0000902) | 2.34 | 1.55E-08 | 5.75E-06 |
| neuron projection morphogenesis (GO:0048812) | 2.33 | 2.81E-06 | 3.44E-04 |
| actin filament-based process (GO:0030029) | 2.32 | 2.85E-07 | 5.56E-05 |
| plasma membrane bounded cell projection morphogenesis (GO:0120039) | 2.30 | 5.18E-06 | 5.69E-04 |
| supramolecular fiber organization (GO:0097435) | 2.30 | 4.78E-07 | 8.24E-05 |
| ribonucleotide metabolic process (GO:0009259) | 2.30 | 3.32E-04 | 1.77E-02 |
| blood vessel morphogenesis (GO:0048514) | 2.28 | 1.97E-05 | 1.71E-03 |
| negative regulation of locomotion (GO:0040013) | 2.27 | 2.72E-04 | 1.52E-02 |
| cell morphogenesis involved in neuron differentiation (GO:0048667) | 2.26 | 2.47E-05 | 2.00E-03 |
| extracellular matrix organization (GO:0030198) | 2.25 | 7.68E-04 | 3.38E-02 |
| embryonic organ morphogenesis (GO:0048562) | 2.25 | 5.67E-04 | 2.69E-02 |
| muscle cell differentiation (GO:0042692) | 2.25 | 7.93E-04 | 3.47E-02 |
| extracellular structure organization (GO:0043062) | 2.25 | 7.93E-04 | 3.46E-02 |
| ribose phosphate metabolic process (GO:0019693) | 2.25 | 4.21E-04 | 2.12E-02 |
| tube development (GO:0035295) | 2.24 | 2.01E-09 | 9.30E-07 |
| external encapsulating structure organization (GO:0045229) | 2.24 | 8.20E-04 | 3.53E-02 |
| regulation of neurogenesis (GO:0050767) | 2.23 | 1.82E-04 | 1.08E-02 |
| circulatory system development (GO:0072359) | 2.23 | 1.64E-09 | 7.85E-07 |
| angiogenesis (GO:0001525) | 2.23 | 2.52E-04 | 1.43E-02 |
| positive regulation of cell projection organization (GO:0031346) | 2.23 | 2.52E-04 | 1.43E-02 |
| cell junction organization (GO:0034330) | 2.22 | 3.33E-06 | 3.86E-04 |
| regulation of anatomical structure morphogenesis (GO:0022603) | 2.21 | 2.74E-08 | 9.23E-06 |
| positive regulation of nervous system development (GO:0051962) | 2.21 | 1.27E-03 | 4.83E-02 |
| embryonic morphogenesis (GO:0048598) | 2.20 | 3.29E-06 | 3.83E-04 |
| muscle tissue development (GO:0060537) | 2.20 | 4.61E-04 | 2.29E-02 |
| cell-cell adhesion (GO:0098609) | 2.19 | 1.27E-05 | 1.18E-03 |
| cellular response to growth factor stimulus (GO:0071363) | 2.18 | 3.93E-05 | 2.97E-03 |
| chordate embryonic development (GO:0043009) | 2.17 | 8.86E-07 | 1.30E-04 |
| regulation of developmental growth (GO:0048638) | 2.16 | 8.66E-04 | 3.68E-02 |
| regulation of epithelial cell proliferation (GO:0050678) | 2.15 | 5.25E-04 | 2.56E-02 |
| animal organ morphogenesis (GO:0009887) | 2.15 | 6.61E-09 | 2.72E-06 |
| modulation of chemical synaptic transmission (GO:0050804) | 2.13 | 6.02E-05 | 4.31E-03 |
| striated muscle tissue development (GO:0014706) | 2.13 | 1.08E-03 | 4.27E-02 |
| regulation of trans-synaptic signaling (GO:0099177) | 2.12 | 6.14E-05 | 4.35E-03 |
| growth (GO:0040007) | 2.11 | 1.64E-04 | 1.00E-02 |
| developmental growth (GO:0048589) | 2.11 | 1.64E-04 | 9.97E-03 |
| embryo development ending in birth or egg hatching (GO:0009792) | 2.11 | 2.07E-06 | 2.67E-04 |
| synapse organization (GO:0050808) | 2.11 | 6.37E-04 | 2.94E-02 |
| regulation of nervous system development (GO:0051960) | 2.10 | 2.34E-04 | 1.36E-02 |
| cell cycle process (GO:0022402) | 2.09 | 1.06E-07 | 2.67E-05 |
| sensory organ development (GO:0007423) | 2.08 | 1.52E-05 | 1.39E-03 |
| cell migration (GO:0016477) | 2.08 | 9.81E-08 | 2.64E-05 |
| axonogenesis (GO:0007409) | 2.07 | 7.76E-04 | 3.40E-02 |
| positive regulation of cell development (GO:0010720) | 2.07 | 3.61E-04 | 1.88E-02 |
| phosphorylation (GO:0016310) | 2.07 | 5.96E-04 | 2.78E-02 |
| neuron projection development (GO:0031175) | 2.07 | 4.53E-06 | 5.00E-04 |
| cytoskeleton organization (GO:0007010) | 2.07 | 4.61E-10 | 3.10E-07 |
| regulation of locomotion (GO:0040012) | 2.06 | 2.19E-08 | 7.53E-06 |
| cell adhesion (GO:0007155) | 2.06 | 8.94E-08 | 2.45E-05 |
| response to growth factor (GO:0070848) | 2.05 | 1.29E-04 | 8.23E-03 |
| regulation of plasma membrane bounded cell projection organization (GO:0120035) | 2.04 | 1.97E-05 | 1.72E-03 |
| regulation of cell migration (GO:0030334) | 2.04 | 2.22E-07 | 4.64E-05 |
| regulation of cell projection organization (GO:0031344) | 2.04 | 1.59E-05 | 1.45E-03 |
| positive regulation of molecular function (GO:0044093) | 2.02 | 7.00E-05 | 4.85E-03 |
| anatomical structure morphogenesis (GO:0009653) | 2.01 | 3.38E-16 | 2.51E-12 |
| cell population proliferation (GO:0008283) | 2.01 | 1.09E-05 | 1.04E-03 |
| cell cycle (GO:0007049) | 2.00 | 2.35E-07 | 4.64E-05 |
| positive regulation of cell migration (GO:0030335) | 2.00 | 1.37E-04 | 8.57E-03 |
| regulation of cell cycle process (GO:0010564) | 1.99 | 1.61E-05 | 1.45E-03 |
| regulation of cell motility (GO:2000145) | 1.99 | 3.02E-07 | 5.82E-05 |
| anatomical structure formation involved in morphogenesis (GO:0048646) | 1.97 | 6.01E-07 | 9.90E-05 |
| positive regulation of locomotion (GO:0040017) | 1.97 | 1.41E-04 | 8.77E-03 |
| epithelium development (GO:0060429) | 1.96 | 1.06E-07 | 2.75E-05 |
| regulation of catalytic activity (GO:0050790) | 1.95 | 1.53E-04 | 9.39E-03 |
| embryo development (GO:0009790) | 1.93 | 4.26E-07 | 7.51E-05 |
| neuron development (GO:0048666) | 1.92 | 6.20E-06 | 6.60E-04 |
| positive regulation of cell motility (GO:2000147) | 1.91 | 3.26E-04 | 1.75E-02 |
| positive regulation of cell differentiation (GO:0045597) | 1.91 | 1.15E-05 | 1.09E-03 |
| negative regulation of apoptotic process (GO:0043066) | 1.91 | 9.07E-06 | 8.84E-04 |
| skeletal system development (GO:0001501) | 1.90 | 1.00E-03 | 4.03E-02 |
| negative regulation of cell population proliferation (GO:0008285) | 1.90 | 8.39E-05 | 5.73E-03 |
| microtubule cytoskeleton organization (GO:0000226) | 1.90 | 3.53E-04 | 1.86E-02 |
| regulation of cell development (GO:0060284) | 1.89 | 2.90E-05 | 2.28E-03 |
| positive regulation of developmental process (GO:0051094) | 1.89 | 6.35E-08 | 1.88E-05 |
| cell death (GO:0008219) | 1.87 | 1.56E-06 | 2.12E-04 |
| apoptotic process (GO:0006915) | 1.86 | 3.91E-06 | 4.42E-04 |
| positive regulation of cell population proliferation (GO:0008284) | 1.86 | 1.50E-05 | 1.38E-03 |
| programmed cell death (GO:0012501) | 1.85 | 2.40E-06 | 2.99E-04 |
| muscle structure development (GO:0061061) | 1.84 | 1.28E-03 | 4.87E-02 |
| negative regulation of programmed cell death (GO:0043069) | 1.83 | 3.28E-05 | 2.56E-03 |
| multicellular organismal-level homeostasis (GO:0048871) | 1.83 | 4.78E-04 | 2.35E-02 |
| regulation of multicellular organismal development (GO:2000026) | 1.83 | 2.07E-07 | 4.38E-05 |
| head development (GO:0060322) | 1.83 | 9.68E-05 | 6.43E-03 |
| neurogenesis (GO:0022008) | 1.82 | 2.26E-07 | 4.64E-05 |
| neuron differentiation (GO:0030182) | 1.80 | 5.41E-06 | 5.89E-04 |
| generation of neurons (GO:0048699) | 1.79 | 4.40E-06 | 4.91E-04 |
| positive regulation of multicellular organismal process (GO:0051240) | 1.76 | 1.05E-07 | 2.78E-05 |
| regulation of cell population proliferation (GO:0042127) | 1.75 | 2.27E-07 | 4.61E-05 |
| tissue development (GO:0009888) | 1.74 | 8.90E-08 | 2.49E-05 |
| regulation of cell cycle (GO:0051726) | 1.73 | 5.36E-05 | 3.91E-03 |
| response to lipid (GO:0033993) | 1.71 | 5.05E-04 | 2.48E-02 |
| regulation of cell differentiation (GO:0045595) | 1.71 | 1.81E-06 | 2.44E-04 |
| negative regulation of developmental process (GO:0051093) | 1.71 | 2.87E-04 | 1.58E-02 |
| positive regulation of cellular component organization (GO:0051130) | 1.70 | 7.57E-05 | 5.21E-03 |
| regulation of developmental process (GO:0050793) | 1.70 | 1.51E-09 | 7.99E-07 |
| cell motility (GO:0048870) | 1.69 | 5.33E-05 | 3.91E-03 |
| regulation of apoptotic process (GO:0042981) | 1.69 | 6.37E-06 | 6.74E-04 |
| negative regulation of multicellular organismal process (GO:0051241) | 1.66 | 1.78E-04 | 1.07E-02 |
| regulation of organelle organization (GO:0033043) | 1.66 | 1.18E-04 | 7.65E-03 |
| regulation of cellular component biogenesis (GO:0044087) | 1.65 | 3.81E-04 | 1.96E-02 |
| regulation of cellular component organization (GO:0051128) | 1.65 | 9.13E-09 | 3.66E-06 |
| negative regulation of signal transduction (GO:0009968) | 1.65 | 3.50E-05 | 2.70E-03 |
| regulation of programmed cell death (GO:0043067) | 1.64 | 2.51E-05 | 2.02E-03 |
| nervous system development (GO:0007399) | 1.63 | 1.45E-07 | 3.36E-05 |
| central nervous system development (GO:0007417) | 1.63 | 5.84E-04 | 2.73E-02 |
| animal organ development (GO:0048513) | 1.63 | 1.64E-09 | 8.08E-07 |
| negative regulation of transcription by RNA polymerase II (GO:0000122) | 1.62 | 1.05E-03 | 4.16E-02 |
| regulation of molecular function (GO:0065009) | 1.61 | 9.80E-04 | 3.95E-02 |
| system development (GO:0048731) | 1.61 | 1.61E-11 | 2.17E-08 |
| negative regulation of cell communication (GO:0010648) | 1.61 | 5.92E-05 | 4.28E-03 |
| negative regulation of signaling (GO:0023057) | 1.61 | 5.97E-05 | 4.29E-03 |
| cell development (GO:0048468) | 1.59 | 4.84E-07 | 8.24E-05 |
| multicellular organism development (GO:0007275) | 1.58 | 4.04E-12 | 7.49E-09 |
| regulation of multicellular organismal process (GO:0051239) | 1.57 | 2.04E-08 | 7.38E-06 |
| regulation of signaling (GO:0023051) | 1.56 | 5.72E-10 | 3.68E-07 |
| regulation of cell communication (GO:0010646) | 1.56 | 7.93E-10 | 4.70E-07 |
| phosphate-containing compound metabolic process (GO:0006796) | 1.55 | 3.17E-04 | 1.73E-02 |
| phosphorus metabolic process (GO:0006793) | 1.55 | 3.24E-04 | 1.75E-02 |
| cell projection organization (GO:0030030) | 1.54 | 9.21E-04 | 3.83E-02 |
| regulation of signal transduction (GO:0009966) | 1.52 | 1.25E-07 | 3.04E-05 |
| anatomical structure development (GO:0048856) | 1.50 | 3.76E-13 | 9.28E-10 |
| regulation of protein metabolic process (GO:0051246) | 1.49 | 1.08E-03 | 4.27E-02 |
| negative regulation of response to stimulus (GO:0048585) | 1.49 | 3.49E-04 | 1.84E-02 |
| cellular component organization (GO:0016043) | 1.48 | 2.49E-13 | 7.36E-10 |
| cellular component assembly (GO:0022607) | 1.46 | 3.29E-05 | 2.55E-03 |
| cellular component organization or biogenesis (GO:0071840) | 1.46 | 4.80E-13 | 1.02E-09 |
| cell differentiation (GO:0030154) | 1.46 | 1.44E-07 | 3.39E-05 |
| cellular developmental process (GO:0048869) | 1.46 | 1.46E-07 | 3.34E-05 |
| negative regulation of cellular process (GO:0048523) | 1.45 | 3.16E-10 | 2.34E-07 |
| developmental process (GO:0032502) | 1.44 | 9.32E-12 | 1.53E-08 |
| negative regulation of biological process (GO:0048519) | 1.43 | 4.28E-10 | 3.02E-07 |
| cellular component biogenesis (GO:0044085) | 1.43 | 4.09E-05 | 3.06E-03 |
| regulation of biological quality (GO:0065008) | 1.42 | 2.15E-05 | 1.82E-03 |
| organelle organization (GO:0006996) | 1.41 | 2.07E-05 | 1.77E-03 |
| regulation of intracellular signal transduction (GO:1902531) | 1.41 | 1.03E-03 | 4.11E-02 |
| positive regulation of cellular process (GO:0048522) | 1.37 | 1.34E-08 | 5.22E-06 |
| positive regulation of macromolecule metabolic process (GO:0010604) | 1.34 | 3.89E-04 | 1.98E-02 |
| positive regulation of biological process (GO:0048518) | 1.34 | 4.85E-08 | 1.53E-05 |
| regulation of response to stimulus (GO:0048583) | 1.32 | 6.27E-05 | 4.42E-03 |
| positive regulation of metabolic process (GO:0009893) | 1.29 | 1.17E-03 | 4.59E-02 |
| multicellular organismal process (GO:0032501) | 1.28 | 3.56E-06 | 4.09E-04 |
| cellular response to stimulus (GO:0051716) | 1.19 | 1.09E-03 | 4.30E-02 |
| regulation of cellular process (GO:0050794) | 1.17 | 8.14E-07 | 1.24E-04 |
| cellular process (GO:0009987) | 1.17 | 1.24E-16 | 1.84E-12 |
| response to stimulus (GO:0050896) | 1.17 | 5.29E-04 | 2.55E-02 |
| regulation of biological process (GO:0050789) | 1.17 | 7.18E-07 | 1.16E-04 |
| biological regulation (GO:0065007) | 1.15 | 2.58E-06 | 3.19E-04 |
| biological_process (GO:0008150) | 1.11 | 3.22E-15 | 1.59E-11 |
| Unclassified (UNCLASSIFIED) | .32 | 3.22E-15 | 1.19E-11 |
| detection of stimulus involved in sensory perception (GO:0050906) | .17 | 5.31E-05 | 3.91E-03 |
| sensory perception of chemical stimulus (GO:0007606) | .12 | 8.12E-06 | 8.19E-04 |
| detection of chemical stimulus involved in sensory perception (GO:0050907) | .07 | 6.76E-06 | 7.05E-04 |
| detection of chemical stimulus (GO:0009593) | .06 | 2.13E-06 | 2.73E-04 |
| detection of chemical stimulus involved in sensory perception of smell (GO:0050911) | < 0.01 | 1.84E-06 | 2.43E-04 |
| sensory perception of smell (GO:0007608) | < 0.01 | 5.46E-07 | 9.08E-05 |
