## Supplementary material for "Metabolic Maturation Unveils Left Ventricular Identity in WNT ON/OFF Human Pluripotent Stem Cell-Derived Cardiomyocytes": Table S3

**Table S3. Primer sequences used for quantitative PCR (qPCR) analysis:**

List of forward and reverse primers used for qPCR in this study. Primer sequences are shown in the 5′→3′ orientation, along with their corresponding target genes.

| **Gene** | **Primer Direction** | **Sequence** |
| --- | --- | --- |
| GAPDH | Fw | ACAGCCTCAAGATCATCAG |
| GAPDH | Rv | GAGTCCTTCCACGATACC |
| RPL7 | Fw | AATGGCGAGGATGGCAAG |
| RPL7 | Rv | TGACGAAGGCGAAGAAGC |
| TNNT (cTnT) | Fw | TGGTGCCTCCCAAGATCCCCG |
| TNNT (cTnT) | Rv | AAGTGAGCCTCGATCAGCGCC |
| SOX2 | Fw | AGCATGGAGAAAACCCGGTACGC |
| SOX2 | Rv | CGTGAGTGTGGATGGGATTGGTGT |
| TBXT | Fw | TCCCAGGTGGCTTACAGATGA |
| TBXT | Rv | GGTGTGCCAAAGTTGCCAAT |
| NKX2.5 | Fw | CCCACGCCCTTCTCAGTCAA |
| NKX2.5 | Rv | GTAGGCCTCTGGCTTGAAGG |
| LEF1 | Fw | TCTGCATCAGGTGGAAAACGA |
| LEF1 | Rv | GCGTCTCTAGCAGTGACCTC |
| CX43 | Fw | ATTAGGGGGAAGGCGTGAGG |
| CX43 | Rv | CCTAAGGCGCTCCAGTCAC |
| MESP1 | Fw | GCCACTTCACACCTCGGGCTC |
| MESP1 | Rv | CCAGGCCGCAGAGAGCATCCA |
| PDGFRa | Fw | CGTTCCTGGTCTTAGGCTGTC |
| PDGFRa | Rv | ACTTCACTCTCCCCAAAGCATC |
| TNNI1 | Fw | TACCTGGCAGAGCGCATCC |
| TNNI1 | Rv | CTTCAGGTCCTTAATCTCCCTGG |
| MYL2 | Fw | ACGTTCGGGAAATGCTGACC |
| MYL2 | Rv | TTCTCCGTGGGTGATGATGTG |
| MYL7 | Fw | AGGCCCAACGTGGTTCTTC |
| MYL7 | Rv | GCCATCACGATTCTGGTCGA |
| PPARGC1A | Fw | TGGATGAAGACGGATTGCCC |
| PPARGC1A | Rv | TAGCTGAGTGTTGGCTGGTG |
| RYR2 | Fw | TGCCTCTAAGCAGCGATCAG |
| RYR2 | Rv | TGTAAGCTGCCGTTGCCATA |
| TNNI3 | Fw | CGTGTGGACAAGGTGGATGA |
| TNNI3 | Rv | AGAGATCCTCACTCTCCGCA |
| COX6A2 | Fw | AACTCCTATCTCCACTCGGG |
| COX6A2 | Rv | GGTGTTCGTAGCCCGTGG |
| MYH6 | Fw | TCCGTGAAGGGATAACCAGG |
| MYH6 | Rv | CTCTTCCTTGTCATCGGGCA |
| MYH7 | Fw | TTGGCCCCTTTCCTCATCTG |
| MYH7 | Rv | TGAGGTCAAAAGGCCTGGTC |
| EOMES | Fw | CGGAGCCCTTTGTCAACACT |
| EOMES | Rv | TTCGCTCTGTTGGGGTGAAA |
| OCT4 | Fw | CTGGGTTGATCCTCGGACCT |
| OCT4 | Rv | CACAGAACTCATACGGCGGG |
| HAND1 | Fw | GGAGACGCACTGAGAGCATTA |
| HAND1 | Rv | GGAAAACCTTCGTGCTGCAG |
| KDR | Fw | CCCGGAGTGACCAAGGATTG |
| KDR | Rv | TGCCCCGCTTTAATTGTGTG |
| TBP | Fw | AAGACCATTGCACTTCGTGCC |
| TBP | Rv | TGGACTGTTCTTCACTCTTGGC |
| mtDNA rRNA | Fw | GCCTTCCCCCGTAAATGATA |
| mtDNA rRNA | Rv | TTATGCGATTACCGGGCTCT |
| b2-microglobulin | Fw | TGCTGTCTCCATGTTTGATGTATCT |
| b2-microglobulin | Rv | TCTCTGCTCCCCACCTCTAAGT |
| FABP3 | Fw | TGACAGGAAGGTCAAGTCCATTG |
| FABP3 | Rv | TGCAGTCAGGTCATGCCTCT |
| CPT1B | Fw | AGGAGCCCTCTCATGGTGAA |
| CPT1B | Rv | CAGTGCCATCACAGGCTTGA |
| CD36 | Fw | GCAAAACGGCTGCAGGTCAA |
| CD36 | Rv | TTCTCATCACCAATGGTCCCAG |
