## Supplementary Data for "Metabolic Maturation Unveils Left Ventricular Identity in WNT ON/OFF Human Pluripotent Stem Cell-Derived Cardiomyocytes"

#### SUPPLEMENTAL FIGURES

##### Figure S1

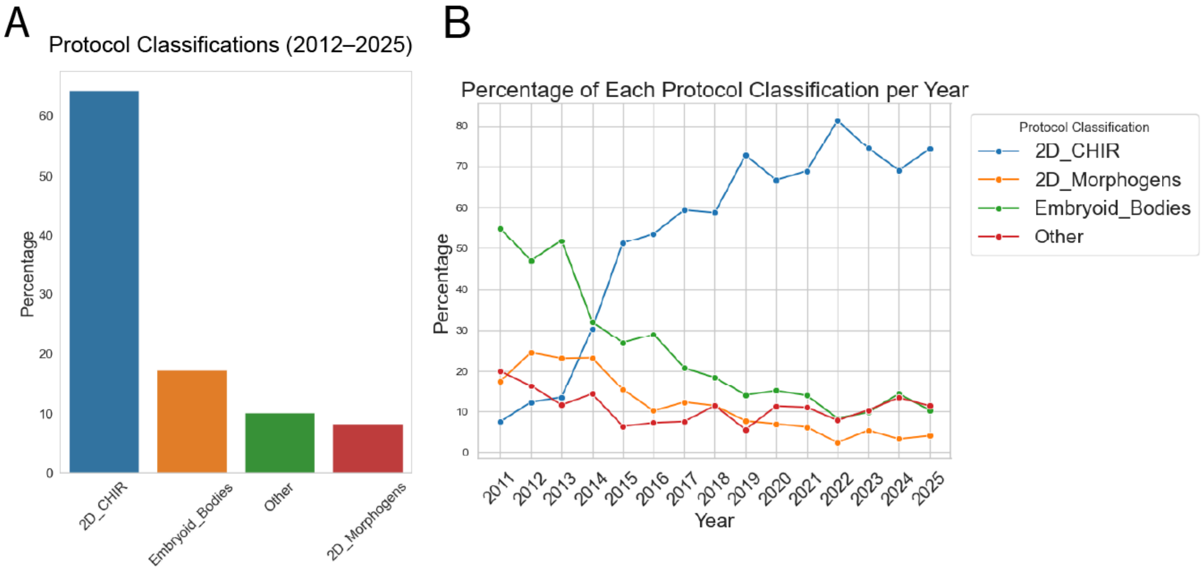


**Figure S1: Quantitative bibliographic analysis of over 1800 freely available PubMed Central papers published between 2011 and 2025**. A) Frequency of small molecule WNT ON/OFF protocols since 2012. B) Cardiac differentiation protocol category tendency shifts over time for over 1800 published papers.

**Figure S2**

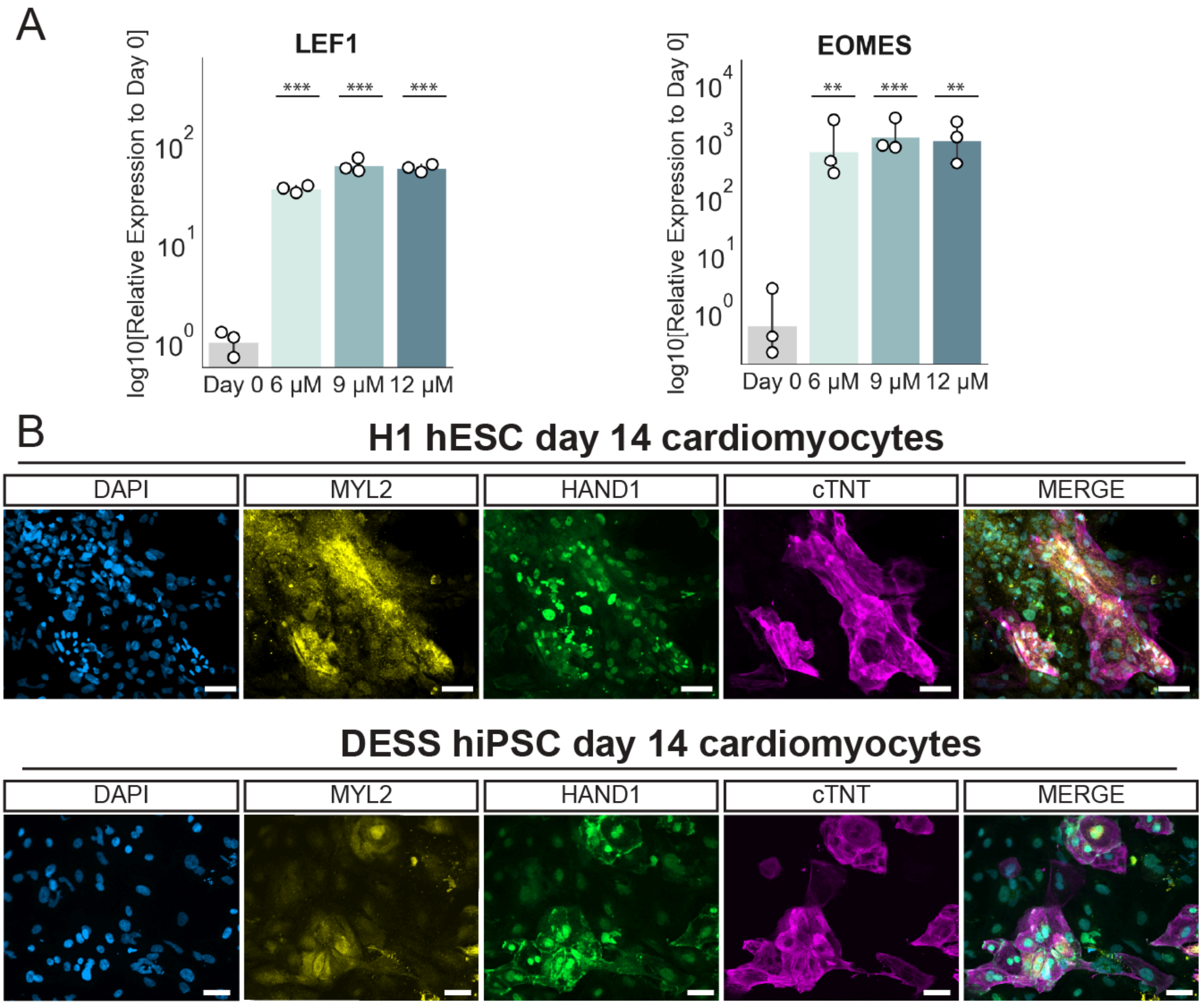


**Figure S2: WNT ON/OFF differentiation produces cardiomyocytes via a first heart field–like mesodermal stage in hESC and hiPSC.** A) RT-qPCR analysis of LEF1 and EOMES at day 1, n=3. B) Immunofluorescence staining of HAND1, MYL2, cTNT, and DAPI in cardiomyocytes derived from H1 hESC and DESS hiPSC cell lines using 12 μM CHIR, n=3. Scale bar = 50 μm. Data represent mean and individual values from independent experiments. *$p < 0.05$, **$p < 0.01$, ***$p < 0.001$; n.s., not significant.

#### Figure S3

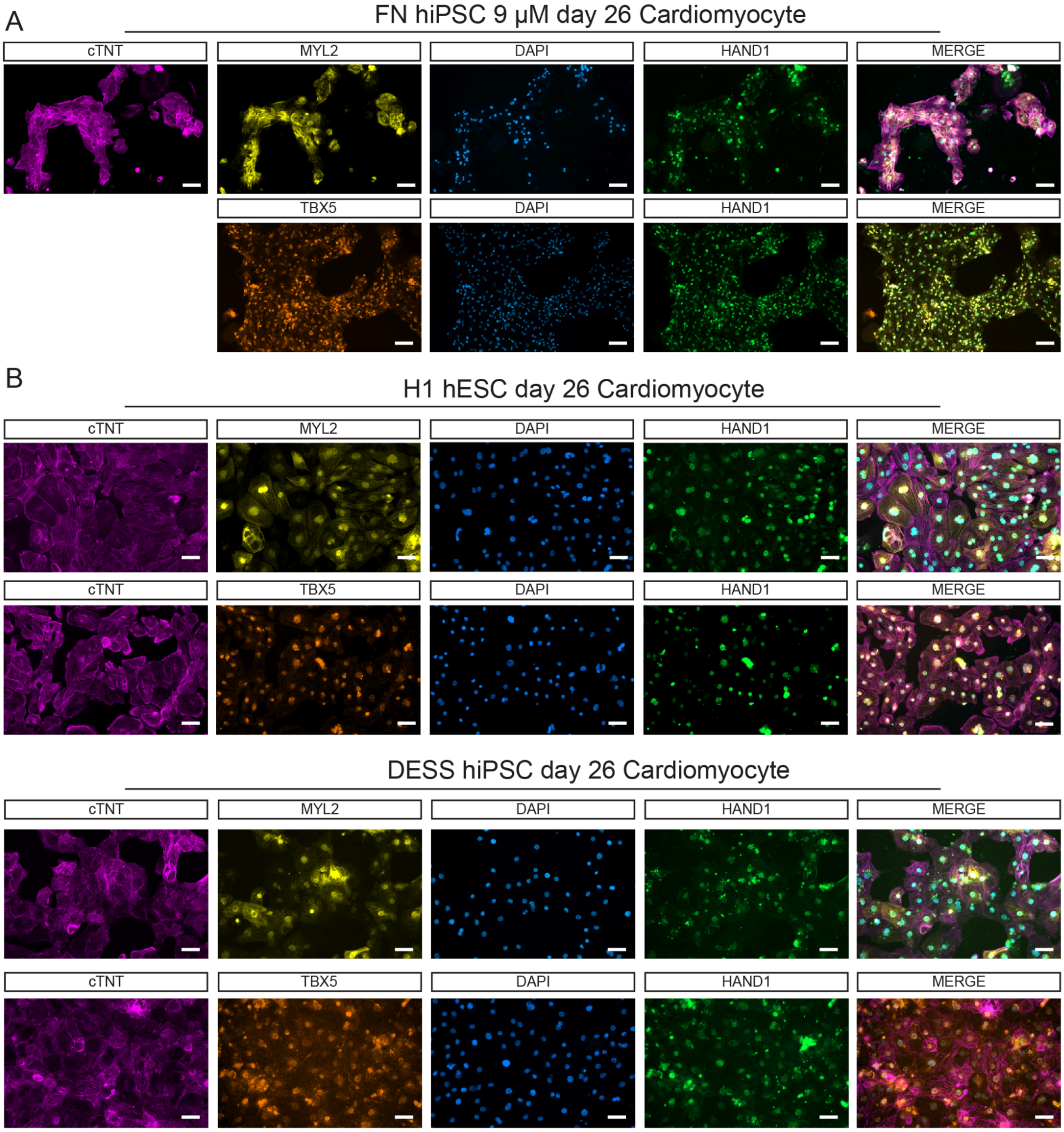


**Figure S3: Extended culture and lactate selection enhance MYL2 expression in cardiomyocytes in different stem cell lines.** A) Immunofluorescence staining of HAND1, MYL2, cTNT, and DAPI in pure cardiomyocytes at day 26 for the 9 μM CHIR condition, n=4. Scale bar = 100 μm. B) Immunofluorescence staining of HAND1, MYL2, cTNT, and DAPI in pure cardiomyocytes at day 26 for H1 hESC and DESS hiPSC cell lines differentiated using 12 μM CHIR, n=3. Scale bar = 100 μm.

Figure S4

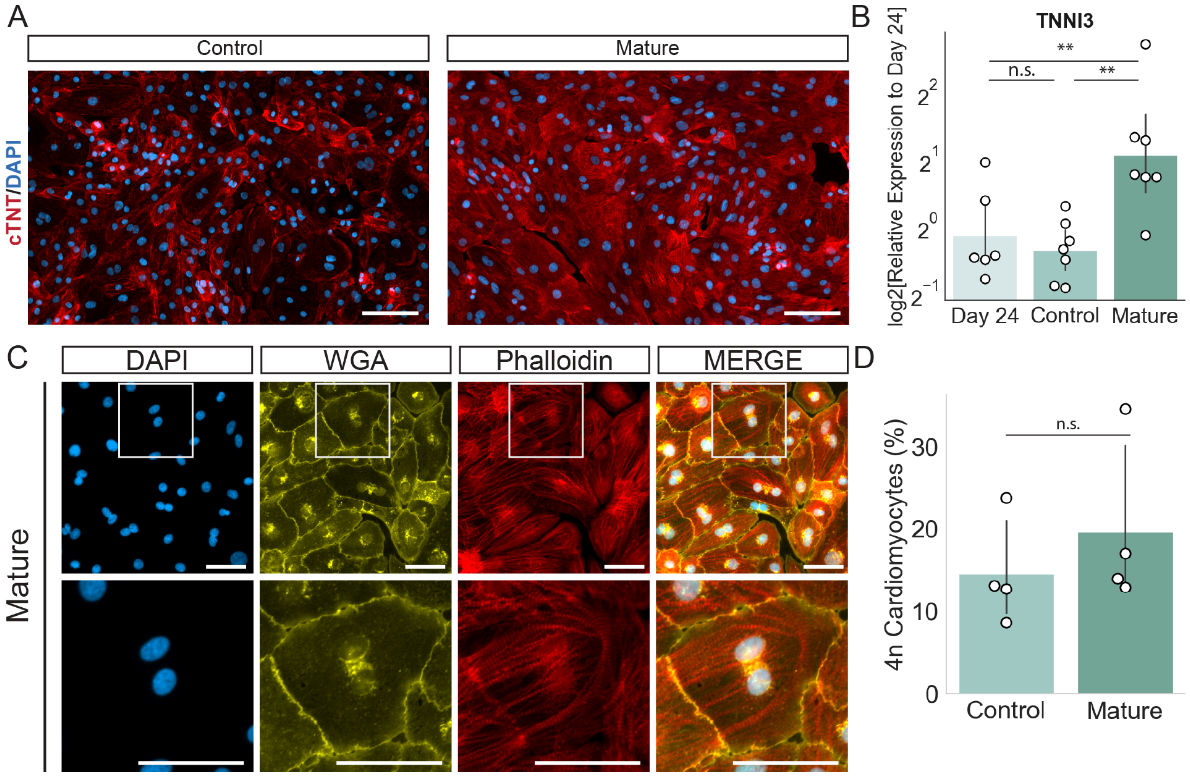


**Figure S4: Ploidy Quantification of mature cardiomyocytes.** A) Immunofluorescence staining of cTNT and DAPI in day 38 control and mature CMs. Scale bar = 100 μm. B) RT-qPCR of TNNI3 in day 24, control and mature CMs, n=6. C) Representative immunofluorescence of DAPI, WGA, and Phalloidin in mature CMs, Scale bar = 100 μm. D) Ploidy quantification using DAPI and immunofluorescence for control and mature CMs, n=4. Data are represented as mean and individual values *p < 0.05, **p < 0.01, ***p 0.001; n.s., not significant.

**Figure S5**

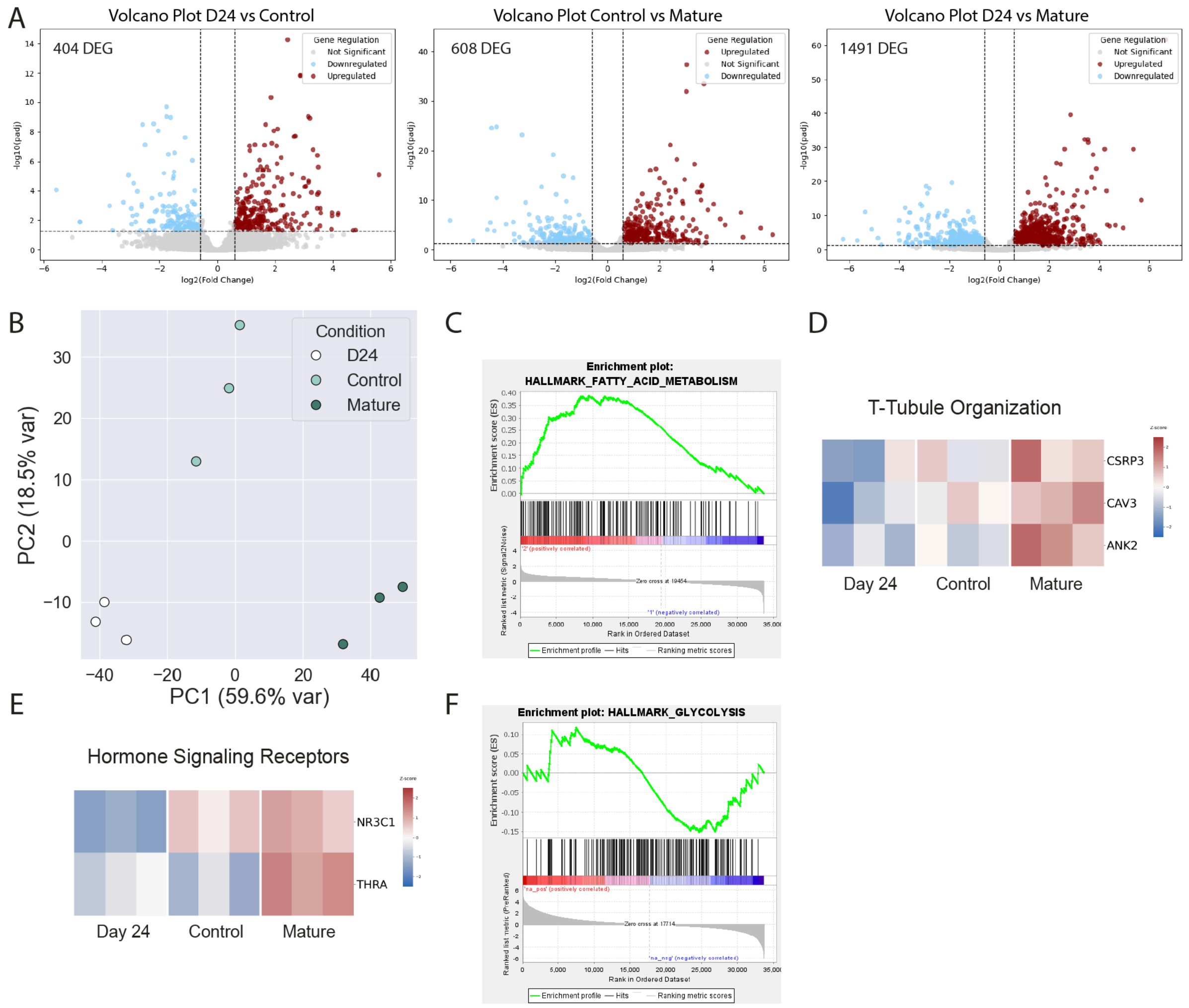


**Figure S5. Transcriptomic maturation signatures and pathway analysis.** A) Volcano plots of differentially expressed genes for the following comparisons (left to right): Day 24 vs. Control, Control vs. Mature, and Day 24 vs. Mature. B) Principal component analysis (PCA) of the 1,800 differentially expressed genes across all conditions. C) Gene Set Enrichment Analysis (GSEA) enrichment plot for the Fatty Acid Metabolism pathway in the mature condition. D) Heatmap of genes involved in T-tubule organization. E) Heatmap of hormone signaling receptor genes. F) GSEA enrichment plot for the Glycolysis pathway in the control condition.

**Figure S6**

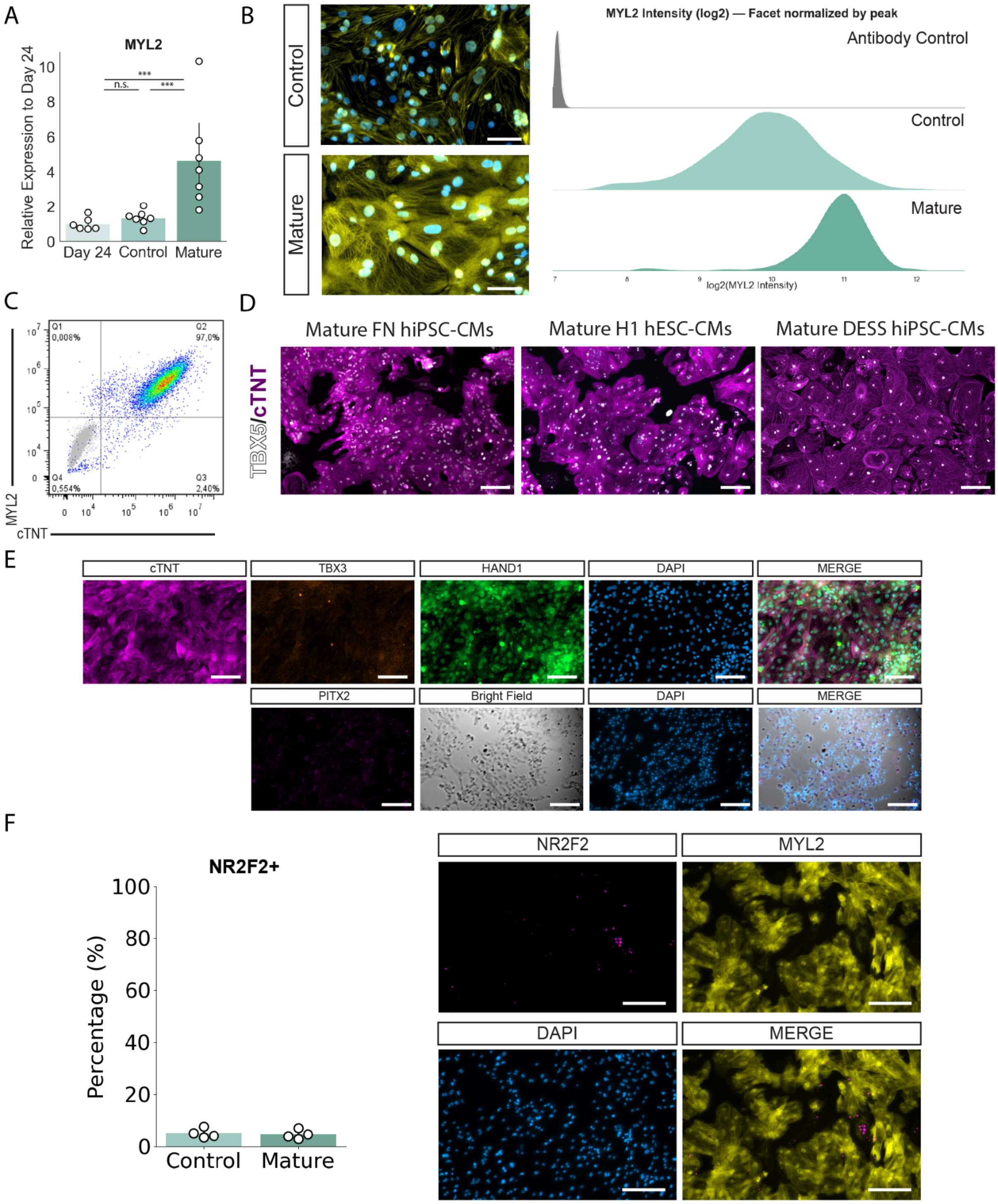


**Figure S6. Mature cardiomyocytes derived via WNT ON/OFF differentiation exhibit strong left ventricular identity.** A) RT-qPCR of MYL2 in day 24, control, and mature CMs, n=6. B) Image cytometry of MYL2 in control and mature CMs. Scale bar= 50 µm. C) Flow cytometry for MYL2+/cTNT+ in Mature cardiomyocytes. Gray: antibody control. D) Inmunofluorescence of TBX5 and cTNT on mature CMs derived from the three different PSC lines, n=3. Scale bar= 100 µm. E) Immunofluorescence staining of Atrial and SAN markers in mature cardiomyocytes for the FN hiPSC cell lines, n=2. F) Image cytometry of the atrial marker NR2F2 in mature CMs, n=1. Scale bar= 100 µm. Data are represented as mean and individual values *p < 0.05, **p < 0.01, ***p 0.001; n.s., not significant.

#### SUPPLEMENTAL VIDEOS

**Video S1:** day 14 cultures generated via 6 uM CHIR during the first 24 hours.

**Video S2:** day 14 cultures generated via 9 uM CHIR during the first 24 hours.

**Video S3:** day 14 cultures generated via 12 uM CHIR during the first 24 hours.

**Video S4:** day 38 CMs in Control conditions.

**Video S5:** day 38 CMs in Mature conditions.

#### SUPPLEMENTAL TABLES

**Table S1**: Gene Ontology (GO) analysis of upregulated genes in mature cardiomyocytes.

**Table S2**: Gene Ontology (GO) analysis of downregulated genes in mature cardiomyocytes.

**Table S3**: Primer sequences used for quantitative PCR (qPCR) analysis.

### SUPPLEMENTAL EXPERIMENTAL PROCEDURES

#### Quantitative Bibliographic Analysis of Cardiac Protocols

To assess the prevalence of different cardiac differentiation protocols, we performed a bibliographic analysis using automated approaches implemented in Python. We searched PubMed for cardiac differentiation studies from hPSCs, identifying approximately 4000 publications from 2011-2025. We then determined which publications were available through PubMed Central, which resulted in a list of 3282 publications. From the PMC-available subset, we first filtered out review articles, papers that did not perform cardiac differentiation from hPSCs, and papers that used commercially acquired hPSCs-CMs. The remaining ~1800 publications underwent automated protocol classification using GPT-5 API (OpenAI) with predefined classification rules to categorize articles by differentiation protocol type (WNT ON/OFF, embryoid body, other methodologies, etc). A sample of more than 50 papers was manually assessed to validate the classification efficiency. Protocol data was grouped by year and plotted in Python.

#### hPSC Culture

Human induced pluripotent stem cell (hiPSC) line FN2.1 (FN) was previously developed in Miriuka Lab from male foreskin fibroblasts (Questa et al., 2016) and registered in the cell line registry platform hPSCreg (https://hpscreg.eu/) under the name INEUi002-A. Female hiPSC line DESS was also previously developed in our lab from female donor PBMCs (Castañeda et al., 2023). H1 hESCs (Thomson et al., 1998) were commercially acquired from WiCell.

hPSCs were cultured from passages 10 to 25 and routinely grown on Geltrex™-coated (ref. A1413302, Thermo Fisher Scientific) dishes in StemFlex Medium (ref. A3349401, Thermo Fisher Scientific) without antibiotics. Cells were passed every 3-4 days to maintain a confluence under 60-70% TrypLE Select 1X (ref. A1217702, Thermo Fisher Scientific) following the manufacturer's instructions. Cell culture medium was supplemented with 10 µM Rho kinase inhibitor (ROCKi) Y-27632 (ref 1254, Tocris) for the first 24 h. Cultures were maintained in a humidified atmosphere at 37 °C with 5% $CO_2$ and were routinely tested for mycoplasma contamination using PCR.

#### Cardiac Differentiation Protocol

hPSCs were differentiated into cardiomyocytes in vitro as described by Lian et al (Lian et al., 2012). Cells were seeded at 200-300,000 cells/24 well on Geltrex™-coated plates in StemFlex Medium supplemented with 10 µM Y-27632 (ref. Cat#1254, Tocris) for 24 h. The medium was then replaced, and cells were cultured for an additional 48 h to form a confluent monolayer, which was designated as day 0. On day 0, the basal medium was replaced with RPMI 1640 (ref. 22400-089, Thermo Fisher) supplemented with B27 without insulin (ref. A1895601, Gibco), and cells were treated for 24 h with CHIR99021 (6 µM, 9 µM, or 12 µM; ref. 4423, Tocris). On day 3, cells were treated with 5 µM IWP2 (ref. 3533, Tocris) for 48 h. From then on, the medium was renewed every 48 h until self-contractile beating cells appeared, approximately at day 8. At this stage, the medium was switched to RPMI 1640 supplemented with B27 with insulin (ref. #17504044, Gibco). On day 12, cardiomyocytes were replated, and the medium was changed to cardiomyocyte enrichment medium. On day 14, cardiomyocytes were quantified by staining for cardiac troponin T (TNNT2, see Key Resources Table) and analyzed via flow cytometry.

#### Cardiomyocyte Dissociation Protocol

Day 12 cardiomyocytes were dissociated as follows: cells were washed twice with PBS and incubated for 8 minutes with Trypsin-EDTA 1X (ref. #25200-072, Gibco). Trypsin was inactivated using DMEM containing 10% FBS (Natocor). The cells were then centrifuged at 1300 RPM for 5 minutes and resuspended in replating medium, which consisted of DMEM supplemented with 20% FBS , 10 µM Y-27632, GlutaMAX (ref. #35050-061, Gibco), MEM NEA (ref. #11140-050, Gibco), and Pen/Strep (ref. #15140-122, Gibco). Cells were replated and incubated for 24 h.

Day 21 and day 38 cardiomyocytes (both matured and immature) were dissociated using Trypsin-EDTA 1X. The cells were washed twice with PBS, incubated with Trypsin/EDTA for 8 minutes, and inactivated using DMEM with 10% FBS. Centrifugation was performed at 1300 RPM for 5 minutes, and cells were replated using replating medium for 24 h.

##### Cardiomyocyte Enrichment Protocol

Day 12 cardiomyocytes were replated at a 1:5 ratio in replating medium. After 24 h, the medium was switched to cardiomyocyte enrichment medium, consisting of RPMI1640 without glucose (ref. 11879020, Thermo Fisher Scientific) supplemented with 7 mM sodium DL-lactate (ref. L4263, Sigma), 213 µg/ml L-ascorbic acid 2-phosphate (ref. A8960, Merck) and 0.5 µg/ml BSA fraction V (ref. 15260, Thermo Fisher Scientific) for 6 days (three medium changes with PBS washes in between). On day 19, the medium was replaced with RPMI 1640 supplemented with B27 with insulin for 48 h. On day 21, cells were replated for maturation.

#### Cardiomyocyte Maturation Protocol

After replating on day 21, day 24 cardiomyocytes were exposed to either control or maturation medium. Control medium: RPMI 1640 supplemented with B27 with insulin. Basal maturation medium (M-): DMEM containing 1 g/L glucose, 200 µM palmitic acid (ref. #P0500, Sigma), 500 µg/mL BSA fraction V, B27 without insulin, 50 µg/mL monothioglycerol (ref. #M6145, Sigma), 50 µg/mL L-ascorbic acid 2-phosphate, 150 µg/mL holo-transferrin (ref. #T0665, Sigma), and GlutaMAX. Full maturation medium (M+): M- supplemented with 200 ng/mL dexamethasone (ref. #D4902, Sigma), 4 nM T3 hormone (ref. #T6397, Sigma), and 1 µM GW7647 (PPAR-α agonist; ref. #G6793, Sigma).

Control condition cells were exposed to the control medium from day 24 to day 38. Mature condition cells were exposed to M+ for 9 days, followed by M- for 5 days. Media for both conditions was refreshed every 2 days, replacing 75% of the volume.

#### Flow Cytometry

After dissociation, cells were centrifuged and fixed for 10 minutes with 4% PFA at 4 °C. hiPSC-CMs were permeabilized and blocked for 30 minutes with permeabilization/block solution PBTS consisting of  1X PBS, 0.1% Triton, 3% normal donkey serum (ref. #D9663-10ML, Sigma). Primary antibodies were diluted in PBST and incubated for 1 hour at 4 °C. After three PBST washes, cells were incubated with 1:400 PBST-diluted secondary antibodies for 30 minutes at room temperature. Samples were washed three times with PBST and resuspended in 1X PBS. A total of 12500 events were analyzed using the C6 Accuri Cytometer. A list of the antibodies used is available in the Key Resource Table.

#### RT-qPCR

Total RNA was prepared with TRIzol Reagent (ref. #15596018, Ambion) following the manufacturer's instructions, treated with DNAse I (ref. #18047-019, Thermo Fisher Scientific), and then reverse transcribed into cDNA using MMLV reverse transcriptase (ref. #M1705, Promega) and random primers (ref. #48190011, Invitrogen). Quantitative real-time PCR (qPCR) was performed in a StepOne Real Time PCR system (Applied Biosystems) using FastStart Universal SYBR Green Master (Rox) (ref. #4913914001, Roche). Gene expression was quantified, normalized to the geometrical mean of GAPDH and RPL7 housekeeping genes for samples until day 14 and TBP and TPL7 for samples from day 14 to day 38. Primer sequences can be found in Table S3.

#### Immunofluorescence

For intracellular staining, cells were fixed with 4% PFA for 10 minutes at 4 °C and washed three times with PBS. Cells were blocked and permeabilized with PBST for 30 minutes. Primary antibodies were diluted in PBST according to the manufacturer's indications and incubated overnight at 4 °C. After three PBST washes, cells were incubated with the corresponding secondary antibodies diluted in PBST, along with DAPI, for 1 hour at room temperature. Images were obtained using the EVOS XL Core Imaging System (Thermo Fisher Scientific) and the Zeiss Axio Observer Z1 under fixed exposure and brightness conditions.

For WGA staining (ref. #W11261, Invitrogen), cells were incubated for 10 minutes at 37 °C in 1:200 HBSS WGA solution, washed three times with PBS, and fixed with 4% PFA.

For Mitotracker Green FM staining (ref. ##M7514, Invitrogen), cells were incubated for 10 minutes at 37 °C at 100 nM in a HBSS solution. Washed three times with PBS and observed in the microscope.

#### Bioimaging Pipeline

For image cytometry analysis, all images within each experiment were acquired using fixed exposure times and intensity settings across all conditions to ensure comparability. Images were analyzed using ImageJ and Python (v3.10). Nuclei were segmented using StarDist, while cells were segmented using Cellpose.

For experiments involving cardiac troponin T (cTNT), this channel was first used to identify and filter cTNT-positive cells (cardiomyocytes), and subsequent quantification of other fluorescence markers was performed exclusively within this cardiomyocyte population. Nuclei quantification was performed by calculating the average fluorescence intensity for each marker within each nucleus, after ensuring strict correlation between nuclei and cytoplasm fluorescence intensity.

Polynucleation was quantified by co-staining cells with DAPI (nuclei) and wheat germ agglutinin (WGA, cell membranes). Nuclei were segmented using StarDist, and cell membranes were segmented using Cellpose, generating separate label images for each structure. A custom Python algorithm was then used to count the number of nuclei per cell and calculate the nucleation status of each cardiomyocyte for each condition.

Ploidy was determined by leveraging the stoichiometric intercalation of DAPI into DNA. Following nuclear segmentation, the integrated fluorescence intensity of DAPI within each nucleus was quantified. Ploidy levels were determined by analyzing the distribution of intensity values, where populations falling at multiples of the baseline 2n intensity correspond to higher ploidy states (4n, 8n, etc.) (Yücel and Solinsky, 2021).

The mean order parameter for phalloidin staining was calculated using the AFT-Tool Python library (Marcotti et al., 2021).

Cardiomyocyte contractility was assessed by brightfield microscopy using a 10X objective. Videos of spontaneously beating cells were acquired for 20 s in multiple fields per well with a Nikon DS-Fi2 camera. Contraction dynamics, including beat rate and contraction amplitude, were quantified from the recordings using ContractionWave software (Scalzo et al., 2021).

##### Alternative Splicing Analysis

Alternative splicing events were identified and quantified using rMATS Turbo v4.3.0 (Shen et al., 2014). STAR-aligned BAM files from three biological replicates per condition (D23, Control, and Mature) were analyzed using the GENCODE v43 human annotation. rMATS was run with the following parameters: --readLength 198 --variable-read-length --allow-clipping --nthread 6, enabling detection of variable-length reads and soft-clipped alignments typical of RNA-seq data.

Five types of alternative splicing events were examined: skipped exon (SE), alternative 5' splice site (A5SS), alternative 3' splice site (A3SS), mutually exclusive exons (MXE), and retained intron (RI). Junction count (JC) files were used for downstream analysis, which quantify reads spanning splice junctions.

Pairwise comparisons were performed between D24 vs Control and D24 vs Maturation conditions to identify maturation-associated splicing changes. Percent Spliced In (PSI) values were extracted for each replicate, and differential splicing was assessed using rMATS statistical framework with likelihood-ratio tests. Events were considered significant at FDR < 0.05 with |ΔPSI| > 0.1.

For visualization, PSI values were organized into a matrix across all three conditions (9 samples total) and Z-score normalized per event. Hierarchical clustering was applied to the top 50 differentially spliced events using correlation distance and complete linkage. Gene-specific splicing patterns (TNNT2, MYOM1, FN1, and CMYA5) were visualized using scatterplots showing individual replicate PSI values with mean ± SE bars. Statistical significance across conditions was assessed using one-way ANOVA followed by Tukey's HSD post-hoc test. All visualizations were generated using R (v4) with ggplot2, pheatmap, and rstatix packages.

##### DNA Extraction

Cells were detached and centrifuged for 5 minutes at 300 g. The pellet was resuspended in lysis buffer I (5 mM PIPES, 85 mM KCl, 0.5% Igepal) and incubated on ice for 10 minutes. Nuclei were centrifuged at 400 g at 4 °C, then resuspended in lysis buffer II (50 mM Tris, pH 8; 1% SDS; 10 mM EDTA, pH 7.5) and incubated for 2 h on ice. DNA was extracted using UltraPure™ phenol:chloroform: isoamyl alcohol (25:24:1; Thermo Fisher Scientific) followed by ethanol precipitation with GlycoBlue™ (Thermo Fisher Scientific). The DNA pellet was washed with cold 70% ethanol, air-dried, and resuspended in nuclease-free water.

##### Statistical Analysis

All experimental data are presented as mean and individual values and 95% confidence interval, unless otherwise specified. Statistical analyses were performed using Python (v3.10) and SciPy library. Each experiment was independently repeated at least three times unless otherwise stated in the figure legend.

Comparisons between the two groups were performed using Student’s t-test. For experiments involving more than two groups, a one-way analysis of variance (ANOVA) followed by appropriate post hoc tests was used to determine statistical significance.

For time-course experiments evaluating maturation versus control conditions, a difference-in-differences (DiD) approach was applied to account for baseline differences and isolate the effect of the maturation intervention over time.

A p-value of < 0.05 was considered statistically significant. The exact p-values, sample sizes (n), and statistical tests used are indicated in the corresponding figure legends.
